# Genetic crosses within and between species of *Cryptosporidium*

**DOI:** 10.1101/2023.08.04.551960

**Authors:** Sebastian Shaw, Ian S. Cohn, Rodrigo P. Baptista, Guoqin Xia, Bruno Melillo, Fiifi Agyabeng-Dadzie, Jessica C. Kissinger, Boris Striepen

**Author notes:** Corresponding author Boris Striepen.

## Abstract

Parasites and their hosts are engaged in rapid coevolution that balances competing mechanisms of virulence, resistance, and evasion. This often leads to host specificity, but genomic reassortment between different strains can enable parasites to jump host barriers and conquer new niches. In the apicomplexan parasite *Cryptosporidium* genetic exchange has been hypothesized to play a prominent role in adaptation to humans. The sexual lifecycle of the parasite provides a potential mechanism for such exchange; however, the boundaries of *Cryptosporidium* sex are currently undefined. To explore this experimentally, we established a model for genetic crosses. Drug resistance was engineered using a mutated phenylalanyl tRNA synthetase gene and marking strains with this and the previously used Neo transgene enabled selection of recombinant progeny. This is highly efficient, and genomic recombination is evident and can be continuously monitored in real time by drug resistance, flow cytometry, and PCR mapping. Using this approach multiple loci can now be modified with ease. We demonstrate that essential genes can be ablated by crossing a Cre recombinase driver strain with floxed strains. We further find that genetic crosses are also feasible between species. Crossing *C. parvum,* a parasite of cattle and humans, and *C. tyzzeri* a mouse parasite resulted in progeny with a recombinant genome derived from both species that continues to vigorously replicate sexually. These experiments have important fundamental and translational implications for the evolution of *Cryptosporidium* and open the door to reverse- and forward-genetic analysis of parasite biology and host specificity.

**Significance statement:** The parasite *Cryptosporidium* is a leading cause of diarrheal disease. While infection is common all around the world, young children experiencing malnutrition are impacted most profoundly, and the disease is an important contributor to early childhood mortality. This study experimentally demonstrates that different strains and even species of *Cryptosporidium* can recombine their genomes through sex. The progeny of such genetic crosses shows combined features of both parents, with resistance to multiple drugs being one example. Sex thus provides a critical mechanism for the parasite to rapidly adapt to changing environments and hosts.

Genetic crosses as an experimental tool may also be harnessed in the future to discover the genes underlying differences in virulence, drug sensitivity, and immunogenicity between parasite isolates.

## Introduction

The gastrointestinal parasite *Cryptosporidium* is a leading cause of diarrheal disease around the world and responsible for frequent waterborne outbreaks in the United States [1,2]. *Cryptosporidium* is an AIDS defining opportunistic infection and a severe threat to immunocompromised individuals [3]. While the disease is typically self-limiting in healthy adults, more recent epidemiological studies revealed that young children, in particular those who are malnourished, are highly susceptible to severe disease, and cryptosporidiosis is an important cause of early childhood mortality [2] as well as growth and development stunting [4–6]. Vaccines and fully effective treatments are lacking.

Initially, a single species, *Cryptosporidium parvum,* was recognized in a wide range of mammals including humans [7]. Extensive population genetic analysis [8] led to the current model of numerous species, subspecies, and strains [9]. While these organisms have very similar genome sequences [10], they show profound phenotypic differences in host specificity [11], virulence [12], antigen repertoire [13,14] and drug susceptibility [15], which impacts the epidemiology [16,17], treatment, and prevention of the disease.

*Cryptosporidium* has a single host, obligate sexual lifecycle, and sex is hypothesized to drive parasite adaption to new hosts, and changing environments [17–19]. However, the true boundaries for sex and recombination within and between *Cryptosporidium* species are unknown, and our mechanistic understanding of gamete compatibility, fertilization, and meiosis is limited. Genetic crossing experiments have been attempted, but technical challenges in isolating and typing progeny limited the conclusions that could be drawn [20–22]. Here we sought to develop a rigorous model for genetic crosses in *Cryptosporidium* by using drugs to isolate recombinant progeny to the exclusion of parental organisms. However, thus far, only a single drug-selectable marker was available for this organism, neomycin phosphotransferase (neo) which confers resistance to paromomycin [23]. *Cryptosporidium* is naturally resistant to the inhibitors of protein, nucleotide, and folate synthesis used in other apicomplexans due to its highly reduced metabolism [23–25]. The necessity to propagate the parasite in animals excludes many resistance genes due to the toxicity of the drugs used in their selection. We therefore turned our attention to aminoacyl-tRNA synthetases, essential enzymes that have emerged as targets of anti-parasitic compounds [26–31]. Among these, phenylalanyl tRNA synthetase (PheRS) was recently identified as a promising antimalarial target by phenotypic screening followed by genetic and biochemical validation studies [27,32]. The bicyclic azetidine BRD7929, an analog of the screening hits, was highly potent in multiple models of malaria. *Cryptosporidium* is susceptible to BRD7929 in vitro and in vivo, and a single point mutation of the pheRS gene confers enhanced resistance [26]. Building on this finding here, we design, test, and validate a dominant drug selection marker. We demonstrate that using mutated pheRS and neo as markers, recombinant progeny is readily isolated in genetic crossing experiments, and use this model to construct complex mutants to explore the species boundaries of sex in *Cryptosporidium*.

## Results

### Mutation of *Cryptosporidium* pheRS permits selection for transgenic parasites

*Cryptosporidium* modified to carry a mutation changing PheRS leucine 482 to valine showed enhanced resistance to BRD7929 [26]. We hypothesized that this mutation could function as a dominant marker for transgenesis. We therefore engineered a template for homologous recombination to modify the endogenous pheRS locus that also introduces a nano luciferase/tdNeonGreen reporter cassette (Nluc/tdNG) (Fig. 1A). *C. parvum* sporozoites were electroporated with this construct along with a CRISPR/Cas9 plasmid targeting the pheRS locus (Fig. 1A). Ifnγ ^-/-^ mice infected with these parasites were treated with 10 mg/kg/day BRD7929. BRD7929 is not yet commercially available and we therefore synthesized 1g of the compound; please refer to the SI appendix and supplementary figures S7-9 for detail on optimized synthesis and documentation of structure and purity. Sustained luciferase activity was detectable in the feces of infected mice beginning on day 8 (Fig. 1B). Oocysts shed from day 9-17 were purified and subjected to flow cytometry, which showed green fluorescence in the transgenic parasites (pheRS^r^-Nluc-tdNG, Fig. 1C) but not in wildtype (*Cp* WT, Fig. 1C). Genomic DNA was isolated from transgenic and wildtype oocysts and PCR mapping of the pheRS locus confirmed transgene integration (SI Appendix, Fig. S1A). Human colorectal (HCT-8) cells were infected and showed green fluorescent parasites at all time points and lifecycle stages observed (SI Appendix, Fig. S1B). We have since used this strategy numerous times and found it to reliably result in the isolation of stable transgenic organisms.

**Figure 1.**
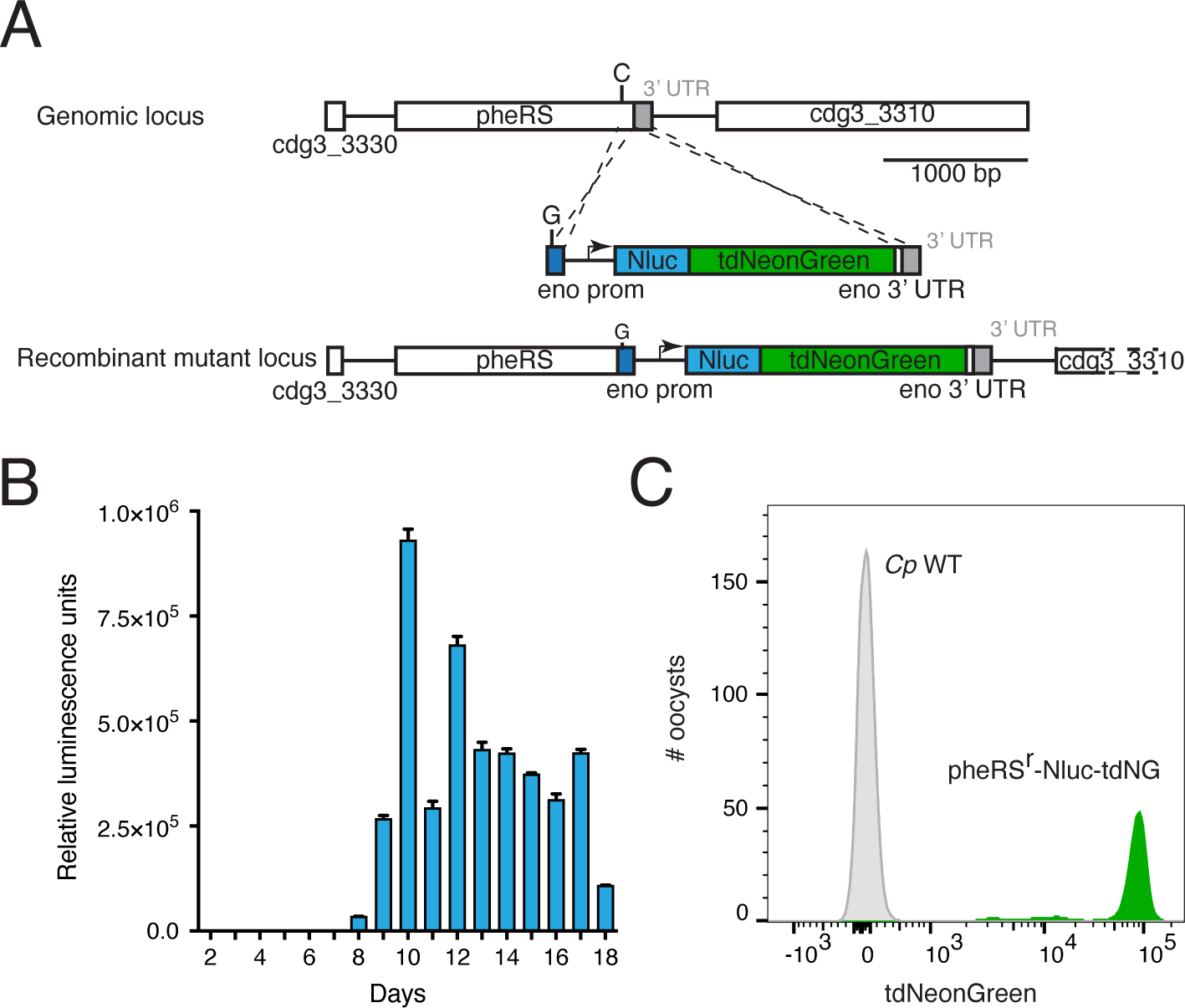
PheRS can be used as a selection marker for stable transgenesis. **A** Map of the pheRS locus targeted by insertion of a construct that includes the single base mutation that confers resistance to BRD7929, nano luciferase (Nluc), and the fluorescent protein reporter tdNeonGreen. **B** Ifnγ^-/-^ mice were infected with transfected parasites and burden was monitored by fecal luciferase activity. **C** Oocyst were purified from the feces and subjected to flow cytometry and transgenic parasites were highly fluorescent (grey: *Cp* WT; green: *Cp*-pheRS^r^-Nluc-tdNG).

### A mutated pheRS gene cassette can be used as a selection marker in trans

Mutating the pheRS locus is suitable to introduce transgenes but does not allow modification of other positions in the genome. We thus asked whether introducing a mutated copy of the pheRS gene into a different locus would allow selection in trans. We engineered a construct that consisted of the entire pheRS gene including its promoter and the resistance mutation (Fig 2A) and Nluc and targeted three dispensable loci that we previously successfully modified using the Neo marker. However, we only observed luciferase activity in one of these experiments targeting the thymidine kinase (*tk*) locus [23], and noted much lower luciferase reads than in previous experiments (Fig. 2B). We collected feces and infected a second group of mice, this time omitting BRD7929 treatment, and observed 100-fold higher luciferase activity in this second passage (SI Appendix, Fig S2A). PCR mapping showed integration of the mutated pheRS version into the TK locus (SI Appendix, Fig S2B) and amplicon sequencing demonstrated that only the transgenic copy carried the resistance mutation, while the endogenous version was unchanged (SI Appendix, Fig S2C). We conclude that pheRS can be used as a selection marker in trans to disrupt genes (pheRS^r^-TK-KO), but that the success rate appeared to be lower compared to modification of the endogenous locus.

**Figure 2.**
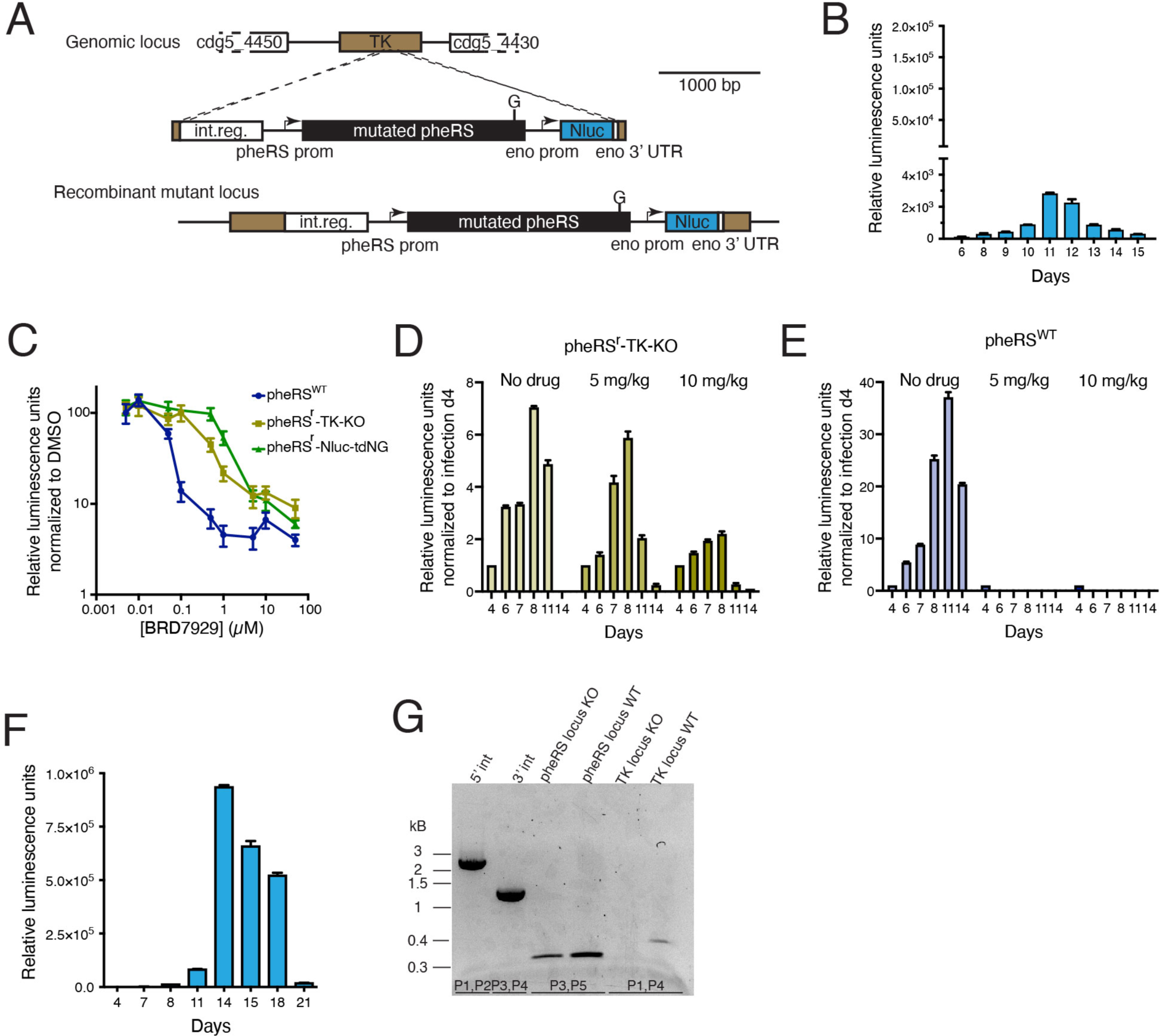
Using pheRS as selection marker *in trans*. **A** Map of the thymidine kinase (TK) locus targeted by insertion of a construct that includes the pheRS gene (the last 113 bp were recodonized and carry the resistance mutation) and nano luciferase (Nluc). **B** Fecal luciferase activity from mice infected with parasites transfected using the transgene shown in **A**. **C** Parasite growth in HCT-8 cells assessed by measuring luminescence in the presence of the indicated concentrations of BRD7929. Means and SD of 3 biological replicates are shown, the entire experiment was conducted twice with similar results. **D** and **E** Ifnγ^-/-^ mice were infected with pheRS^r^-TK-KO (**D**) or pheRS^WT^ (**E**), treated with the indicated doses of BRD7929 on days four to eight, and fecal nano luciferase activity was measured. **F** Parasites were transfected with construct shown in A Nano and infected Ifnγ^-/-^ mice were treated with selected with 5 mpk BRD7929 and fecal luciferase activity was measured. **G** PCR mapping using genomic DNA from transgenic parasites selected in **F** demonstrating integration into the TK locus.

We considered that pheRS modification in cis and trans might confer different levels of drug resistance. To test this, we compared the susceptibilities of the trans pheRS^r^-TK-KO (Fig. 2C, yellow), cis pheRS^r^-Nluc-tdNG (green), and pheRS^WT^ (blue) selected with paromomycin. Sporozoites were used to infect HCT-8 cultures, and luciferase activity was measured after 48h over a range of BRD7929 concentrations (Fig. 2C). The IC_50_ for pheRS WT was 56.5 nM and 1028 nM for pheRS^r^-Nluc-tdNG, respectively, similar to values previously reported for BRD7929 resistant and susceptible *C. parvum* [26]. Interestingly, for the pheRS^r^-TK-KO strain we determined an intermediate level of susceptibility (IC_50_=462.7 nM). We also measured susceptibility in animals. Ifnγ ^-/-^ mice were infected with the pheRS^r^-TK-KO strain and treated with 10mg/kg, the dose previously used, half that dose (5mg/kg) to roughly match the difference in IC_50_, or carrier. Fecal luminescence measurements were normalized to the values measured on day 4 when treatment was initiated (Fig. 2D). Mice treated with the half dose showed values similar to untreated mice (Fig. 2D), treatment with 10 mg/kg resulted in reduced shedding. Both doses were equally effective in curing mice infected with pheRS^WT^ (Fig. 2E). We conclude that introducing the pheRS^r^ marker in trans confers resistance, yet at a lower level. We therefore conducted transfection experiments employing BRD7929 dosing reduced to half and found that transgenic parasites were readily isolated with a parasite burden matching those seen when modifying pheRS in cis, or using Neo and paromomycin (Fig. 2F and G).

### Drug selection enables genetic cross and isolation of recombinant progeny in *C. parvum*

Genetic exchange by sex has been reported for *Cryptosporidium* [33], but isolating recombinant progeny has been difficult. We therefore tested whether parasites resistant to different drugs, can be crossed using a dual selection protocol. We constructed *C. parvum* strains with complementary drug markers and fluorescent reporters: one BRD7929 resistant and green fluorescent (*Cp*-pheRS^r^-Nluc-tdNG), the other paromomycin resistant and red (*Cp*-Paro^r^-tdTom) (Fig. 3A). Both strains carried luciferase reporter to track the infection. Ifnγ ^-/-^ mice were infected with each strain individually or with both, and treatment with paromomycin and BRD7929 was initiated after four days (Fig. 3B). Mice infected with individual strains were rapidly cured by this treatment (Fig. 3C and 3D), and infection did not relapse when the drug treatment ended on day 8. In contrast, mice co-infected with both strains showed continued heavy shedding (Fig. 3E). We conclude that parasites, carrying a single marker, are fully susceptible to the drug cocktail, and that coinfection produced double-drug resistance, suggesting successful recombination and cross.

**Figure 3.**
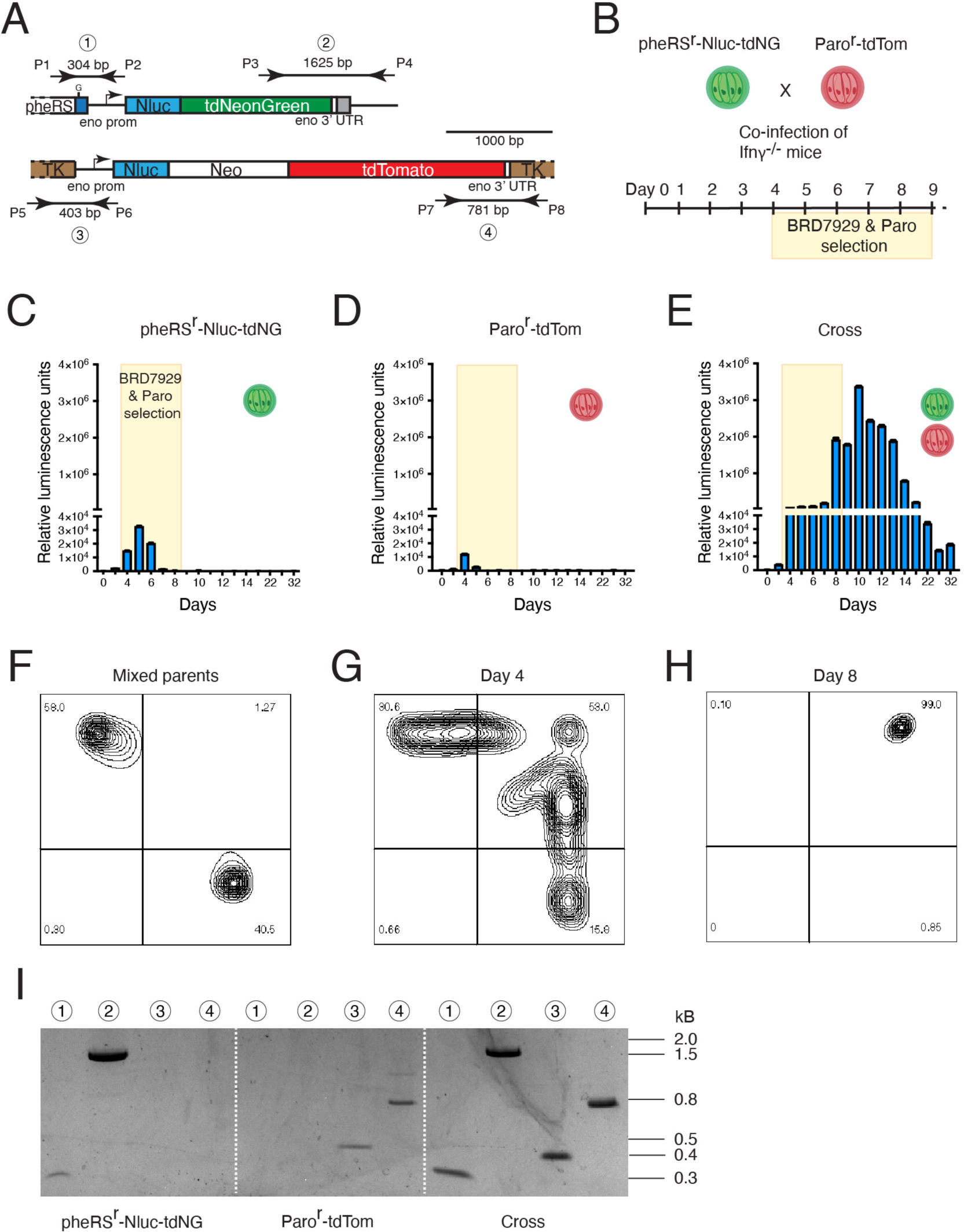
Establishment of highly efficient crosses in *Cryptosporidium*. **A** Maps of the loci used to mark each of the parental *C. parvum* lines. **B** Experimental set up of the crossing experiment between pheRS^r^-Nluc-tdNG which is susceptible to BRD7929 and expresses tdNeonGreen and Paro^r^-tdTom which is susceptible to paromomycin and expresses tdTomato. **C-E** Ifnγ^-/-^ mice were infected with the indicated parasite strains, subjected to dual drug treatment, and parasite burden was measured by fecal luciferase assay. Note that single infected mice are cured while co-infected mice remain infected. BRD7929 treatment was stopped on day 8, paromomycin treatment was continued until the end of the experiment. **F-H** Flow cytometric analysis of oocysts isolated from **E** At the days indicated (days 11 and 14 were indistinguishable from 8) **G** shows a mixture of both parental strains. **I** PCR mapping of the marked TK and pheRS loci in the parental lines and the cross (numbers indicate primer pairs highlighted in **A**). Note that the progeny of the cross is recombined and inherited markers and reporters from both parents.

We next used fluorescence to phenotype the progeny at the single-oocyst level. Oocysts were purified from the feces and subjected to flow cytometry on days 4 and 8. Parental parasites were used to establish strain specific cytometry gates, and we found exclusive staining of each strain (Fig. 3F). On day four of co-infection, we observed red, green, and importantly, double positive oocysts (Fig. 3G), which accounted for roughly half of the recorded events. Following the impact of treatment, we observed uniform red and green fluorescence (Fig. 3H). We note populations displaying intermediate fluorescence for either red or green on day four that are subsequently lost during selection (see discussion). When we PCR genotype oocysts using transgene specific primer pairs on the parental strains, single amplicons were detected in a strain-specific fashion, however, when using oocyst selected in crosses both amplicons were present (Fig. 3I). We repeated this genetic cross with similar outcome and thus conclude that applying the pressure of two drugs results in the rapid selection of recombinant progeny to the exclusion of the parental strains.

### Sexual cross permits isolation of conditionally lethal mutants

The ease and efficiency of recombination led us to ask whether crosses may enable more complex genetic manipulations in *C. parvum.* We recently demonstrated the use of rapamycin inducible Cre recombinase (DiCre) in conditional gene ablation, however, introducing loxP sites and recombinase in a single transfection is challenging (see extended technical discussion in [34]). To overcome this, we devised a model in which we cross a Cre driver and a floxed strain (Fig. 4A). We generated DiCre-expressing parasites using BRD7929 selection (pheRS^r^-Nluc-DiCre, Fig. 4A and SI Appendix, Fig. S3A) and, using paramomycin a color switch reporter (loxP-tdTom-loxP-tdNeon-Nluc-Paro^r^, Fig. 4A and SI Appendix, Fig. S3A). Introns harboring loxP sites were placed to allow Cre mediated excision of the red marker to activate the green. The strains were crossed as described (Fig. 3A), and purified oocysts were used to infect HCT-8 cells. After three days genomic DNA was isolated, and the floxed gene was amplified by PCR. Rapamycin treatment led to loss of the large amplicon and concomitant accumulation of a smaller molecule (Fig. 4B) consistent with Cre-mediated excision. We also noted some leaky Cre activity in the absence of rapamycin as previously described [34]. Infected cultures grown in the presence or absence of rapamycin for 48h, were fixed and imaged. Parasites not exposed to rapamycin expressed tdTomato, while those exposed to rapamycin expressed tdNeonGreen (Fig. 4C) and we quantified this switch using flow cytometry (Fig. 4D).

**Figure 4.**
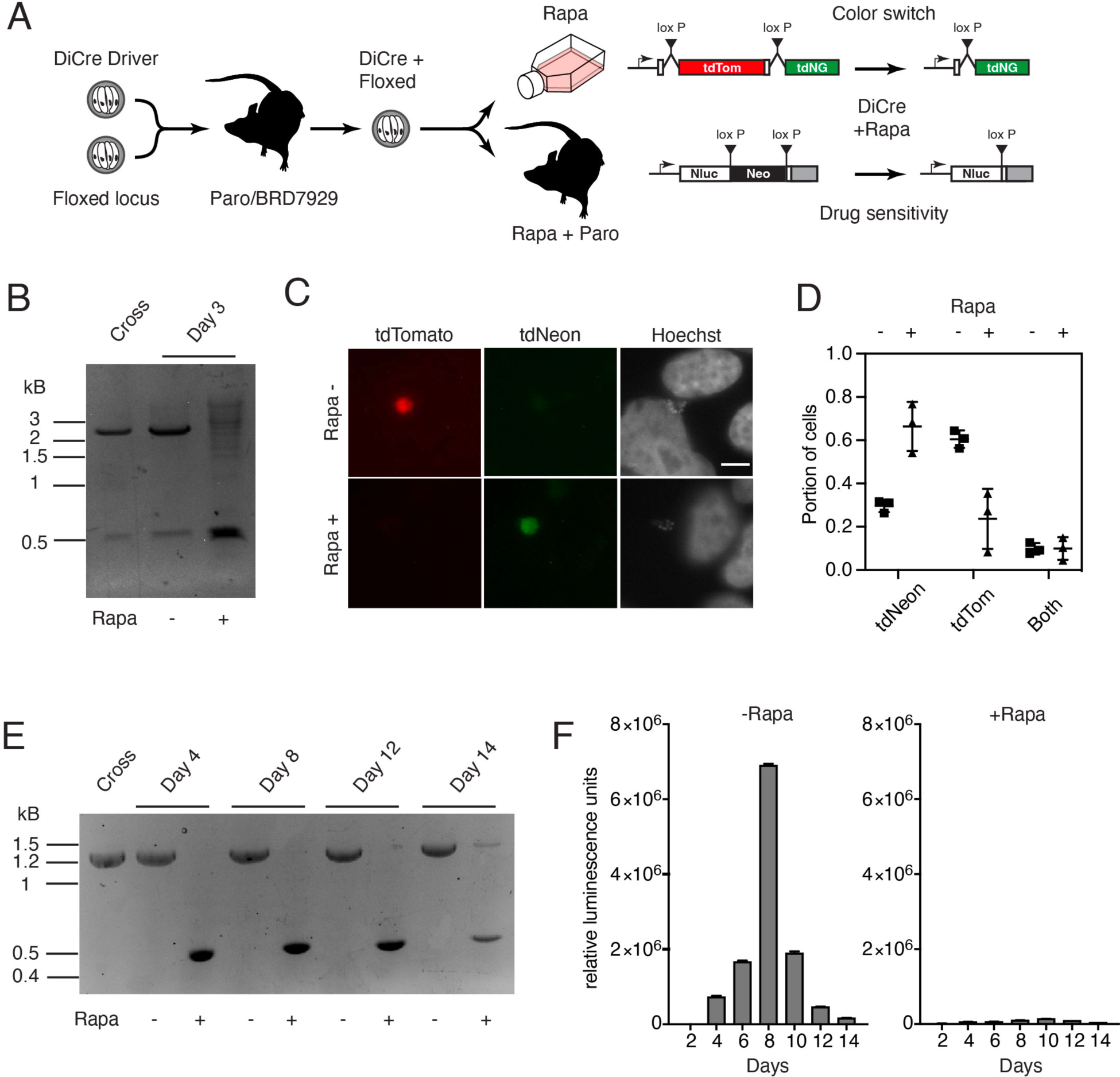
Conditional gene ablation in crossed progeny. **A** Illustration of the experimental set up crossing a Cre driver with floxed strains and the respective phenotypic assays for gene excision. **B-D** Progeny of the loxP-tdTom-loxP-tdNeon-Nluc-Paro^r^ color switch cross was used to infect HCT-8 cells and cultures were treated with rapamycin (rapa). **B** Genomic DNA was extracted from oocysts (cross) or from cultures infected for three days, and the floxed gene was amplified by PCR. Amplicons are 2076 bp for the floxed, and 467 bp for the excised locus, respectively. **C** Representative micrographs of HCT-8 cells infected with cross progeny after 48h with or without rapamycin. Scale bar, 5 μm. **D** HCT-8 cells infected with cross progeny were analyzed by flow cytometry after 72 h of rapamycin treatment and relative transgene expression is plotted as their portions of total fluorescent cells. Means and SDs of three biological replicates are shown. **E** and **F** Ifnγ^-/-^ mice were infected with progeny of the cross between the DiCre and *Cp*-loxP-Paro^r^-loxP strains. **E** Ifnγ^-/-^ mice infected with the progeny of a cross between the DiCre and *Cp*-loxP-Paro^r^-loxP lines, treated with rapamycin or vehicle and genomic DNA was extracted from the feces at the indicated times. PCR analysis revealed amplicons for the floxed (1269 bp) and excised locus (432bp), respectively. Note strong rapamycin dependent induction. **F** Fecal nano luciferase measurements from these mice revealed rapamycin induced drug sensitive as a consequence of in vivo Cre-mediated excision.

We next engineered a strain that carried a floxed drug resistance cassette (*Cp*-Nluc-loxP-Paro^r^-loxP, Fig. 4A and SI Appendix, Fig. S3A and B). This strain was again crossed with the DiCre driver and recombinant progeny was isolated. HCT-8 cells infected with these parasites were incubated in the presence or absence of rapamycin, and diagnostic PCR showed progressive excision under rapamycin (SI Appendix, Fig S3C). To test the functional consequence of treatment we turned to in vivo experiments. Two groups of Ifnγ ^-/-^ mice were infected with the progeny, and all animals were treated with paromomycin. One group was also treated with rapamycin (10mg/kg/day by gavage). PCR of the floxed locus demonstrated efficient rapamycin dependent excision in vivo (Fig 4E). Following fecal luminescence, we observed robust infection in the vehicle treated group, confirming the progeny’s resistance to paromomycin. In contrast, mice receiving rapamycin showed dramatic reduction of parasite burden (Fig 4F). We conclude that sexual cross permits modification of multiple independent loci, and that applying this system to gene deletion results in conditional mutants allowing the study of essential genes in vivo.

### Testing the boundaries of genetic exchange between *Cryptosporidium* species

Numerous species of *Cryptosporidium* have been described largely defined by ecological isolation through host specificity [19], reflected in names like *C. hominis, C. meleagridis,* or *C. canis*. Whether they are reproductively isolated, is unknown, but of great epidemiological importance [23,33,35]. We sought to test the species barrier experimentally using *C. parvum* and *C. tyzzeri,* two species with 95% identical genome sequence [36–38]. *C. tyzzeri* naturally infects mice, and *C. parvum* cattle and humans, however, both can infect mice lacking interferon γ. We therefore engineered *C. tyzzeri* strain STL ([39], a kind gift from Dr. Chyi-Song Hsieh) to express red fluorescent protein using paramomycin selection (optimized dose of 32 g/L*, Ct* Paro^r^-Nluc-tdTom, SI Appendix, Fig, S4A). PCR demonstrated transgenesis (SI Appendix, Fig. S4B), and purified oocysts showed homogenous red fluorescence (SI Appendix, Fig. S4C).

We next co-infected Ifnγ^-/-^ mice with 2000 oocysts of *Ct* Paro^r^-Nluc-tdTom and 8000 oocysts of *Cp* pheRS^r^-Nluc-tdNG and initiated dual drug selection on day 4. Parasite burden rapidly declined upon treatment (Fig. 5A), however, rebounded and increased to 100-fold by day 21, indicating the emergence of dual drug resistance. When analyzed by flow cytometry the parents were either red or green (Fig. 5B). In contrast, oocyst isolated on day 21 of coinfection were exclusively double positive (Fig. 5C). We amplified segments of the progeny’s genome carrying the resistance markers as well as the 18S and *gp60* loci, both commonly used in genotyping [40,41]. Sequencing these amplicons revealed *C. tyzzeri* specific SNPs in TK and *gp60* and *C. parvum* specific sequences in the PheRS and 18S loci (Fig. 5D). To test this genome-wide, we conducted whole genome sequencing of both parents and the progeny and obtained ∼75 fold coverage for the progeny (Fig. 5E). The *Cp* parent is near identical to the *C. parvum* reference genome (868 SNPs), while the *Ct* parent shows 186,218 SNPs. In Fig. 5F, the relative frequency of these *Ct* SNPs across all eight chromosomes of the progeny are plotted. This showed a high degree of interspecies recombination for multiple chromosomes. Some chromosomes are largely derived from a single parent, and this is particularly noticeable (but not exclusive) for those chromosomes carrying the selection marker (3 for *Cp* and 5 for *Ct*). We also obtained long-read sequence for smaller portions of the genome from single oocysts isolated by flow cytometry; it is currently impractical to clone this organism. Again, we find sequences from both parents, and bridging cross over points were supported by hundreds of long reads (Fig. 5G and SI Appendix, Fig. S5 show an example from chromosome 8).

**Figure 5.**
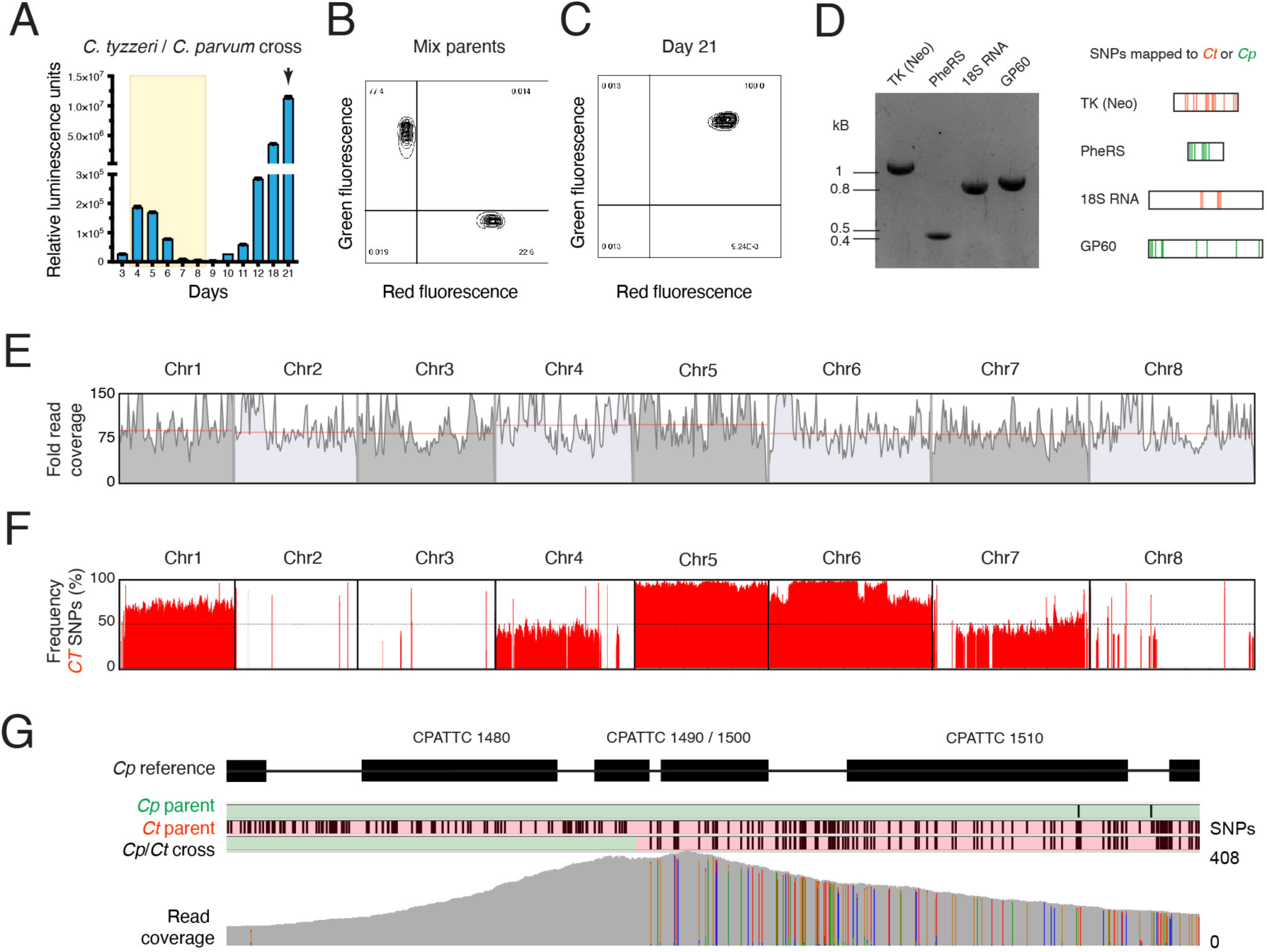
Testing the species boundaries of genetic exchange in *Cryptosporidium*. **A** Ifnγ^-/-^ mice were infected with *C. parvum* pheRS^r^-Nluc-tdNG and *C. tyzzeri* Paro^r^-tdTom and subjected to dual drug treatment over the time indicated by shading in yellow. Parasite burden was measured by fecal nano luciferase assay. **B** Flow cytometry of a mixture of both parental oocysts, and **C,** oocyst isolated on day 21 (arrow in **A**) following coinfection showing uniform double positive fluorescence. **D** The indicated loci were amplified by PCR from day 21 progeny oocysts and amplicons were subjected to Sanger sequencing. Amplicons were genotyped using specie specific SNPs. *C. tyzzeri* SNPS are shown in red, those specific for *C. parvum* in green. Agarose gel shows the fragments that were sent in for sequencing. Red and green lines depict SNPs between *C. parvum* and C. tyzzeri. **E** Coverage of nanopore reads of the cross. Red line indicates median coverage for each chromosome. **F** Calculation of the common *C. tyzzeri* SNP position and frequency obtained from bulk sequencing of the crossed progeny plotted across all 8 chromosomes. **G** Progeny was also subjected to single-oocyst sequencing and a recombinant locus on chromosome 8 is depicted (see SI Appendix, Fig. S5 for a multiple sequence alignment of long reads at this locus) and Supplemental data alignment of 50 randomly selected reads.

## Discussion

Parasitism is one of the most powerful ecological drivers of speciation and parasites like braconid wasps, mites, or apicomplexans are among the most species-rich eukaryotic taxa [42,43]. The host-parasite arms race of restriction and evasion results in rapid diversifying evolution, and favors the emergence of host specificity [44]. In turn, host specificity provides isolation by habitat, further accelerating speciation. This is evident in the genus *Cryptosporidium* where numerous different species and strains infect a variety of animals [45]. *Cryptosporidium parvum*, one of the most important and best studied species, alone comprises of at least 20 subtypes [46], some appear in the process of speciation by adaptation to humans as exclusive host [47]. Species boundaries are also maintained by reproductive isolation. What role such sexual incompatibility plays for parasites is largely unknown, but is an important question, as sexual recombination can result in hypervirulence and rapid global expansion of new strains as documented for e.g. *T. gondii* [48].

Here we develop an experimental model for genetic crosses in *Cryptosporidium parvum* in mice to study parasite sex. Using fluorescent parents, we show evidence of robust recombination by flow cytometry. Interestingly, early in infection we observed oocysts with different fluorescence levels, which likely reflects maternal cytoplasmic inheritance (see SI Appendix, Fig. S6). The female gamete is much larger, and responsible for the bulk of the oocyst proteome [35,49]. In the first cross, maternal fluorescence thus outweighs the male, following selection both parents are double positive, yielding uniform highly fluorescent progeny. Highly fluorescent oocysts therefore are the product of at least two subsequent crosses, indicating that four days into the infection, some parasites already ran through their lifecycle twice. This may appear fast but matches the compact 48h lifecycle recently established by time lapse microscopy [50]. Using two mechanisms of drug resistance selects for recombinant progeny to the exclusion of their parents. This is very efficient, underscoring that *Cryptosporidium* oocysts are indeed highly auto infective and continuously reset the lifecycle sustaining infection. This differs from other well studied apicomplexans where infection is sustained by the continuous replication of an asexual form and sexual differentiation is linked to transmission to the next host [51].

Progeny is readily observed when crossing *C. parvum* strains, and maybe surprisingly, also when pairing *C. parvum* with a different species, *C. tyzzeri.* We rigorously documented genetic exchange using measurements of drug resistance, fluorescent protein expression, PCR mapping, and whole genome sequencing, which demonstrated the presence of SNPs from both species on the same DNA molecule. We conclude that, using selection and a host susceptible to both species (we used Ifnγ^-/-^ mice here [38,52]) the species barrier can be breached. Not all chromosomes showed equal levels of recombination and further studies are needed to determine whether this reflects strong selection (e.g. due to the presence of the marker) or local meiotic incompatibilities.

The availability of a second selection marker, and the ease of crossing now permits modification or ablation of multiple loci. Crossing floxed parasites with Cre driver strains, resulted in highly penetrant conditional gene ablation, a critical tool to dissect essential phenomena in a haploid organism. Most importantly, the model reported here enables forward genetic studies in *Cryptosporidium*. Genetic mapping revealed mechanisms of drug resistance and virulence in *Plasmodium* [53,54] and *Toxoplasma* [55–58]. *Cryptosporidium* is well suited to this approach: it has a small genome, follows mendelian rules of inheritance [33,59,60], and strains that differ in host specificity, virulence, and drug susceptibility have been reported [12,15,62]. Much of the biology of *Cryptosporidium* remains poorly understood, we expect the use of genetic crosses to discover gene function by mapping to be transformative.

### Material and Methods Parasites and mice

*C. parvum* Iowa II (IIa) strain oocysts were obtained from Bunch Grass Farms. A *C. tyzzeri* strain isolated from laboratory mice at Washington University in St. Louis was kindly provided by Dr. Chyi-Song Hsieh, University of Washington [39]. Parasites were maintained in 6-8 week old male and female Ifnγ^-/-^ (Jackson Laboratory stock no. 002287) or C57BL/6 (Ct) mice bred in-house. 6-8 week old mice were handled, treated with antibiotics, and infected as detailed in [23].

### Generation of transgenic parasites

Guide oligonucleotides (Sigma-Aldrich) were introduced into the *C. parvum* Cas9/U6 plasmid [23] by restriction cloning (see [62] for guide design) and repair templates were constructed by Gibson assembly (New England Biolabs). Excysted sporozoites were transfected as described [62]. Detail on the transgenic strains generated in this study can be found in SI appendix. To generate transgenic parasites, 5 × 10^7^ oocysts were incubated at 37 °C for 1 hour in 10 mM HCl followed by two PBS washes and an incubation at 37 °C for 1 hour in 0.2mM sodium taurocholate and 22 mM sodium bicarbonate to induce excystation [63]. Excysted sporozoites were electroporated and used to infect mice as [62]. Integration was evaluated by PCR using primers detailed in SI appendix, Table S2.

### Cell culture and microscopy

HCT-8 cells were purchased from ATCC (CCL-224TM) and maintained in RPMI 1640 medium (Sigma-Aldrich) supplemented with 10% Cosmic calf serum (HyClone). Cells were infected with bleached and washed oocysts and serum was reduced to 1%. Infected cells were fixed with 4% paraformaldehyde (Electron Microscopy Science) in PBS and stained with Hoechst (1 μg/ml). Coverslips were then mounted on glass slides with fluorogel (Electron Microscopy Science) mounting medium.

### *Cryptosporidium* in vitro drug assays and IC_50_ determination

HCT-8 cells were infected with 1000 oocysts per well and incubated with drug. Medium was aspirated after 48 h, cells were lysed and mixed with NanoGlo substrate (Promega) and luminescence was measured using a Glomax reader (Promega) [62]. IC_50_ values were calculated in GraphPad Prism software v9 (two experiments, each conducted with triplicate wells).

### Rapamycin treatment

Rapamycin (LC Laboratories) was dissolved in 95% ethanol (50 mg/mL stock). Mice were weighed and treated daily by oral gavage with 100 μL of drug solution adjusted with water to deliver 10 mg/kg body weight.

### Flow cytometry

Oocysts were enriched using a miniaturized sucrose floatation of feces from infected mice (see SI appendix for a detailed protocol), resuspended in FACS buffer (1x PBS, 0.2% bovine serum albumin, 1 mM EDTA). Infected HCT8 cells were trypsinized, washed in FACS buffer, blocked (99.5% FACS Buffer, 0.5% normal rat IgG, 1 µg/ml 2.4G2), washed, and stained with Hoechst (1 μg/ml) on ice. All samples were passed through a 35 μm filter prior to flow cytometry using a LSRFortessa or FACSymphony A3 Lite and analyzed with FlowJo v10 software (TreeStar).

### Genome sequencing

DNA extraction, sequencing, and computational analyses including variant mapping are detailed in the SI Appendix. In brief, 150 bp paired-end Illumina sequencing was used for parental strains, cross progeny was subjected to multiple displacement amplification and Oxford Nanopore sequencing.

### Data Availability

Raw sequence read data can be obtained from NCBI BioProject PRJNA1000584 and BioSamples SAMN36772387, SAMN36772388, SAMN36772389.

## Acknowledgements

This work was supported in part by the National Institutes of Health (NIH) with grants to BS (R01AI112427 and R01AI127798), JK and Travis Glenn (R01AI148667), a fellowship to IC (F30AI169744), support to FAD (T32GM142623), and fellowships from the Swiss National Science Foundation to SS (P2BEP3_191774 and P500PB_211097). We thank Chyi Hsieh for sharing *C. tyzzeri* oocysts, and Abhijit Kundu (TCGLS) for small molecule synthesis, and Travis Glenn for help with single oocyst sequencing and mentorship of FAD.

