## Supplementary material for "Genetic crosses within and between species of *Cryptosporidium*": Shaw_Supplementary Information.pdf

### Supporting Information for
### Genetic crosses within and between species of *Cryptosporidium*

*Sebastian Shaw[1], Ian S. Cohn[1], Rodrigo P. Baptista[2], Guoqin Xia[3], Bruno Melillo[3,4], Fiifi Agyabeng-Dadzie[5], Jessica C. Kissinger[5,6], and Boris Striepen[1]**

*Corresponding author Boris Striepen.

**Supplementary Information Text**

#### Material and Methods

##### Detailed description of generation of transgenic parasites

*C. parvum* transgenics

Strain pheRS$^r$-Nluc-tdNG: The repair encodes the last 113 bp of the pheRS gene (cgd3_3320, recodonized) including the mutation that confers resistance (L482V) to BRD7929. This short sequence is followed by a nanoluciferase reporter fused to tdNeonGreen under the enolase promoter.

Guide: PheSF_guide_New_SV / PheSR_guide_New_SV

Repair template: ohgF_PheS_N_wmpam / ohg_PheR

Strain pheRS$^r$-TK-KO: The repair encodes for intergenic region upstream of the pheRS gene, the pheRS gene itself including the mutation that confers resistance (L482V) to BRD7929, followed by the enolase promoter and the neomycin phosphotransferase drug-selection marker fused to a nanoluciferase reporter.

Guide: TK_int_fwd / TK_int_rev

Repair template: pheRSKO fwd / pheRSKO rev

pheRS$^r$-Nluc-DiCre: The repair encodes the last 113 bp of the pheRS gene (cgd3_3320, recodonized) including the mutation that confers resistance (L482V) to BRD7929. This short sequence is followed by a nanoluciferase reporter under the enolase promoter. The Dicre is under the alpha tubulin promoter downstream of the nanoluciferase reporter.

Guide: PheSF_guide_New_SV / PheSR_guide_New_SV

Repair template: ohgF_PheS_N_wmpam / ohg_PheR

loxP-tdTom-loxP-tdNeon-Nluc-Paro$^r$: The repair encodes tdTomato (without a start codon), flanked by introns that contain loxP sites followed by a tdNeonGreen (without a start codon), Nluc and the neomycin phosphotransferase gene. There is a start codon in front of the first intron. The fluorescent reporters are riven by the alpha tubulin promoter und Nluc and the Neo cassette are under the enolase promoter.

Guide: TK_int_fwd / TK_int_rev

Repair template: loxP_repair_F / pheRSKO rev

Nluc-loxP-Paro$^r$-loxP: The repair encodes Nluc followed by the neomycin phosphotransferase drug-selection marker. The neomycin phosphotransferase gene is flanked by loxP sites.

Guide: TK_int_fwd / TK_int_rev

Repair template: loxP_repair_F / pheRSKO rev

*C. tyzzeri* transgenic

*Ct* Paro$^r$-Nluc-tdTom: The repair encodes Nluc/Neo fused to tdTomato with a T2A sequence.

The whole construct is running under the enolase promoter.

Guide: TK_int_fwd / TK_int_rev

Repair template: AG160 / Pr8522

**Mini-sucrose gradient and immune staining of infected fecal material**

Fecal material was collected, and 20 mg were homogenized in 1 ml cold tap water by pipetting. Slurry was passed through a cell strainer (40 µm Nylon, Falcon, Ref: 352340) and flushed once with 1ml cold tap water. Filtered slurry was collected in a 2 ml Eppendorf tube and pelleted at 3000 g for 5 min. Supernatant was aspirated and pellet resuspended in 1 ml cold tap water. 1 ml cold sucrose solution (1.33 specific gravity sucrose solution) was added and mixed well by inverting the tube 5 times. The solution was span for 5 min at 1000 g and the supernatant was divided into two new 2 ml tubes. 1 ml cold tap water was added to each tube and mixed well by inverting the tubes 5 times. Suspension was pelleted at 10'000 g for 5 min and supernatant aspirated. Pellets were resuspended and collected in 1 ml cold tap water. The cells were washed once in tap water followed by a wash in 1 ml cold FACS buffer (1x PBS, 0.2% BSA, 1mM EDTA). Cells were pelleted again, resuspended in 50 µl Crypt-o-Glo-biotin (A400BIOY-R-20X, Waterborne, Inc., New Orleans, LA) and incubated for 30 min at 4 °C. After the incubation, the cells were washed twice in FACS buffer and then incubated with 100 µl Streptavidin APC (1:1000, Invitrogen, Ref: 17-4317-82) for 30 min. Cells were washed twice in FACS buffer, pellet resuspended in 500 µl FACS buffer and stored at 4 °C until flow analysis.

**Illumina sequencing of the parental strains/species**

The DNA extraction from the oocysts involved a 5x freeze-thaw cycle method, which included subjecting the oocysts to 25-second freezing in liquid nitrogen, followed by a 5-minute thaw at room temperature, and a subsequent 30-second vortex step to break the oocysts. The DNA extraction was then carried out using the phenol-chloroform DNA extraction protocol. The library preparation from the oocysts for Illumina sequencing was carried out using the Illumina DNA Prep (former Nextera DNA Flex kit, Illumina Inc). Subsequently, sequencing was performed on the Illumina NextSeq 2000 sequencer, utilizing the P2 300 cycle flowcell kit to generate approximately 2-2.5 million 150-base pair paired-end reads per sample. The reads were trimmed with trimmomatic v0.39 [1], employing a Q-scores threshold of >25 and to remove adapters. The SPADES assembler v3.15.5 [2] was used for whole genome assembly.

**DNA preparation from the *C. parvum* x *C. tyzzeri* progeny**

Single oocysts were sorted into 96-well plates with 1 $\mu$L of TE buffer using the MoFlo XDP-Astrios EQ (Beckman Coulter, CA, USA) cell sorter using a 0.5 drop envelope. Selection of oocysts was based on mNeonGreen and tdTomato fluorescence markers and visualized using the Summit 6.3.1, MoFlo Asterios software (Beckman Coulter). Fecal material of mice infected with the cross was collected from day 17 to day 21 and oocysts were purified. To overcome the anticipated low DNA concentration levels in both bulk and single-oocyst *Cryptosporidium spp.* isolates, extracted DNA from the crosses underwent whole genome amplification (WGA). The amplification was performed using the modified REPLI-g Mini/Midi protocol from Qiagen, employing a multiple displacement amplification (MDA) approach optimized for Oxford Nanopore (ONT) rapid kits. To facilitate ONT long-read sequencing, the amplified sequences were subsequently debranched using T7 endonuclease from New England Biolabs (Manuscript in preparation).

*Variant detection in the progeny with long-read bulk sequences:*

The ONT GridION was used with R9.4.1 flow cells, along with the Rapid barcode kit SQK-RBK110.96. Base calling for ONT data utilized the high-accuracy model in dorado (https://github.com/nanoporetech/dorado). For the parent Illumina data, single nucleotide variant (SNV) calling was conducted by aligning reads with BWA v0.7 [3], duplicate removal using

PICARD v3.0 (http://broadinstitute.github.io/picard/), and SNV calling with GATK 4.2.3.0 Haplotypecaller [4], variants with QUAL < 30.0, DP < 10, MQ < 25.0 and FS > 60.0 were filtered and just SNPs were selected using GATK SelectVariants. For the Oxford Nanopore bulk DNA sequencing from the cross, the alignment was made using Minimap2 v2.26 map-ont [5] and the variant call was performed using NanoCaller v.3.4.0 [6]. For this analysis, we utilized the SNP mode along with the default trained call model, designed specifically for R9.4.1. The *C. parvum* BGF (CpBGF) telomere-to-telomere assembly was used as the reference genome for variant calling (BioProject PRJNA983265 and SRA accession SRR13777123 in [7]. The sequences of the selectable markers from each parent were inserted in the orthologus loci (Chr3 and Chr5) of the reference genome sequence to facilitate read mapping of the parental and progeny reads to a single reference sequence. As the positions of *C. tyzzeri* parental SNPs on the reference genome were known (see below), these locations were used to pull *C. tyzzeri* SNP frequencies from the VCF file created by mapping the bulk reads to the same reference. This approach was taken because of the noise present in the bulk sequencing reads resulting from a population of progeny and ONT sequence errors (the reads are uncorrected). The *C. tyzzeri* SNP frequency was plotted using a custom python script using pandas and matplotlib modules (Appendix SI).

*Recombinant detection in single-oocyts*

ONT GridION R10.4.1 (double pore) flow cells and the Rapid barcode kit SQK-RBK114.24 were used following WGA to generate long-read sequences. Base calling was as above. Sequence reads of both parents and progeny were quality controlled (but not corrected) and aligned to the *Cp*BGF modified reference genome using BWA v.0.7 [3]. Single oocyst coverage (10 read depth minimum required) ranged from 9.1-42.18% depending on the chromosome with an average of 26.25%. The BCF Toolkit [8] was used to call SNVs using -ploidy 1 QUAL <40, MQ <60, DP <10. The SNP distribution was visualized using IGV [9]. The BAM files of the parental and progeny reads in recombinant regions were extracted and converted to fastq files using SAMtools [9]. The extracted single-oocyst progeny long reads were then aligned with parental contigs. Multiple sequence alignments were generated Clustal Omega in Geneious Prime 2022.2.2 with default settings (https://www.geneious.com).

**Chemical synthesis of BRD7929**

**BRD7929** was prepared by adapting previously reported procedures [10]. Specifically, early synthetic steps were modified to increase material throughput from commercially available compound **1** to known intermediate **10a** (Fig. S7). Absolute configuration was established by measurement of the optical rotation of compound **10a** and comparison to reference values [11].

**General information**

NMR spectra were recorded on a Bruker 400 spectrometer (400 MHz $^{1}H$, 101 MHz $^{13}C$). Proton chemical shifts are reported in ppm (δ) referenced to the corresponding NMR solvent [12]. Data are reported as follows: chemical shifts, multiplicity (br = broad, s = singlet, d = doublet, t = triplet, q = quartet, p = pentet, m = multiplet; coupling constant(s) in Hz; integration). Unless otherwise indicated NMR data were collected at 25 °C. Analytical TLC was performed on Merck TLC Silica gel 60 $F_{254}$ plates. Visualization was accomplished with UV light and aqueous potassium permanganate ($KMnO_4$) stain followed by heating. Optical rotation was measured on an Anton Paar MCP 200 polarimeter. Accurate mass measurement analyses (HRMS) were conducted on a Waters LCT Premier XE, time-of-flight, LCMS with electrospray ionization (ESI). The signals were mass measured against an internal lock mass reference of leucine enkephalin. Waters software calibrates the instruments, and reports measurements, by use of neutral atomic masses. The mass of the electron is not included.

**2-((1*R*,2*R*)-1-(4-bromophenyl)-1,3-dihydroxypropan-2-yl)isoindoline-1,3-dione (4)**

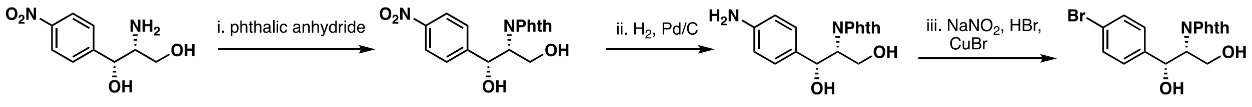


1 2 3 4

To a stirred suspension of (1*R*,2*R*)-2-amino-1-(4-nitrophenyl)propane-1,3-diol (**1**) (150.0 g, 706.9 mmol) in toluene (1.80 L) were added triethylamine (9.90 mL, 70.7 mmol) and phthalic anhydride (105 g, 707 mmol). The resulting mixture was warmed to 100 °C and stirred for 16 h, at which point the suspension became a clear solution. After cooling to room temperature, the reaction mixture was concentrated under reduced pressure. Toluene (100 mL) was added to the residue and the mixture was concentrated under reduced pressure to afford crude 2-((1*R*,2*R*)-1,3-

dihydroxy-1-(4-nitrophenyl)propan-2-yl)isoindoline-1,3-dione (**2**) (240.0 g) as light yellow solid, which was used in the next reaction without further purification.

Compound **2** (150.0 g) was dissolved in MeOH (HPLC grade, 900 mL) and the solution was degassed by bubbling nitrogen for 15 minutes. Next, palladium on activated carbon (10 wt%, 5 g) was added to the solution, and the reaction was placed under hydrogen (30 psi) in a Parr shaker for 16 h. After completion of the reaction, the reaction mixture was filtered over a celite pad. The celite pad was washed with methanol (300 mL × 3). The filtrate was concentrated under vacuum to obtain crude 2-((1*R*,2*R*)-1-(4-aminophenyl)-1,3-dihydroxypropan-2-yl)isoindoline-1,3-dione (**3**) (136.9 g) as a pale yellow foam that solidified upon standing. This material was used in the next reaction without further purification.

To a round-bottom flask containing compound **3** (136.9 g) were added water (500 mL) and HBr (47% solution in water, 1.00 L). The suspension was stirred for 30 min and then cooled in an ice-water bath. A solution of sodium nitrite (33.3 g, 482 mmol) in water (500 mL) was added dropwise. A brown color formed with each added drop and then dissipated upon stirring. After complete addition of sodium nitrite, the brown mixture was stirred for an additional 2 h at 0 °C. In a separate round-bottom flask were introduced CuBr (75.4 g, 526 mmol), water (400 mL), and HBr (47% solution in water, 200 mL), then the diazonium salt solution was added to this suspension slowly at 0 °C. After addition, the reaction mixture was stirred for 16 hours at room temperature. The resulting mixture was extracted with EtOAc (600 mL × 3). The combined organic layers were washed with water (500 mL × 2), aqueous ammonia solution (500 mL), brine (500 mL), dried over anhydrous sodium sulfate, filtered, and concentrated under vacuum to obtain a crude residue. This residue was washed with MTBE/n-hexane (3:7) to obtain 2-((1*R*,2*R*)-1-(4-bromophenyl)-1,3-dihydroxypropan-2-yl)isoindoline-1,3-dione (**4**) (128 g, 77% yield over three steps) as an off-white solid

$^1$H NMR (400 MHz, DMSO-d$^6$) δ 7.94 – 7.82 (m, 4H), 7.58 (d, *J* = 8.4 Hz, 2H), 7.35 (d, *J* = 8.4 Hz, 2H), 5.68 (d, *J* = 4.3 Hz, 1H), 4.97 (dd, *J* = 9.7, 4.3 Hz, 1H), 4.76 (t, *J* = 6.0 Hz, 1H), 4.23 (td, *J* = 10.2, 4.1 Hz, 1H), 3.91 (td, *J* = 10.9, 5.7 Hz, 1H), 3.06 (ddd, *J* = 11.0, 6.4, 3.9 Hz, 1H).

**2-((1*R*,2*R*)-1-(4-bromophenyl)-1-hydroxy-3-(trityloxy)propan-2-yl)isoindoline-1,3-dione (5)**

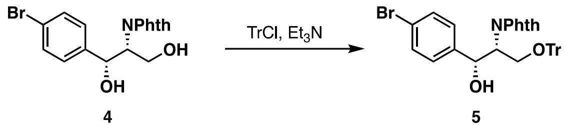

To a stirred, precooled (0 °C) solution of compound **4** (105.5 g, 280.4 mmol) in dichloromethane (1.00 L) was added triethylamine (81.0 mL, 561 mmol). Next, trityl chloride (117.3 g, 420.6 mmol) was added portion-wise at 0°C. The reaction was slowly warmed to room temperature and stirred for 16 h. The reaction was quenched with saturated aqueous ammonium chloride and extracted with dichloromethane (500 mL × 2). The combined organic layers were washed with brine (500 mL), dried over anhydrous sodium sulfate, filtered, and concentrated under reduced pressure to obtain a crude residue, which was purified on a silica gel column (100-200 mesh, eluted at 3:7 ethyl acetate/hexanes) to afford 2-((1*R*,2*R*)-1-(4-bromophenyl)-1-hydroxy-3-(trityloxy)propan-2-yl)isoindoline-1,3-dione (**5**) (139.0 g, 88% purity [EtOAc] by $^{1}$H NMR, 71% yield) as an off-white foam.

$^{1}$H NMR (400 MHz, $CDCl_3$) δ 7.78 – 7.73 (m, 2H), 7.73 – 7.67 (m, 2H), 7.36 – 7.27 (m, 9H), 7.21 – 7.12 (m, 10H), 5.16 (dd, *J* = 9.7, 5.4 Hz, 1H), 4.90 (dt, *J* = 9.3, 4.7 Hz, 1H), 4.43 (d, *J* = 9.6 Hz, 1H), 3.72 (t, *J* = 9.5 Hz, 1H), 3.62 (dd, *J* = 10.3, 4.1 Hz, 1H).

**2-(((1*R*,2*R*)-1-(4-bromophenyl)-1-hydroxy-3-(trityloxy)propan-2-yl)amino)acetonitrile (7)**

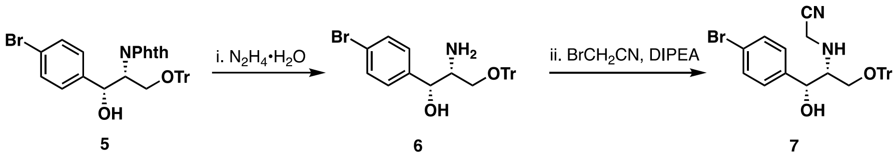

To a stirred solution of compound **5** (139.0 g, 88% purity [EtOAc], 198 mmol) in ethanol (500 mL) was carefully added hydrazine monohydrate (21.8 mL, 449 mmol). The mixture was warmed to 80 °C and stirred for 16 hours. The reaction was cooled to ambient temperature and the solid generated during the reaction was filtered off. The filtrate was concentrated to afford crude (1*R*,2*R*)-2-amino-1-(4-bromophenyl)-3-(trityloxy)propan-1-ol (**6**) (86.5 g) as a white solid, which was used in the next reaction without further purification.

To a stirred solution of compound **6** (59.0 g) in dry acetonitrile (500 mL) were added *N*,*N*-diisopropylethylamine (DIPEA, 44.6 mL, 121 mmol) and bromoacetonitrile (10.1 mL, 145 mmol) portion-wise, and the reaction mixture was warmed to 70 °C and stirred for 5 h. Next, the reaction mixture was cooled to ambient temperature, quenched with cold water and extracted with ethyl acetate (200 mL × 3). The combined organic phase was sequentially washed with water (50 mL × 2), saturated aqueous ammonium chloride (100 mL × 2), and brine (50 mL × 1). The combined organic layers were dried over anhydrous sodium sulfate, filtered, and concentrated under reduced pressure. The resulting crude was purified by column chromatography (silica gel 100-200 mesh, eluted at 1:9 ethyl acetate/hexanes) to afford 2-(((1*R*,2*R*)-1-(4-bromophenyl)-1-hydroxy-3-(trityloxy)propan-2-yl)amino)acetonitrile (**7**) (51.7 g, 83% purity [EtOAc] by $^1$H NMR, 56% yield over two steps) as a brown amorphous solid.

$^1$H NMR (400 MHz, $CDCl_3$) δ 7.44 – 7.20 (m, 17H), 7.15 (d, *J* = 8.3 Hz, 2H), 4.66 (d, *J* = 7.1 Hz, 1H), 3.37 – 3.29 (m, 3H), 3.08 (dd, *J* = 10.5, 4.4 Hz, 1H), 2.86 (dt, *J* = 7.6, 4.0 Hz, 1H).

**2-(allyl((1*R*,2*R*)-1-(4-bromophenyl)-1-hydroxy-3-(trityloxy)propan-2-yl)amino)acetonitrile (8)**

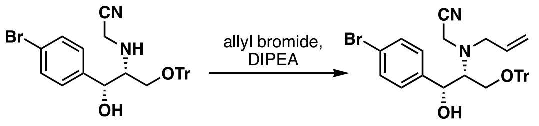


7 8

To a stirred solution of compound **7** (51.7 g, 83% purity [EtOAc], 81.3 mmol) in acetonitrile (250 mL) was added DIPEA (54.2 mL, 294 mmol) at 28 °C. The mixture was heated to 80 °C, then allyl bromide (18.7 mL, 216 mmol) was added, and the reaction was stirred at 80 °C for 16 h. The reaction mixture was cooled to room temperature, quenched with water, and extracted with ethyl acetate (350 mL × 2). The combined organic phase was sequentially washed with water (200 mL × 2), saturated aqueous ammonium chloride solution (200 mL × 2), and brine (100 mL). The combined organic layers were dried over anhydrous sodium sulfate, filtered, and concentrated under reduced pressure. The crude material was purified by column chromatography (silica: 100-200 mesh, eluted at 1:9 ethyl acetate/hexanes) to obtain 2-(allyl((1*R*,2*R*)-1-(4-bromophenyl)-1-hydroxy-3-(trityloxy)propan-2-yl)amino)acetonitrile (**8**) (47.4 g, 92% purity [EtOAc] by $^1$H NMR, 95% yield) as a brown amorphous solid. Analytical data are in agreement with previous reports [11].

$^1$H NMR (400 MHz, $CDCl_3$) δ 7.42 – 7.37 (m, 2H), 7.34 – 7.18 (m, 15H), 7.10 (dd, *J* = 8.8, 2.3 Hz, 2H), 5.80 – 5.67 (m, 1H), 5.28 – 5.17 (m, 3H), 4.37 (d, *J* = 9.3 Hz, 1H), 3.76 – 3.60 (m, 2H), 3.45 (dd, *J* = 14.0, 5.5 Hz, 1H), 3.34 – 3.23 (m, 1H), 3.18 (dd, *J* = 10.9, 3.5 Hz, 1H), 3.11 – 2.99 (m, 2H).

**(2*R*,3*R*,4*S*)-1-allyl-3-(4-bromophenyl)-4-((trityloxy)methyl)azetidine-2-carbonitrile (10a)**

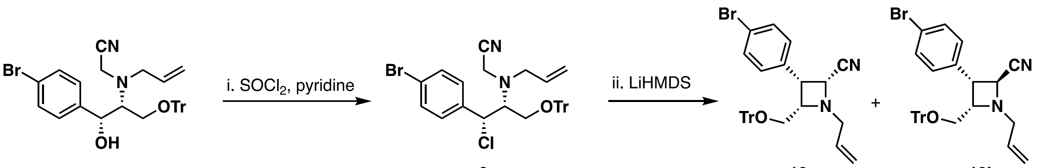


To a precooled (0 °C) solution of pyridine (33.8 mL, 418 mmol) in dry dichloromethane (200 mL) was added thionyl chloride (12.2 mL, 167 mmol) in a dropwise manner, and the resulting mixture was stirred for 15 min. A solution of compound **8** (47.4 g, 92% purity [EtOAc], 77 mmol) in dry dichloromethane (200 mL) was added dropwise at 0 °C. After stirring for 20 min at 0 °C, the reaction mixture was quenched by slow addition of sat. aq. $NaHCO_3$ and stirred for 5 minutes. Layers were separated and the aqueous phase was extracted with dichloromethane (300 mL × 2). The combined organic layers were washed with water (500 mL × 2), followed by brine (500 mL), dried over anhydrous $Na_2SO_4$, filtered, and concentrated under reduced pressure to obtain crude 2-(allyl((1*R*,2*R*)-1-(4-bromophenyl)-1-chloro-3-(trityloxy)propan-2-yl)amino) acetonitrile (**9**) (60.0 g), which was used in the next reaction without further purification.

A solution of crude compound **9** (60.0 g) in dry THF (800 mL) was cooled to –50 °C under nitrogen atmosphere. Lithium bis(trimethylsilyl)amide (LiHMDS, 1 M solution in THF, 155 mL, 155 mmol) was added dropwise and the reaction mixture was stirred for 1 h at –50 °C. The reaction mixture was quenched with sat. aq. $NH_4Cl$ and the aqueous layer was extracted with EtOAc (× 2). The combined organic extracts were dried over anhydrous $Na_2SO_4$, filtered and concentrated under reduced pressure to provide the crude product, which was purified by column chromatography (silica: 100-200 mesh) to provide the undesired diastereomer of (2*S*,3*R*,4*S*)-configuration (**10b**, eluted with 5:95 ethyl acetate/hexane) followed by (2*R*,3*R*,4*S*)-1-allyl-3-(4-bromophenyl)-4-((trityloxy)methyl)azetidine-2-carbonitrile (**10a**, eluted with 15:85→20:80 ethyl acetate/hexane, 16.7 g, 39% over two steps) as a pale yellow amorphous solid. Analytical data are in agreement with previous reports [11].

$^1$H NMR (400 MHz, $CDCl_3$) δ 7.39 (d, *J* = 8.4 Hz, 2H), 7.32 (d, *J* = 8.5 Hz, 2H), 7.25 – 7.18 (m, 9H), 7.18 – 7.11 (m, 6H), 5.84 – 5.69 (m, 1H), 5.24 (dd, *J* = 17.2, 1.4 Hz, 1H), 5.13 (d, *J* = 10.2 Hz, 1H),

4.10 (d, $J$ = 8.0 Hz, 1H), 3.82 (t, $J$ = 7.8 Hz, 1H), 3.69 (td, $J$ = 7.9, 5.2 Hz, 1H), 3.33 (dd, $J$ = 12.9, 6.3 Hz, 1H), 3.14 (dd, $J$ = 12.9, 7.1 Hz, 1H), 3.06 (dd, $J$ = 9.6, 5.3 Hz, 1H), 2.86 – 2.79 (m, 1H).

$[\alpha]_D^{25}$ = +48.7 (*c* 0.801, $CHCl_3$); reported [11]: **10a**: +29.6 (*c* 1.92, $CHCl_3$), *ent*-**10a**: –50.0 (*c* 2.02, $CHCl_3$).

**(8*R*,9*S*,10*S*)-10-((dimethylamino)methyl)-*N*-(4-methoxyphenyl)-9-(4-(phenylethynyl)phenyl)-1,6-diazabicyclo[6.2.0]decane-6-carboxamide (BRD7929).** BRD7929 (1.2 g) was prepared from compound **10a** by following previously reported procedures [10,11]. Analytical data are in agreement with previous reports [10].

$^1H$ NMR (400 MHz, $CDCl_3$) δ 7.61 – 7.42 (m, 6H), 7.41 – 7.31 (m, 3H), 7.30 – 7.21 (m, 2H), 6.88 – 6.78 (m, 2H), 6.08 (s, 1H), 3.92 – 3.80 (m, 1H), 3.77 (s, 3H), 3.71 – 3.59 (m, 2H), 3.55 (ddd, $J$ = 10.3, 7.6, 2.4 Hz, 1H), 3.50 – 3.39 (m, 1H), 3.33 – 3.18 (m, 1H), 3.11 – 3.00 (m, 1H), 2.87 (dd, $J$ = 14.5, 10.5 Hz, 1H), 2.52 – 2.29 (m, 3H), 2.04 (s, 6H), 1.95 – 1.58 (m, 4H).

$^{13}C$ NMR (101 MHz, $CDCl_3$) δ 155.15, 154.24, 136.53, 131.56, 131.00, 130.51, 127.77, 127.66, 122.69, 121.58, 120.98, 113.46, 88.97, 88.70, 66.23, 66.16, 57.28, 57.04, 54.94, 50.04, 48.22, 45.16, 44.22, 27.37, 26.91.

HRMS (ESI) calcd for $C_{33}H_{39}N_4O_2$ $[M+H]^+$: 523.3073. Found: 523.3085.

**High-resolution mass-spectra of BRD7929**

Accurate mass measurement analyses were conducted on either a Waters GCT Premier, time-of-flight, GCMS with electron ionization (EI), or an LCT Premier XE, time-of-flight, LCMS with electrospray ionization (ESI). Samples were taken up in a suitable solvent for analysis. The signals were mass measured against an internal lock mass reference of perfluorotributylamine (PFTBA) for EI-GCMS, and leucine enkephalin for ESI-LCMS. Waters software calibrates the instruments, and reports measurements, by use of neutral atomic masses. The mass of the electron is not included.

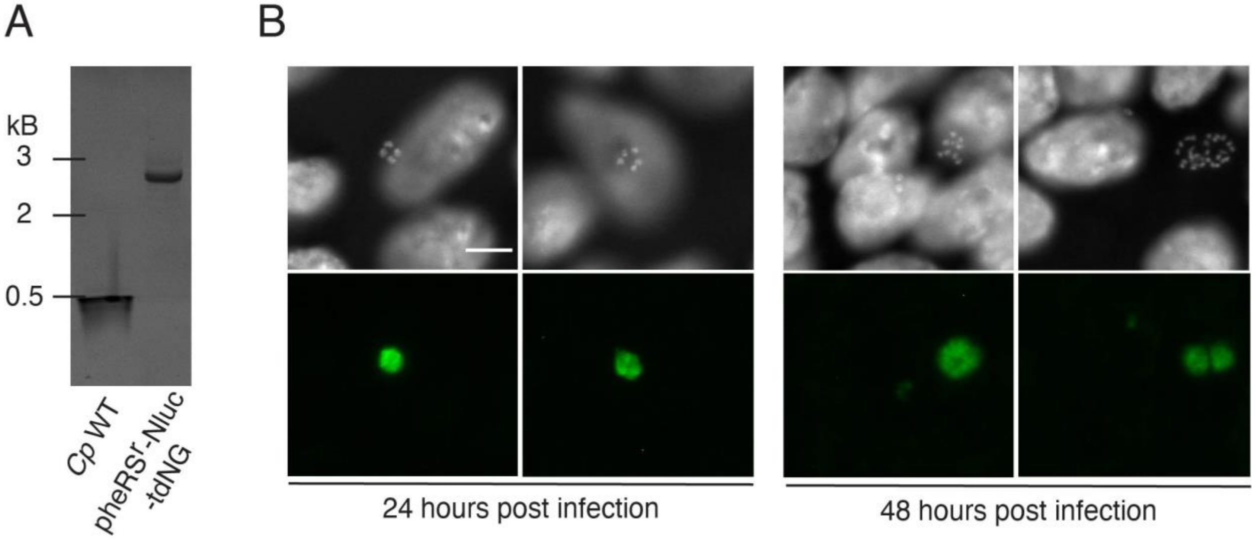


**Figure S1. Genotyping and phenotyping *Cp*-pheRS$^{r}$-Nluc-tdNG. A** Integration PCR gel. PCR mapping using genomic DNA from wild-type (*Cp* WT) and transgenic parasites (*Cp*-pheRS$^{r}$-Nluc-tdNG) showing diagnostic amplicons of the insertion locus. **B** Micrographs of infected HCT-8 cultures after 24 h and 48 h (gray: Hoechst; green: tdNeonGreen). Scale bar, 10 µm.

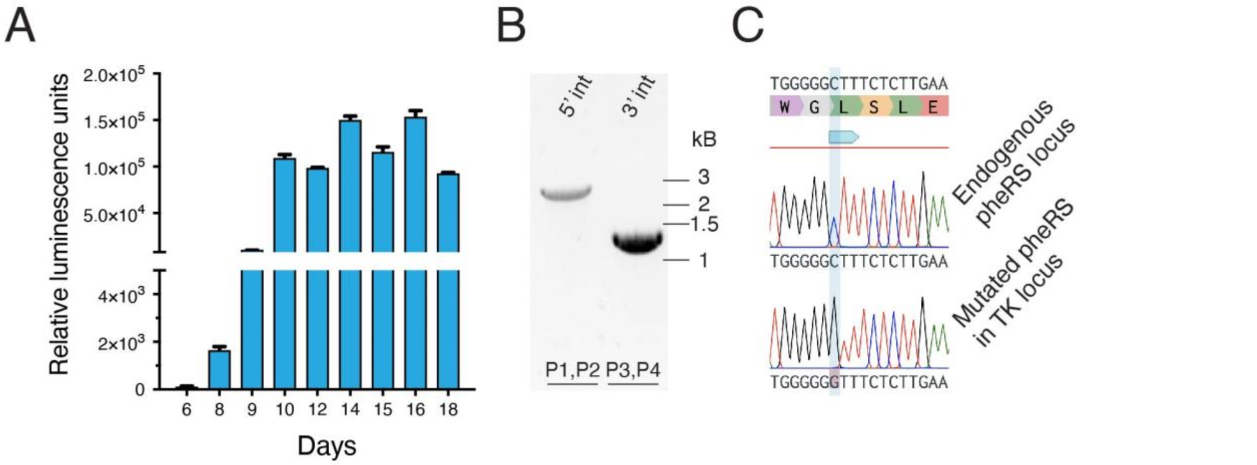


**Figure S2. Further characterization of pheRS$^{r}$-TK-KO. A** Fecal nano luciferase activity of the drug-free passage of the initial pheRS$^{r}$-TK-KO transgenic. **B** Integration PCR gel. PCR mapping showing diagnostic 5' and 3' amplicons documenting integration at the target locus. **C** Sequencing of 3' end of PCR amplicons covering the position of the resistance mutation in the endogenous pheRS locus and the pheRS transgene in the TK. Note that the endogenous locus remained WT and its product thus susceptible to BRD7929.

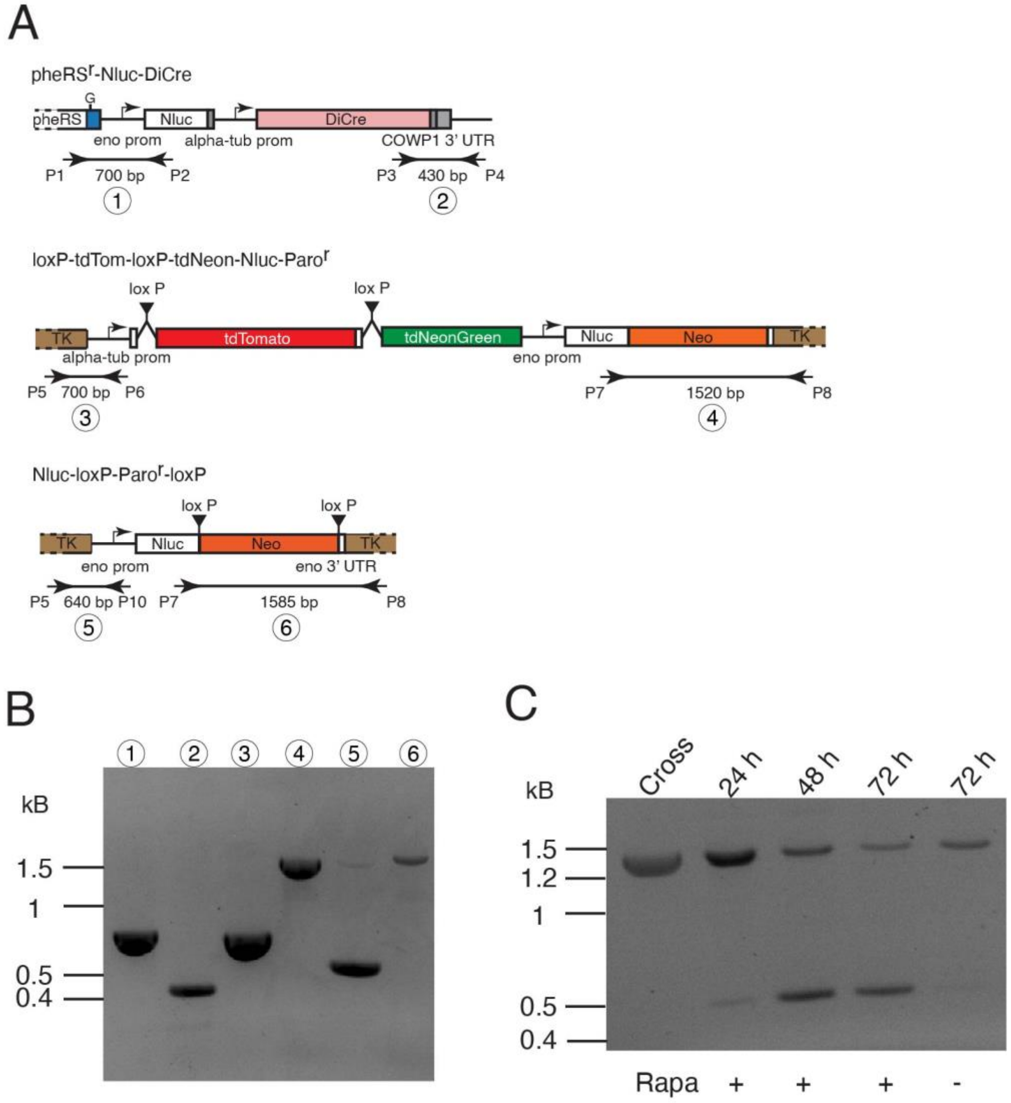


**Figure S3. A** Maps of the pheRS$^r$-Nluc-DiCre strain, the loxP-tdTom-loxP-tdNeon-Nluc-Paro$^r$ strain, and the loxP-Paro$^r$-loxP strains used for crosses. **B** Gel shows PCR mapping of the parental lines (encircled numbers refer to the amplicons shown in the maps in **A)**. **C** Cross progeny was used to infect HCT-8 cell cultures for the indicated times in the presence or absence of rapamycin. Genomic DNA was extracted and used for PCR analysis targeting amplicon 6 prior to (1269 bp) and following rapamycin induced Cre-mediated excision (432bp).

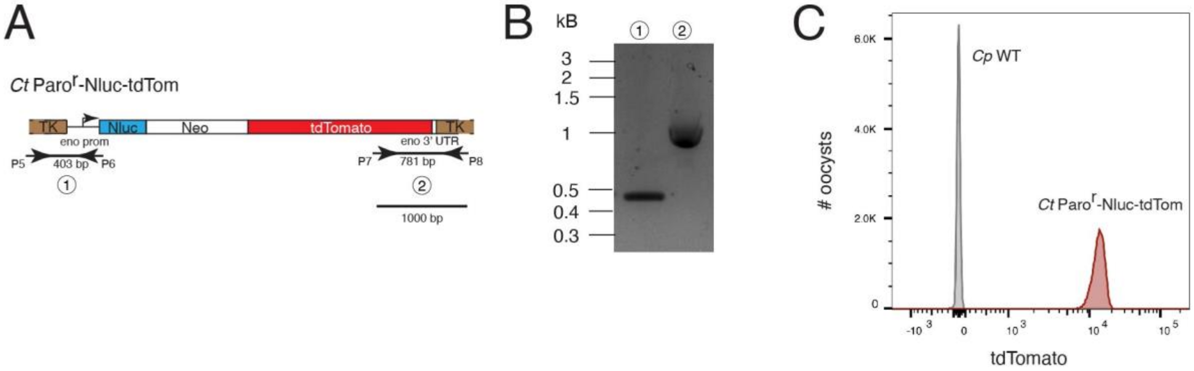


**Figure S4. Genotypic and phenotypic characterization of the transgenic *C. tyzzeri* strain used in cross. A** Map of the parental *C. tyzzeri* line used for the cross. **B** Integration PCR gel. PCR mapping using genomic DNA from transgenic parasites showing transgene insertion into the TK locus. **C** Flow cytometry analysis of purified oocysts showing red fluorescence in the transgenic but not the WT *C. tyzzeri*.

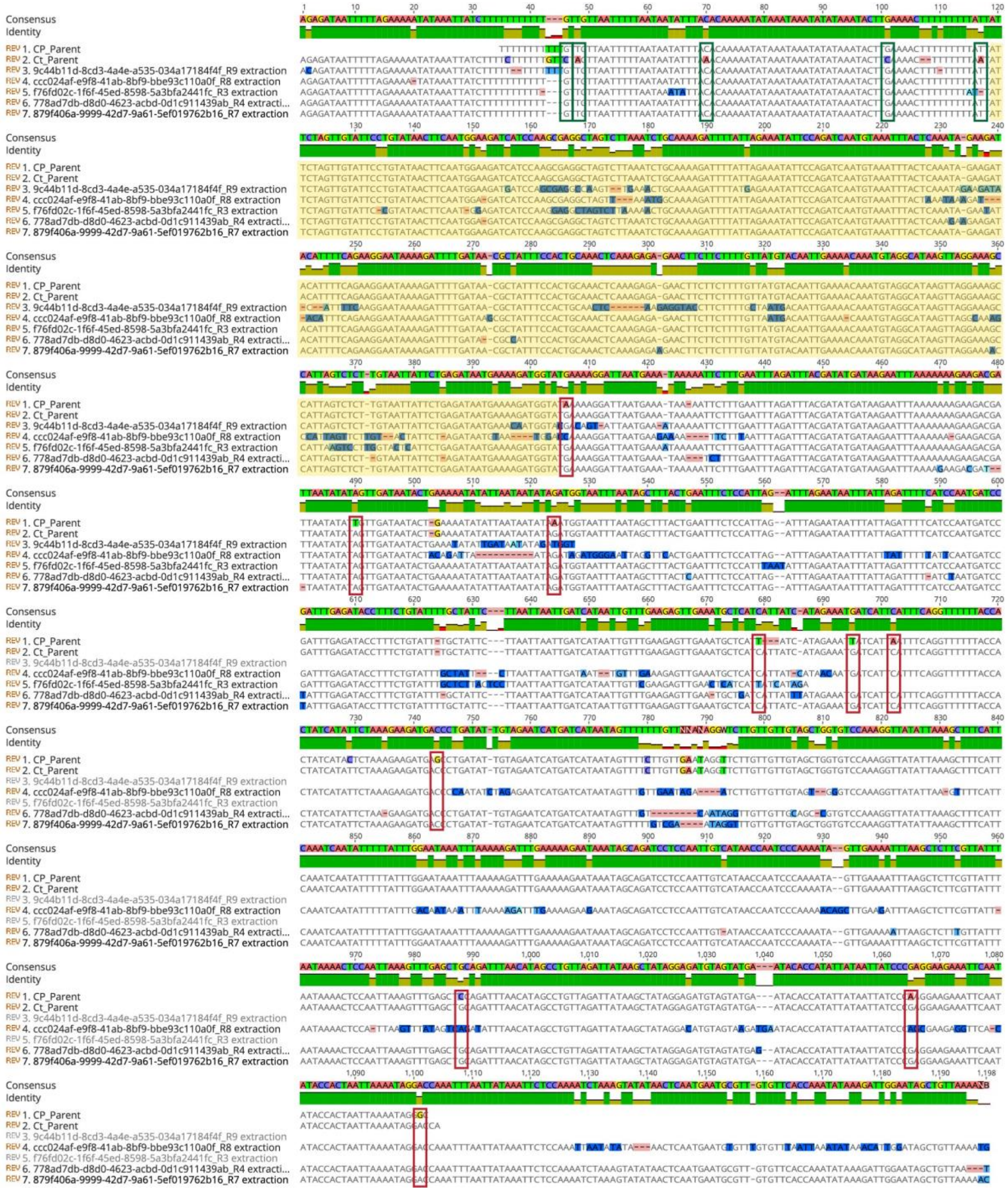

**Figure S5.** Multiple sequence alignment of the center of the region shown in Fig. 5G. The rows represent the *C. parvum* and *C. tyzzeri* parental consensus sequences and a portion of 5 single, long-molecule, Oxford Nanopore reads. The complete reads mapping to the recombinant region identified with single-oocyst sequencing on chromosome 8 can be found in BioProject PRJNA1000584 (BioSample SAMN36772389). Green vertical boxes indicate *C. parvum* specific

SNPs, red boxes highlight *C. tyzzeri* specific SNPs and the large yellow shaded area defines the region of recombination. Based on the SNVs present in the parents, the crossover happened somewhere between positions 118 and 405 on the alignment. In the consensus sequence, each of the 4 nucleotides, gaps and ambiguous bases are identified by a different color. The identity track indicates the level of read consensus at each position with green being best and red the worst. See additional detail in Supplemental data alignment that contains 50 out of 408 reads for this region.

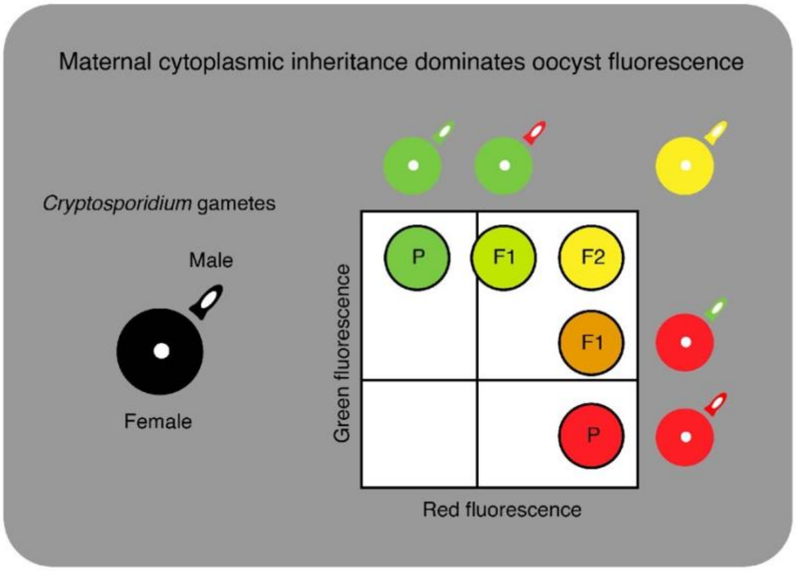


**Figure S6. Hypothetical model of maternal inheritance.** Female gametes are much larger than male gametes and dominate the proteome of the oocyst. The hypothetical flow cytometry plot shows the consequence of paternal or maternal inheritance of each fluorescent reporter shown for various gamete pairings. Based on this hypothesis, the first cross (F1) will initially produce oocysts of intermediate fluorescence, and 'fully' double fluorescent oocysts will be the product (F2) of a fertilization event where both parents carry both transgenic reporters.

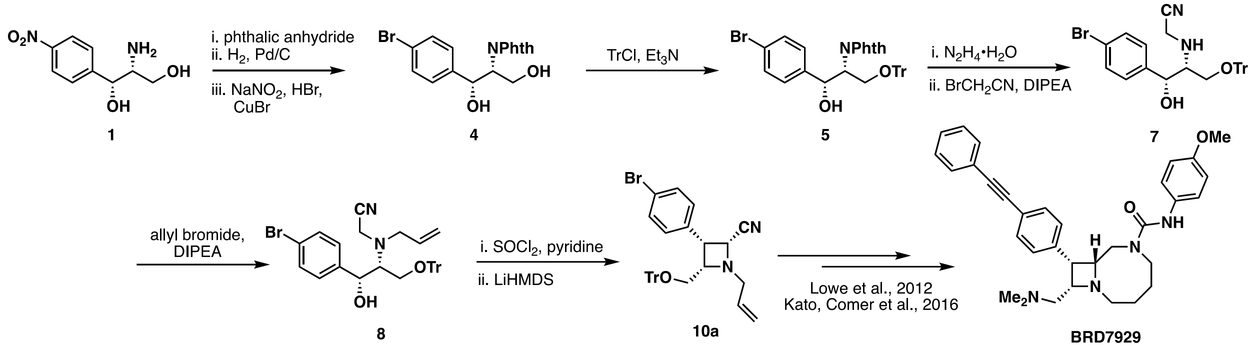

**Fig. S7.** Modified preparation of intermediate **10a** in the synthesis of **BRD7929**.

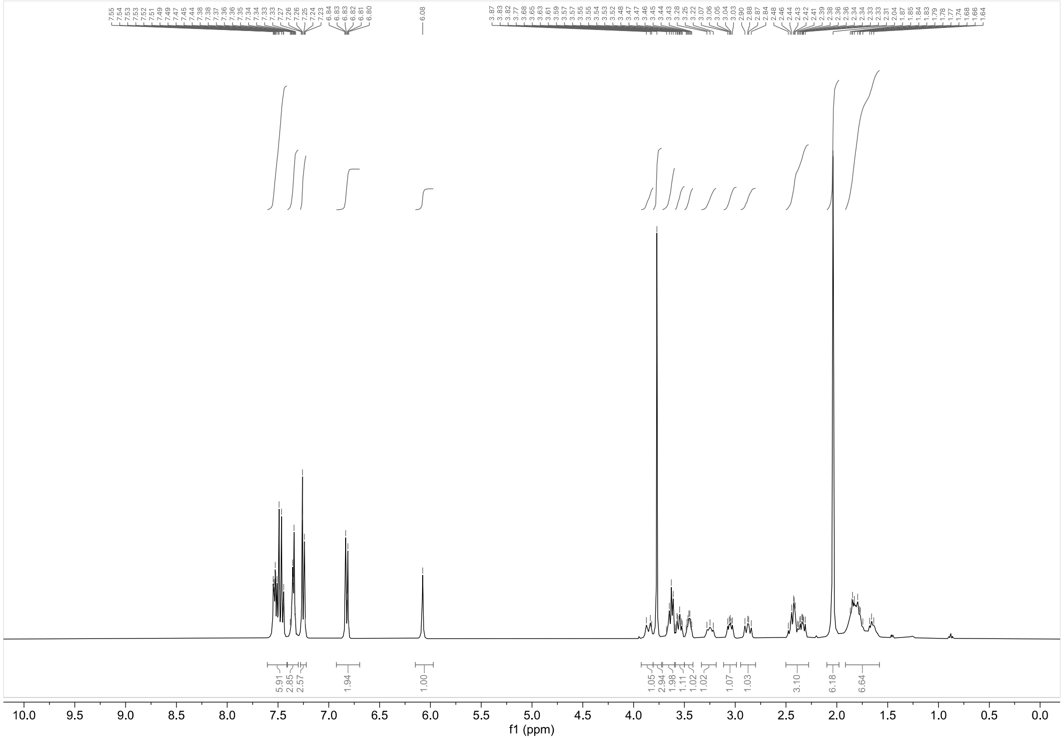

**Fig. S8.** $^1$H NMR spectrum of BRD7929 in $CDCl_3$ (400 MHz).

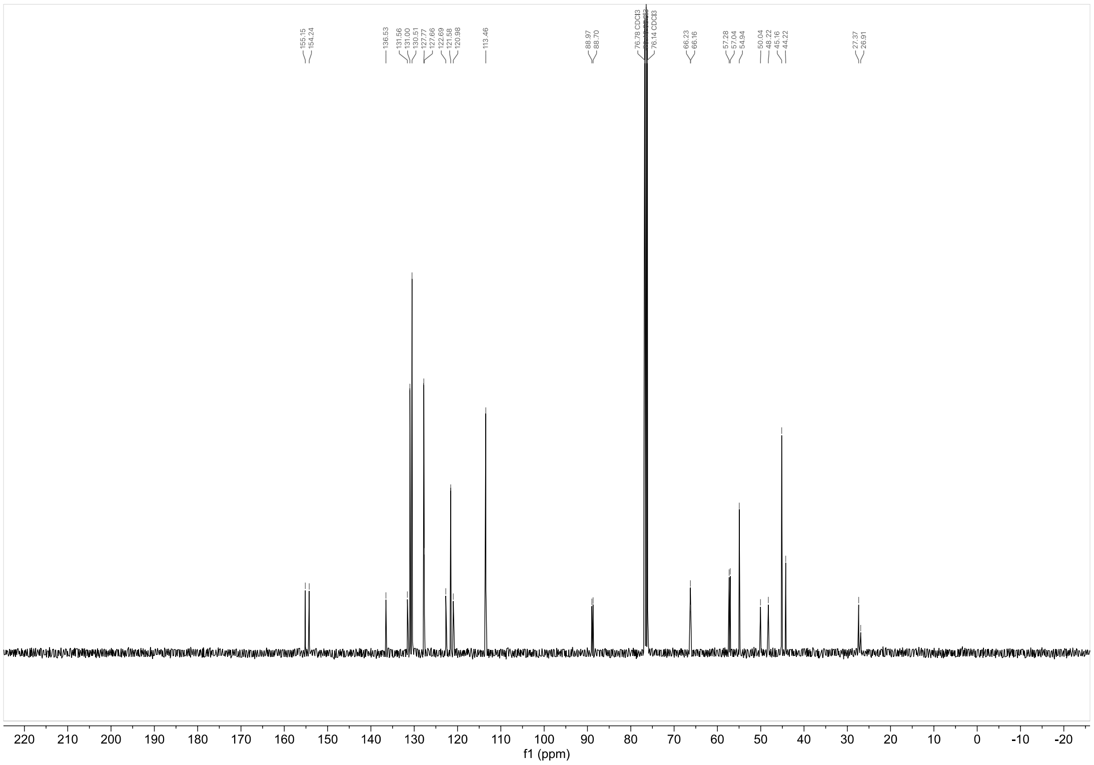


**Fig. S9.** $^{13}$C NMR spectrum of BRD7929 in $CDCl_3$ (101 MHz).

#### Table S1: Oligonucleotides for guide RNAs and primers for repair template amplification used in this study

| Name | Sequence |
|---|---|
| PheSF_guide_New_SV | GTTGGGCAATCACAACGATGTCAG |
| PheSR_guide_New_SV | AAACCTGACATCGTTGTGATTGCC |
| ohgF_PheS_N_wmpam | gttggaaattccggcttatttagaccagaaatgcttagaccactcgggtttccAtctgacatcgttgtgattgc |
| ohg_PheR | agtgattataatctaaaaaacaaacattaattcgaaaagtctacattaggAATTAAGATAAAAAGAAAAAC |
| TK_int_fwd | GTTGGAAGTAAATACTTATTAGCA |
| TK_int_rev | AAACTGCTAATAAGTATTTACTTC |
| pheRSKO fwd | TCCAGTACTATGCTATGGTTTGAGAACAGACTTTAAGGGAAATTTATTTGAAAGCGGATCTTGCCAATCTT |
| pheRSKO rev | TAGCTTTTTTGCCACAGCGACAAATAGTTTTGATTTCAGTAAGTTTATCAAATTAAGATAAAAAGAAAAACTTAATCGATACTATCCTAC |
| loxP_repair_F | TCCAGTACTATGCTATGGTTTGAGAACAGACTTTAAGGGAAATTTATTTGAtggggaaactaaatatactgaaattcgg |
| AG160 | TAGCTTTTTTGCCACAGCGACAAATAGTTTTGATTTCAGTAAGTTTATCAcgcgtttaaactgattggtactag |
| Primer 8522 | TCCAGTACTATGCTATGGTTTGAGAACAGACTTTAAGGGAAATTTATTTGATGGGGAAACTAAATATACTGAAATTCGGT |

#### Table S2: Primers used to genotype transgenic strains

| Name | Original name | Sequence | Figure | Remarks |
|---|---|---|---|---|
| P1 | pheRSmut_100bp_upstr | CCTGCATTTAACCCTTATACTGAGC | S1 | |
| P2 | phe_3int_rev | TGTGCGTGTATCGGGAGTC | S1 | |
| P1 | TKf (5390) | ATGGCAAAATTATACTTTTACTATTCAGCAATGAATGC | 2 | |
| P2 | pheRS_qPCR_R_mut | cactaattcatattcccgcccttagag | 2 | |
| P3 | pheRSmut_100bp_upstr | CCTGCATTTAACCCTTATACTGAGC | 2 | |
| P4 | TKr (5391) | TTAGAAATTGTATTCTTCACAATTAATTATATGATGTTTTCTGC | 2 | |
| P5 | phe_3int_rev | TGTGCGTGTATCGGGAGTC | 2 | |
| P1 | TKf (5390) | ATGGCAAAATTATACTTTTACTATTCAGCAATGAATGC | S2 | |
| P2 | pheRS_qPCR_R_mut | cactaattcatattcccgcccttagag | S2 | |
| P3 | pheRSmut_100bp_upstr | CCTGCATTTAACCCTTATACTGAGC | S2 | |
| P4 | TKr (5391) | TTAGAAATTGTATTCTTCACAATTAATTATATGATGTTTTCTGC | S2 | |
| P1 | pheRSmut_100bp_upstr | CCTGCATTTAACCCTTATACTGAGC | 3 | |
| P2 | AG95 | gctgtcccgtgagatataga | 3 | |
| P3 | mNGF | GTTTCTAAGGGTGAAGAAGATAACATGG | 3 | |
| P4 | phe_3int_rev | TGTGCGTGTATCGGGAGTC | 3 | |
| P5 | TKf (5390) | ATGGCAAAATTATACTTTTACTATTCAGCAATGAATGC | 3 | |
| P6 | AG95 | gctgtcccgtgagatataga | 3 | |
| P7 | AG45 | ACTGCCTTTCGCTTGGGATA | 3 | |
| P8 | TKr (5391) | TTAGAAATTGTATTCTTCACAATTAATTATATGATGTTTTCTGC | 3 | |
| P1 | pheRSmut_100bp_upstr | CCTGCATTTAACCCTTATACTGAGC | S3 | |
| P2 | Primer 3810 | CTTTGGATCGGAGTTACGGACAC | S3 | |
| P3 | 94_Pf_LIC_neon_3'int | AACGACAAGACCATCATCAG | S3 | |
| P4 | phe_3int_rev | TGTGCGTGTATCGGGAGTC | S3 | |
| P5 | TKf (5390) | ATGGCAAAATTATACTTTTACTATTCAGCAATGAATGC | S3 | |
| P6 | AG95 | gctgtcccgtgagatataga | S3 | |
| P7 | AG45 | ACTGCCTTTCGCTTGGGATA | S3 | |
| P8 | TK_seq_rev | AGAAAAGCATCCTCCTTTGTAACC | S3 | |
| P1 | Primer 3809 | CACTATGGCACACTGGTAATCGAC | 4 | |
| P2 | TKr (5391) | TTAGAAATTGTATTCTTCACAATTAATTATATGATGTTTTCTGC | 4 | |
| P5 | TKf (5390) | ATGGCAAAATTATACTTTTACTATTCAGCAATGAATGC | S4 | |
| P6 | Primer 7718 | GTTTAACGAATAACTGTTTAACGAATAACTTTAAC | S4 | |
| P7 | Primer 3809 | CACTATGGCACACTGGTAATCGAC | S4 | |
| P8 | TK_seq_rev | AGAAAAGCATCCTCCTTTGTAACC | S4 | |
| P1 | AG45 | ACTGCCTTTCGCTTGGGATA | 5 | TK locus |
| P2 | TK_seq_rev | AGAAAAGCATCCTCCTTTGTAACC | 5 | TK locus |
| P3 | pheRSmut_100bp_upstr | CCTGCATTTAACCCTTATACTGAGC | 5 | pheRS locus |
| P4 | pheRS_qPCR_R_mut | cactaattcatattcccgcccttagag | 5 | pheRS locus |
| P5 | F2 | TTCTAGAGCTAATACATGCG | 5 | 18S locus 1st PCR |
| P6 | RX | CCCATTTCCTTCGAAACAGGA | 5 | 18S locus 1st PCR |
| P7 | F3N | GGAAGGGTTGTATTTATTAGATAAAG | 5 | 18S locus 2nd PCR |
| P8 | R3N | AAGGAGTAAGGAACAACCTCCA | 5 | 18S locus 2nd PCR |
| P9 | GP60_1 | ATA GTC TCC GCT GTA TTC | 5 | gp60 locus 1st PCR |
| P10 | GP60_3 | GGA AGG AAC GAT GTA TCT | 5 | gp60 locus 1st PCR |
| P11 | GP60_2 | TCC GCT GTA TTC TCA GCC | 5 | gp60 locus 2nd PCR |
| P12 | GP60_4 | GCA GAG GAA CCA GCA TC | 5 | gp60 locus 2nd PCR |

#### Supplementary Code

##AF_plot.py – by Rodrigo P. Baptista

```
import pandas as pd
import matplotlib.pyplot as plt

# Step 1: Get the file paths from the command line
import sys
if len(sys.argv) != 3:
    print("Usage: python script.py path/to/data1(AF_file).csv
path/to/data2(Chr_lengths).csv")
    sys.exit(1)

data1_path = sys.argv[1]
data2_path = sys.argv[2]

# Step 2: Read the data from CSV files without header
data1 = pd.read_csv(data1_path, header=None, names=["Chromosome", "Position", "ALT_AF"
])
data2 = pd.read_csv(data2_path, header=None, names=["Chromosome", "ChromosomeLength"])

# Step 3: Merge the data based on the specified chromosome column
chromosome_col = "Chromosome"  # Change this if the column name is different
df_merged = data1.merge(data2, on=chromosome_col)

# Step 4: Plot scatter plot with red dots and lines connecting each point to the x-
axis for each chromosome
unique_chromosomes = df_merged[chromosome_col].unique()

for chromosome in unique_chromosomes:
    df_chromosome = df_merged[df_merged[chromosome_col] == chromosome]
    chromosome_length = df_chromosome["ChromosomeLength"].values[0]

    plt.figure()
    plt.scatter(df_chromosome["Position"], df_chromosome["ALT_AF"], s=1, color="red")
    for idx, row in df_chromosome.iterrows():
        plt.vlines(row["Position"], 0, row["ALT_AF"], color="red", linestyle="-")
    plt.xlabel("Chromosome Position")
    plt.ylabel("Frequency (ALT AF)")
    plt.title(f"Chromosome {chromosome}")
    plt.axhline(y=0.5, color="black", linestyle="--")
    plt.xlim(0, chromosome_length)
    plt.ylim(0, 1)
    plt.savefig(f"scatter_with_lines_chromosome_{chromosome}.svg")
    plt.close()
```

**Supplementary Data**

**50 read alignment in recombinant region (*C. parvum* x *C. tyzzeri*) of chromosome 8 –** The sequences of each parent are shown along with 50 (out of 408) Oxford Nanopore reads that span the recombinant region. Each uncorrected read is shown with its original ID that corresponds to the reads submitted to the SRA. Vertical columns of interest are highlighted with gray. Green text indicates SNPs indicative of *C. parvum* and red indicates SNPs indicative of *C. tyzzeri*. The large block of grey alignment represents the crossover region. No SNPs are available to map the region more finely.

```
Cp_PARENT                             AATTTTA-TTAAAGGAAGAAGAGGTA-TATCGCA-AATTCGAT-GGAGGAAAGAAACCTCTTT------AGAATCAGAA- [   80]
Ct_PARENT                             AATTTTA-TTAAAGGAAGGAGAGGTA-TATCGCA-AATTCGAT-GGAGGAAAGAAACTTCTTT------AGAATCAGAA- [   80]
bdf46f5b-2833-4b26-892f-323dd4c3  AATTTTA-TTAAAGGAAGAAGAGGTA-TATCACA-AATTGGAT-GGAGGAAAGAAACCTCTTTAG-AATAGGAA------ [   80]
c40b234c-b999-47c3-93ca-016af61e  AATTTTA-TTAAAGGAAGAAGAGGTA-TATCGCA-AATTCGAT-GGAGGAAAGAAACCTCTTTAG-AATCAGAA------ [   80]
0d8edcbc-b350-4d00-afbc-fd78a1f6  AATTTTA-TTAAAGGAAGAAGAGGTA-TATCGCA-AATTCGAT-GGAGGAAAGAAACCTCTTTAG-AATCAGAA------ [   80]
9cb07723-65ed-4c96-ac19-4234ea98  ATTTTAT--TAAAGGAAGAAGAGGTA-TATCGCA-AATTCGAT-GGAGGAAAGAAACCTCTTTAG-AATCAGAA------ [   80]
ed18c856-4d5b-4622-9fe8-9b5740af  AATTTTA-TTAAAGGAAGAAGAGGTACTATCGCA-AATTAGAT-GGAGGAAAGAAACCTCTTT-AGAATCCGAA------ [   80]
c344612d-f873-4743-818c-65979638  AATTTTA-TTAAAGGAAGAAGAGGTA-TAT-----AACGTCGT-GGAGGAAAGAAACCTCTTT-AGAATCAGAA------ [   80]
baaaf62e-d0ee-4c49-9d83-0313d3df  AATTTTA-TTAAAGGAAGAAGAGGTA-TATCGCA-AATTGGAT-GGAGGAAAGAAACCTCTTT-AGAATC-GAA------ [   80]
6771788a-69b9-4fd8-b4e0-073881fe  AATTTTA-TTAAAGGAAGAAGAGGTA-TATCGCA-AATTCGAT-GGAGGAAAGAAACCTCTTT-AGAATCAGAA------ [   80]
a6666a5a-6885-427a-9881-430dad33  AATTTTA-TTAAAGGAAGAAGAGGTA-TATCGCA-AATTCGAT-GGAGGAAAGAAACCTCTTTTAGAATCAGAA------ [   80]
5be1c0db-7e16-4441-a2ab-f8107a5d  AATTTTA-TTAAAGGAAG-AGAGGTA-TATCGCA-AATCTGAT-GGAGGAAAGAAACCTCTTCAAGAATCAGAA------ [   80]
99a0bd03-c448-4605-9def-185e9f2d  AATTTTA-TTAAAGGAAGAAGAGGTA-TATCGCA-AATTCGAT-GGAGGAAAGAAACCTCTTT-AGAAT-AGAA------ [   80]
c37ace00-c1f8-46e8-bcdc-c3c33bcb  AA-TTTA-TAAAAGGAAGAAGAGGTA-TATCGCA-AATTCGAT-GGAGGAAAGAAACCTCTTT-AGAATCAGAA------ [   80]
cdc9d647-e2ac-442b-8435-6dd7b82f  AATTTTA-TTAAAGGAAGAAGAGGTA-TATCGCA-AATTGGAT-GGAGGAAAGAAACCTCTTT-AGAATCAGAA------ [   80]
bc10b467-59cf-4610-94fe-267549f8  AGTTTTA-TTAAAGGAAGAAGAGGTA-TATCGCA-AATTCGAT-GGAGGAAAGAAACCTCTTT-AGAATCAGAA------ [   80]
c8c3f0cb-c211-4877-8f07-1765b113  AATTTTA-TTAAAGGAAGAAGAGGTA-TATCGCA-AATTCGAT-GGAGGAAAGAAACCTCTTT-AGAATCAGAA------ [   80]
720a0247-e02e-400b-901f-673d7527  AATTTTA-TTAAAGGAAGAAGAGGTA-TATCGCA-AATTGGAT-GG----AAGAAACCTCTTT-AGAATCAGAA------ [   80]
f3f8defb-2c03-44af-abc8-d94623f7  AATTTTA-TTAAAGGAAGAAGAGGTA-TATCGCA-AATTCGAT-GGAGGAAAGAAACCTCTTTAGAATC-GAA------- [   80]
fc2a9b4c-2c5f-43ed-8ea7-f2af0743  AATTTTA-TTAAAGGAAGAAGAGGTA-TATAGCA-AATTCAAT-GGAGGAAA-AAATGTCTTC-AGAATCAGAA------ [   80]
477953c0-cb9c-400c-bf05-dfadb08e  AATTTTA-TTAAAGGAAGA-GAGGTA-TATCGCA-AATTCAAT-GGAGGAAAGAAACCTCTTT-AGAATCAGAA------ [   80]
28c5afa3-4ebf-4ffa-a71d-2171913e  AATTTAT-TTAAAGGAAGAAGAGGTA-TATCGCA-AATTCGAT-GGAGGAAAGAAACCTATTT-AGAATCAGAA------ [   80]
52a4f968-3394-459b-9ee3-10eab4a7  AATTTTA-TTAAAGGAAGAAGAGGTA-TATCACG-AACTAGAT-GGAGGAAAGAAACCTCTTT-AGAATCAGAA------ [   80]
d36d5471-42a7-440b-bd5e-48d77156  AATTTAT-TTAAAGGAAGA-GAGGTA-TATCGCA-GAATTCGATGGAGGAAAGAAACCTCTTT-AGAATCAGAA------ [   80]
f7886442-23fe-473a-9ead-a2b8a2a7  AATTTTA-TTAAAGGAAGA-GAGGTA-TATCGCA-AATTCAAT-GGAGAAAAG-AACCTCTTT-AGAATCAGAA------ [   80]
f4770134-2a4b-449d-a4f6-c99f3383  AATTTTA-TTAAAAGA--AAGAGGTA-TATC-CA-AATTCGAT-GGAGGAAAGAAACCTCTTT-AGAATCAGAA------ [   80]
4dd53951-3b2d-42af-82ef-399bb712  AATTTTA-TTAAAGGAAGAGAGG----TATGTCA-AATTGGATGGGGGGAAAGAAACCTCTTT-AGAATCAGAA------ [   80]
81ea3bac-8208-46c7-9546-bbd55b7c  AAT-TTA-TTAAAGGAAGAAGAGGTA-TATCGCA-AATTCGAT-GGAGGAAAGAAACCTCTTT-AGAATCAGAA------ [   80]
1e66a4e2-42fd-4fa9-b001-9bb7f263  AATTTTA-TTAAAGGAAGAAGAGGTA-TATCGCA-AATTCGAT-GGAGGAAAGAAACCTCTTTAGAATAGGAAG------ [   80]
b75fad9a-877e-4f52-968b-9fcbb7a4  AATTTTA-TTAAAGGAAGAAGAGGTA-TATCGCA-AATTCGAT-GGAGGAAAAGAACCTCTTTAGAATCAGAAA------ [   80]
f512b649-4f14-4eb6-b5a2-b0543441  AATTTTA-TTAAAGGAAGAAGAGGTA-TATCGCA-AATTCGAT-GGAGGAAAGAAACCTCTTT-AGAATCAGAA------ [   80]
5021e737-de02-42f2-920d-18868672  AATTTTA-TTAAAGGAAGAAGAGGTA-TATCGCA-AATTCGAT-GGAGGAAAGAAACCTCTTTAG-AATCAGAA------ [   80]
```

```
a46adbf5-3ba1-48b8-a762-2403bda7  AATTTTA-TTAAAGGAGAGAGAGGTA-TATAGCA-AATTCGAT-GGAGGAAAGAAACCTCTTTCAGAATCAGAA------ [  80]
088a7e31-4c6b-4ff7-a19b-ff14531f  AATTTTA-TTAAAGGAAGAAGAGGTA-TATCGCA-AATTCGAT-GGAGGAAAGAAACCTCTTTA-GAATCAGAA------ [  80]
e82639d1-c451-4263-90a8-2b1ce6a1  AATTTTA-TTAAAGGAAGAAGAGGTA-TATCGCA-AATTCGAT-GGAGGAAAGAAACCTCTTTCAAGATCAGAA------ [  80]
3cca4221-4eee-44ed-9fb4-ebbac961  AATTTTA-TTAAAGGAAGAAGAGGTA-TATCGCA-AATTCGAT-GGAGGAAAGAAACCTCTTT-CAAATCAGAA------ [  80]
d7a80796-8741-4878-b6ca-23aa66f7  AATTTTA-TTAAAGGAAGAAGAGGTA-TATCGCA-AATTGGAT-GGAGGAAAGAAACCTCTTT-AGAATCAGAA------ [  80]
421e3eda-83a0-4ebd-972c-f24667ee  AATTTTA-TTAAAGGAAGAAGAGGTA-TATCGCA-AATTCGAT-GGAGGAAAGAAACCTCTTT-AGAATCAGAA------ [  80]
e6a39029-4e34-4457-b2a9-0cf6f207  AATTTTA-TTAAAGGAAGAAGAGGTA-TATCGCA-AATTGGAT-GGAGGAAAGAAACCTCTTT-AGAATCAGAA------ [  80]
a1507163-34c9-4b24-b21e-3b5dfc54  ACTTAAATTTAAAGGAAGAAGAGGTA-TATC-CA-AGTCCGTT----GGAAAGAAACCTATTTCA----AAATG---AA- [  80]
51b01061-aaf5-4585-ba72-1a83671b  AAT-TTTATTAAAGAAAGA---GGTA-TATCGCA-AATTGGAT-GGAGGAAAGA-ACCTCTTT------AGAATCAGAA- [  80]
85703d1f-2c69-4541-ab66-acb7b838  AAT-TTTCTTAAAGGAAGA-GAGGTA-TATCGCA-AATTCAAT-GGAGGAAAGAAACCTCTTT------AGAATCAGAA- [  80]
ed39aa3b-d227-4de1-92bd-5ace4420  AAT-TTTCTTAAAGGAAGA---GGTA-TATCGCA-AATTCGAT-GGAGGAAAAGAACCTCTTT-T----AGAATCAGAA- [  80]
40e2af31-875d-446e-90a0-6e16a943  AAT-TTTATTAAAGGAAGA-GAGGTA-TATGGCA-AATTAGAT-GGAGGAAAGAAACCTATAA----------ATCGAA- [  80]
873c76b4-8a29-4846-98df-cb5abcc9  A-ATTTTATTAAAGGAAGAAGAGGT---ATATCAGAATTCGAT-GGAGGAAAGAAACCTCTTT------AGAATCAGAA- [  80]
59ef08a5-aa1d-4d15-ba6f-1a925971  A-ATTTTATTAAAGGAAGAAGAGGTA-TATCGCA-AATTCGAT-GGAGGAAAGAAACCTCTTT------AGAATCAGAA- [  80]
e1cae4d8-4f39-4401-8d82-5c5edaf3  A-ATTTTATTAAAGGAAGAAGAGGTA-TATCGCA-AATTCGAT-GGAGGAAAGAAACCTCTTT------AGAATCAGAA- [  80]
5f47af75-253d-48b0-8b3e-ae59a63d  A-TTTTATTTAAAGGAAGAAGAGGTA-TATCGCA-AATTCGAT-GGAGGAAAGAAACCTCTTT------AGAATATCAG- [  80]
44f1962a-fce6-436a-9ea0-ef4a969d  A-ATTTTCTTAAAGGAAGAAGAGGTA-TATCGCA-AATTGGAT-GGAGGAAAGAAACCTCTTT------AGATAGGAAA- [  80]
4544809e-f4f9-4ff0-8142-5a2a2bc9  AATTTTATTTAAAGGAAGAAGAGGTA-TATCACA-AATTGGAT-G-----AAGAAACCTCTTT------AGAATAGAGGA [  80]
9faa5557-cdc1-466c-815f-fb18a15f  -AATTTTATTAA-AGGAAGAGAGGTA-TATCGCA-AATTCGAT-GGAGGAAAGAAACCTCTTT------AGAATCAGAA- [  80]
ef1a361c-a80e-43f6-8e1f-aeafa681  ATTTTAT--TAAAGGAAGAAGAGGTA-TATCGCA-AATCCGAT-GGAGGAAAGAAACCTCTTTCAGAATCAGAA------ [  80]

Cp_PARENT                         ATTGAAG--ATAAAGAT--AATGA----TTCCAATGAAAAT-G-AG-AT--AGAATTTTGCAGAAAATCAAATTAT--CC [ 160]
Ct_PARENT                         ATTGAAG--ATAAAGAT--AATGA----TTCCAATGAAAAT-G-AG-AT--AGAATTTTGCAGAAAATCAAATTAT--CC [ 160]
bdf46f5b-2833-4b26-892f-323dd4c3  ATTGAAG--ATAAAGAT--AATGA----TTCCAATGAAAAT-G-AG-AT--AGAATTTTGCAGAAAATCAAA-TA-T-CC [ 160]
c40b234c-b999-47c3-93ca-016af61e  ATTGAAG--ATAAAGAT--AATGA----TTCCAATGAAAAT-G-AG-AT--AATTT----CCAGAAAATTAAATA-T-AC [ 160]
0d8edcbc-b350-4d00-afbc-fd78a1f6  ATTGAAG--ATAAAGAT--AATGA----TTCCAATGAAAAT-G-AG-AT--AGAATTTTGCAGAAAATCAAATTA-T-CC [ 160]
9cb07723-65ed-4c96-ac19-4234ea98  ATTGAAG--ATAAAGAT--AATGA----TTCCAATGAAAAT-G-AG-AT--AGAAT--TTTGCAAAATCAAATTA-T-CC [ 160]
ed18c856-4d5b-4622-9fe8-9b5740af  ATTGAAG--ATAAAGAT--AATGA----TTCCAATGAAAAT-G-AG-AT--AAA--TTTGCAGAAAATCAAATTA-T-CC [ 160]
c344612d-f873-4743-818c-65979638  ATTGAAG--ATAAAGAT--AATGA----TTCCAATGAAAAT-G-AG-AT--AGAATTTTGCAGAAAATCAAATTA-T-CC [ 160]
baaaf62e-d0ee-4c49-9d83-0313d3df  ATTGAAG--ATAAAGAT--AATGA----TTCCAATGAAAAT-G-AG-AT--AGAATTTTGCAGAAAATCAAATTA-T-CC [ 160]
6771788a-69b9-4fd8-b4e0-073881fe  ATTGAAG--ATAAAGAT--AATGA----TTCCAATGAAAAT-G-AG-AT--AGAATTTTGCAGAAAATCAAATTA-T-CC [ 160]
a6666a5a-6885-427a-9881-430dad33  ATTGAAG--ATAAAGAT--AATGA----TTCCAATGAAAAT-G-AG-AT--AGAATTTTGCA-AAAATCAAATTA-T-CC [ 160]
5be1c0db-7e16-4441-a2ab-f8107a5d  ATTGAAA--AT-AAGGT--CATGA----TTCCAATGAAAATGG-AG-TT--GGAATTTTGCAGAAAATCAAATTA-T-CC [ 160]
99a0bd03-c448-4605-9def-185e9f2d  ATTGAAG--ATAAAGAT--AATGA----TTCCAATGAAAAT-G-AG-AT--AGAATTTTGCAGAAAATCAAATTA-T-CC [ 160]
c37ace00-c1f8-46e8-bcdc-c3c33bcb  ATTGAAG--ATAAAGAT--AATGA----TTCCAATGAAAAT----G-AG--ATAATTTTGCAGAAAATCAAATTA-T-CC [ 160]
cdc9d647-e2ac-442b-8435-6dd7b82f  ATTGAAG--ATAAAGAT--AATGA----TTCCAATGAAAAT-G-AG-AT--AGAATTTTGCAGAAAATCAAATTA-T-CC [ 160]
bc10b467-59cf-4610-94fe-267549f8  ATTGAAG--ATAAAGAT--AATGA----TTCCAATGAAAAT-G-AG-AT--AGAATTTTGCAGAAAATCAAATTA-T-CC [ 160]
c8c3f0cb-c211-4877-8f07-1765b113  ATTGAAG--ATAAAGAT--AATGA----TTCCAATGAAAAT-G-AG-AT--AATTTTGC--AGAAAATCAAATT--A-TC [ 160]
720a0247-e02e-400b-901f-673d7527  ATTGAAG--ATAAAGAT--AATGA----TTCCAATGAAAAT-G-AG-AT--AGAATTTTGCAGAAAATCAAATTA-T-CC [ 160]
f3f8defb-2c03-44af-abc8-d94623f7  ATTGAAG--ATAAAGAT--AATGA----TTCCAATGAAAAT-G-AG-AT--AGAATTTT-GCAAAAATCAAATTA-T-CC [ 160]
fc2a9b4c-2c5f-43ed-8ea7-f2af0743  ATTGAAG--ATAAAGAT--AATGA----TTCCAATGAAAAT-G-AG------ATAATTTGCAGAAAATGAAATTA-T-CC [ 160]
477953c0-cb9c-400c-bf05-dfadb08e  ATTGAAG--ATAAAGAT--AATGA----TTCCAATGAAAAT-G-AG-AT--AGAATTTTGCAGAAAATCAAATTA-T-CC [ 160]
28c5afa3-4ebf-4ffa-a71d-2171913e  ATTGAAG--ATAAAGAT--AATGA----TTCCAATGAAAAT-G-AG-AT--AGAATTTTGCAGAAAATCAAATTG-T--- [ 160]
52a4f968-3394-459b-9ee3-10eab4a7  ATTGAAG--ATAGAGAT----TGAATGATTCCAATGAAAAT-G-AG-AT--AGAATTTTGCAGAAAATCAAATTA-T-CC [ 160]
d36d5471-42a7-440b-bd5e-48d77156  ATTGAAG--ATAAGAAT--AATGA----TTCCAATGAAAAT-G-AG-AT--AG-ATTTTGCAGAAAATCAAATTA-T-CC [ 160]
```

```
f7886442-23fe-473a-9ead-a2b8a2a7  ATTGAAG--ATAAAGAT--AATGA----TTCCAATGAAAAT-G-AG-AT--AGAATTTTGCAGAAAATCAAATTA-TCCC [ 160]
f4770134-2a4b-449d-a4f6-c99f3383  ATTGAAG--ATAAAGAT--AATGA----TTCCAATGAAAAT-A-ACAAT--AGAATTTTGCAGAAAATCAAATTA-T-CC [ 160]
4dd53951-3b2d-42af-82ef-399bb712  ATTGAAG--ATAAAGAT--AATGA----TTCCAATGAAAAT-G-AATAA--ATT--------ACAAATCAAATTA-T-CC [ 160]
81ea3bac-8208-46c7-9546-bbd55b7c  ATTGAAG--ATAAAGAT--AATGA----TTCCAATGAAAAT-G-AG-AT--AGAATTTTGCAGAAAATCAAATTA-T-CC [ 160]
1e66a4e2-42fd-4fa9-b001-9bb7f263  ATTGAAG--ATAAAGAT--AATGA----TTCCAATGAAAAT-G-AG-AT---AAAATTTGCAGAAAATCAAATTA-T-CC [ 160]
b75fad9a-877e-4f52-968b-9fcbb7a4  TT-GAAG--ATAAAGAT--AATGA----TTCCAATGAAAAT-G-AG-AT--AGAATTTTGCAGAAAATCAAATTA-T-CC [ 160]
f512b649-4f14-4eb6-b5a2-b0543441  ATTGAAG--ATAAAGAT--AATGA----TTCCAATGAAAAT-G-AG-AT--AGAATTTTGCAGAAAATCAAATTA-T-CC [ 160]
5021e737-de02-42f2-920d-18868672  ATTGAAG--ATAAAGAT--AATGA----TTCCAATGAAAAT-G-AG-AT--AGAATTTTGCAGAAAATCAAATTA-T-CC [ 160]
a46adbf5-3ba1-48b8-a762-2403bda7  ATTGAAG--ATAAAGAT--AATGA----TTCCAATGGAATG-T-GG-AT--AAAATT-----ACAAATTAAATTA-T-CC [ 160]
088a7e31-4c6b-4ff7-a19b-ff14531f  ATTGAAG--ATAAAGAT--AATGA----TTCCAATGAAAAT-G-AG-AT--AGAATTTTGCAGAAAATCAAATTA-T-CC [ 160]
e82639d1-c451-4263-90a8-2b1ce6a1  ATTGAAG--ATAAAGAT--AATGA----TTCCAATGAAAAT-G-AG-AT--AGAATTTTGCAGAAAATCAAATTA-T-CC [ 160]
3cca4221-4eee-44ed-9fb4-ebbac961  ATTGAAG--ATAAAGAT--AATGAATTTTTCCAATGAAAAT-G-AG-AT--AGAATTTTGCAGAAAATCAAATTA-T-CC [ 160]
d7a80796-8741-4878-b6ca-23aa66f7  ATTGAAG--ATAAAGTA--GATGA----TTCCAATGAAAAT-G-AG-AT--AGAATTTTGCAGAAAATCAAATTA-T-CC [ 160]
421e3eda-83a0-4ebd-972c-f24667ee  ATTGAAG--ATAAAGAT--AATGA----TTCCAATGAAAAT-G-AG-AT--AGAATTTTGCAGAAAATCAAATTA-T-CC [ 160]
e6a39029-4e34-4457-b2a9-0cf6f207  ATTGAAG--ATAAAGAT--AATGA----TTCCAATGAAAAT-G-AG-AT--AGAATTTTGCAGAAAATCAAATTA-T-CC [ 160]
a1507163-34c9-4b24-b21e-3b5dfc54  ATTGAA--AATAATAAT--GATT--------GAATGAAAAT-G-AG-ATA-AAT----TTGCCAAAATCAAATTAT--CC [ 160]
51b01061-aaf5-4585-ba72-1a83671b  ATTGAA--GATAAAGAT--AATGA----TTCCAATGAAAAT-G-AG-A-T-AGAATTTTGCCAAAATAGAAT------TA [ 160]
85703d1f-2c69-4541-ab66-acb7b838  ATTGAA--GATAAAGAT--AATGA----TTCCAATGAAAAT-G-AG-ATA-AAT----TTGCAAAAATCAAA------TT [ 160]
ed39aa3b-d227-4de1-92bd-5ace4420  ATTGAA--GATAAAGAT--AATGA----TTCCAATGAAAAT-G-AG-A-T-AGAATTTTGCAGAAAATCAAATTAT--CC [ 160]
40e2af31-875d-446e-90a0-6e16a943  ATTGAA--GATAAAGAT--AATGA----TTCCAATGAAAAT-G-AG-A-T-AGAATTTTGCAGAAAATCAAATTAT--CC [ 160]
873c76b4-8a29-4846-98df-cb5abcc9  ATTGAA--GATAAAGAT--AATGA----TTCCAATGAAAAT-G-AG-AT--AAAT-TTTGCAGAAAATCAAATTAT--CC [ 160]
59ef08a5-aa1d-4d15-ba6f-1a925971  ATTGAA--GATAAAGAT--AATGA----TTCCAATGAAAAT-C-AG-AT--AGAATTTTGCA-AAAATCAAATTAT--CC [ 160]
e1cae4d8-4f39-4401-8d82-5c5edaf3  ATTGAA--GATAAAGATACAATGA----TTCCAATGAAAAT-G-AG-AT---GAATTTTGCAGAAAATCAAATTAT--CC [ 160]
5f47af75-253d-48b0-8b3e-ae59a63d  AAATTGAAGATAAAGAT--AATGA----TTCCAATGAAAAT-G-AG-AT---GAATTTTGCAGAAAATCAAATTAT--CC [ 160]
44f1962a-fce6-436a-9ea0-ef4a969d  ATTGAA--GATAAAGAT--AATGA----TTCCAATGAAAAT-G-AG-AT--AGAATTTTGCAGAAAATCAAATTAT--CC [ 160]
4544809e-f4f9-4ff0-8142-5a2a2bc9  AGATTGAAGATAAAGAT--AATAG----TTCCAATGAAAAT-G-GG-ATCAAGAATTTTGCAGAAAATCAAATTAT--CC [ 160]
9faa5557-cdc1-466c-815f-fb18a15f  ATTGAAG--ATAAAGAT--AATGA----TTCCAATGAA--A-A-TG-AG--ATAATTTTGCAGAAAATCAAATTAT--CC [ 160]
ef1a361c-a80e-43f6-8e1f-aeafa681  ATTGAAG--ATAAAGAT--AATGA----TTCCGATGAAAAT-G-AG-AT--AGAATTTTGCAGAAAATCAAATTA-T-CC [ 160]

Cp_PARENT                         AAAAAATTCATGTGAAG--CACAGTGGATGTT--GGTAAGTTCT---GAGAAATATTATTAAAAAAAATTGGTGAG-ACA [ 240]
Ct_PARENT                         AAAAAATTCATGTGAAG--CACAGTGGATGTT--GGTAAGTTCT---GAGAAATATTATTAAAAAAAATTGGTGAG-ACA [ 240]
bdf46f5b-2833-4b26-892f-323dd4c3  AAAAAATTCATGTGAAG--CACAGTGGATGTT--GGTTAAGTTT--GAG-AAATATTATTAAAAAAAATTGGTGAG-ACA [ 240]
c40b234c-b999-47c3-93ca-016af61e  AAAAAATTCATGTGAAG--CACAGTGGATGTT--GGTAAGTTCT--GAG-AAATATTATTAAAAAAAATTGGTGAG-ACA [ 240]
0d8edcbc-b350-4d00-afbc-fd78a1f6  AAAAAATTCATGTGAAG--CACAGTGGATGTT--GGTAAGTTCT--GAG-AAATATTATTAAAAAA-ATTGGTGAG-ACA [ 240]
9cb07723-65ed-4c96-ac19-4234ea98  AAAAAATTCATGTGAAG--CACAGTGGATGTT--GGTAAGTTCT--GAG-AAATATTATTAAAAAAAATTGGTGAG-ACA [ 240]
ed18c856-4d5b-4622-9fe8-9b5740af  AAAAAATTCATGTGAAG--CACAGTGGATGTT--GGTAAGTTCA--G-----ATGTTTCTAAAAAAAATTGGTGAG-ACA [ 240]
c344612d-f873-4743-818c-65979638  AAAAAACCCATGTGAAG--CACAGTGGATGTT--GGTAAGTTCT--AGAGAAATCTA--TAAAAAAAATTGGTGAG-ACA [ 240]
baaaf62e-d0ee-4c49-9d83-0313d3df  AAAAAATTCATGTGAAG--CACAGTGGATGTT--GGTAAGTTCT--GAG-AAATATTATTAAAAAAAATTGGTGAG-ACA [ 240]
6771788a-69b9-4fd8-b4e0-073881fe  AAAAAATTCATGTGAAG--CACAGTGGATGTT--GGTAAGTTAG--GAAATATT---ATTAAAAAAAATTGGTGAG-ACA [ 240]
a6666a5a-6885-427a-9881-430dad33  AAAAAATTCATGTGAAG--CACAGTGGATGTT--GGTAAGTTCT--GAG-AAATATTATTAAAAAAAATTGGTGAG-ACA [ 240]
5be1c0db-7e16-4441-a2ab-f8107a5d  AAAAAATTCATGTGAAG--CACA-TGGATGTT--GGTAAGTTCT--GAG-AATGTTATTTAAAAAAAATTGGTGAG-ACA [ 240]
99a0bd03-c448-4605-9def-185e9f2d  AAAAAATTCATGTGAAG--CACAGTGGATGTT--GGTAAGTTCT--GAG-AAATATTATTAAAAAAAATTGGTGAG-ACA [ 240]
c37ace00-c1f8-46e8-bcdc-c3c33bcb  AAAAAATTCATGTGAAG--CACAGTGGATGTT--GGTAAGTTCT--GAG-AAATATTATTAAAAAAAATTGGTGAG-ACA [ 240]
cdc9d647-e2ac-442b-8435-6dd7b82f  AAAAAATTCATGTGAAG--CACAGTGGATGTT--GGTAACTTAT--AAG-AAATATTATTAAAAAAAATTGGTGAG-ACA [ 240]
bc10b467-59cf-4610-94fe-267549f8  AAAAAATTCATGTGAAG--CACAGTGGATGTT--GGTAAGTTCT--GAG-AAATATTATTAAAAAAAATTGGTGAG-ACA [ 240]
```

```
c8c3f0cb-c211-4877-8f07-1765b113  CAAAAATTCATGTGAAG--CACAGTGGATGTT--GGTAAGTTCT--GAG-AAATATTATTAAAAAAAATTGGTGAG-ACA [ 240]
720a0247-e02e-400b-901f-673d7527  AAAAAATTCATGTGAAG--CACAGTGGATGTT--GGTAAGTTAG--AAA-TATT---ATTAAAAAAAATTGGTGAG-ACA [ 240]
f3f8defb-2c03-44af-abc8-d94623f7  AAAAAATTCATGTGAAG--CACAGTGGATGTT--GGTAAGTTCTGAG--AAATATTA-TTAAAAAAAATTGGTGAG-ACA [ 240]
fc2a9b4c-2c5f-43ed-8ea7-f2af0743  AAAAAATTCATGTGAAG--CACAGTGGATGTT--GGTAAGTTAG-----AAATATTA-CTAAAAAAAATTGGTGAG-ACA [ 240]
477953c0-cb9c-400c-bf05-dfadb08e  AAAAAATTCATGTGAAGACCACAGTGGATGTTAGGGTAAGTTCT--GAGAATGTTTCTTAAAAAAAATTGGTCGAGCACA [ 240]
28c5afa3-4ebf-4ffa-a71d-2171913e  ACAAAATTCATGTGAAG--CACAGTGGATGTT--GGTAAGTTCT--GAGAAATATTA-TTAAAAAAAATTGGTGAG-ACA [ 240]
52a4f968-3394-459b-9ee3-10eab4a7  AAAAAATTCATGTGAAG--CACAGTGGATGTT--GGTAAGTTCTAGAAATATTATTC--------AATTTGGTGAG-ACA [ 240]
d36d5471-42a7-440b-bd5e-48d77156  AAAAAATTCATGTGAAG--CACGGTGGATATT--AATTGAGT-------CAATGTTATTTAAAAAAAATTGGTGAG-ACA [ 240]
f7886442-23fe-473a-9ead-a2b8a2a7  AAAAAATTCATGTGAAG--CACAGTGGATGTT--GGTAAGTTCT--GAGAAATATTA-TTAAAAAAAATTGGTGAG-ACA [ 240]
f4770134-2a4b-449d-a4f6-c99f3383  AAAAAATTCATGTGAAG--CACAGTGGATGTT--GGTAAGTTAT--TAGAAATATTA-TTAAAAAAAATTGGTGAG-ACA [ 240]
4dd53951-3b2d-42af-82ef-399bb712  AAAAAATTCATGTGAAG-ACACAGTGGATGTT--GGTAAGTT-------AAATATTA-TTAAAAAAAATTGGTGAG-ACA [ 240]
81ea3bac-8208-46c7-9546-bbd55b7c  AAAAAATTCATGTGAAG--CACAGTGGATGTT--GGTAAGTTCT--GAGAAATATTATCTAAAAAAAATTGGTGAG-ACA [ 240]
1e66a4e2-42fd-4fa9-b001-9bb7f263  AAAAAATTCATGTGAAG--CACAGTGGATGTT--GGTAAGTTCT--GAGAAATATTA-TTAAAAA-AATTGGTGAG-ACA [ 240]
b75fad9a-877e-4f52-968b-9fcbb7a4  AAAAAATTCATGTGAAG--CACAGTGGATGTT--GGTAAGTTCT--GAGAAATATTA-TTAAAAAAAATTGGTGAG-ACA [ 240]
f512b649-4f14-4eb6-b5a2-b0543441  AAAAAATTCATGTGAAG--CACAGTGGATGTT--GGTAAGTTCTGAGAAATATTCTACTTAAAAAAAATTGGTGAG-ACA [ 240]
5021e737-de02-42f2-920d-18868672  AAAAAATTCATGTGAAA--CTTAATGAGGATG--TTGGTAAGTTCTGAGAAATATTA-TTAAAAAAAATTGGTGAG-ACA [ 240]
a46adbf5-3ba1-48b8-a762-2403bda7  AAAAAATTCATGTGAAG--CACAGTGGATGTT--GGTAAGTTCT--GAGAAATATTA-TTAAAAAAAATTGGTGAG-ACA [ 240]
088a7e31-4c6b-4ff7-a19b-ff14531f  AAAAAATTCATGTGAAG--CACAGTGGATGTT--GGTAAGTTCTG--AGAAATATTA-TTAAAAAAAATTGGTGAG-ACA [ 240]
e82639d1-c451-4263-90a8-2b1ce6a1  AAAAAATTCATGTGAAG--CACAGTGGATGTT--GGTAAGTTATTAGAGAAATATTA-TTAAAAAAAATTGGTGAG-ACA [ 240]
3cca4221-4eee-44ed-9fb4-ebbac961  AAAAAATTCATGTGAAG--CACAGTGGATGTT--GGTAAGTTCTA-GAGAAATATTA-TTAAAAAAAATTGGTGAG-ACA [ 240]
d7a80796-8741-4878-b6ca-23aa66f7  AAAAAATTCATGTGAAG--CACAGTGGATGTT--GGTAACTTCT--GAGAAATATTA-TTAAAAAAAATTGGTGAG-ACA [ 240]
421e3eda-83a0-4ebd-972c-f24667ee  AAAAAATTCATGTGAAG--CACAGTGGATGTT--GGTAAGTTCT--GAGAAATATTA-TTAAAAAAAATTGGTGAG-ACA [ 240]
e6a39029-4e34-4457-b2a9-0cf6f207  AAAAAATTCATGTGAAG--CACAGTGGATGTT--GGTAAGTTCT--GAGAAATATTA-TTAAAAAAAATTGGTGAG-ACA [ 240]
a1507163-34c9-4b24-b21e-3b5dfc54  AAAAAGATTA-ATGAAG--C-CCGTGGATGTT--AACATT----------AGAAATGTATAAAAAAAATTGGTGAG-ACA [ 240]
51b01061-aaf5-4585-ba72-1a83671b  AAAAGATTAATATGAAG--CACAGTGGATGTT--GGTAAGTTCT--AGAGAAATATTATT-AAAAAAATTGGTGAG-ACA [ 240]
85703d1f-2c69-4541-ab66-acb7b838  AAAAAAGTCCTGTGAAG--CACAGTGGATGTT--GGCTAGTTAG--AGAAAT-ATTTATTAAAAAAAATTGGTGAG-ACA [ 240]
ed39aa3b-d227-4de1-92bd-5ace4420  AAAAAATTCATGTGAAG--CACAGTGGATGTT--GGTAAGTTCT---GAGAAATATTATT-AAAAAAATTGGTGAG-ACA [ 240]
40e2af31-875d-446e-90a0-6e16a943  AAAAAATTCATGTGAAG--CACAGTGGAGTAT--GCATCCTCTA------GAAATGTACTAAAAAAAATTGGTGAG-ACA [ 240]
873c76b4-8a29-4846-98df-cb5abcc9  AAAAAATTCATGTGAAG--CACAGTGGATGTT--GGTAAGTTCT--GAGAAATATTA-TTAAAAAAAATTGGTGAG-ACA [ 240]
59ef08a5-aa1d-4d15-ba6f-1a925971  AAAAAATTCATGTGAAG--CACAGTGGATGTT--GGTAAGTTCT--AAGAAATATTA-TTAAAAAAAATTGGTGAG-ACA [ 240]
e1cae4d8-4f39-4401-8d82-5c5edaf3  AAAAAATTCATGTGAAG--CACAGTGGATGTT--GGTAAGTTCT--GAGAAATATTATTAAA-AAAAATTGGTGAG-ACA [ 240]
5f47af75-253d-48b0-8b3e-ae59a63d  AAAAAATTCATGTGAAG--CACAGTGGATGTT--GGTAAGTCTT--GAGAAATATTCTAGAAAAAAAATTGGTGAG-ACA [ 240]
44f1962a-fce6-436a-9ea0-ef4a969d  AAAAAATTCATGTGAAG--CACAGTGGATGTT--GGTAAGTTCT--GAGAAATATTATTCG-------TTGGTTAG-ACA [ 240]
4544809e-f4f9-4ff0-8142-5a2a2bc9  AAAAAATTCATGTGAAG--CACGGTGGATGTT--GGTAAGTTCT--AGAAATGTA----TAAAAAAAATTGGTGAG-ACA [ 240]
9faa5557-cdc1-466c-815f-fb18a15f  AAAAAATTCATGTGAAG--CACAGTGGATGTT--GGTAAG-TTC---TAGAAATATTATTAAAAAAAATTGGTGAG-ACA [ 240]
ef1a361c-a80e-43f6-8e1f-aeafa681  AAAAAATTCATGTGAAG--CACAGTGGATGTT--GGTAAGTTAG--GA---AATATTATTAAAAAAAATTGGTGAG-ACA [ 240]

Cp_PARENT                         ATTTG-AATA---AT-TAAAGAG--AT--AATTTTTAGA----AAAATATAAATTATCTTTTTTTTT--TTTGTTGTTAA [ 320]
Ct_PARENT                         ATTTG-AAAA---AT-TAAAGAG--AT--AATTTTTAGA----AAAATATAAATTATCTTTTCTTTTTTGTTCTAGTTAA [ 320]
bdf46f5b-2833-4b26-892f-323dd4c3  ATTTG-AATA---AT-TAAAGAG--AT--AATTTTTAGA----AAAATATAAATTAT--C---TTTTTT-TTGTTGTTAA [ 320]
c40b234c-b999-47c3-93ca-016af61e  ATTTG-AATA---AT-TAAAGAG--AT--AATTTTAGAA----AAATATAAATTATC--T---TTTTTT-TTGTTGTTAA [ 320]
0d8edcbc-b350-4d00-afbc-fd78a1f6  ATTTG-AATA---AT-TAAAGAG--AT--AATTTTAGAA----AAATATAAATTACT------GTTTTT-TTGTTGTTAA [ 320]
9cb07723-65ed-4c96-ac19-4234ea98  ATTTG-AATA---AT-TAAAGAG--AT--AATTTTTAGA----AAAATATAATTATC----TTTTTTTT-TTGTTGTTAA [ 320]
ed18c856-4d5b-4622-9fe8-9b5740af  ATTTG-AATA---AT-TAAAGAG--AT--AATTTTTAGA----AAAATATAAATTAT--CTT----TTT--TGTTGTTAA [ 320]
c344612d-f873-4743-818c-65979638  ATTCG-AATC---AT-TAAAGAG--AT--AATTTTTAGA----AAAATATAAATTAT--CTTTTTTT---TTGTTGTTAA [ 320]
```

```
baaaf62e-d0ee-4c49-9d83-0313d3df  ATTTG-AATA---AT-TAAAGAG--AT--AATTTTTAGA----AAAATATAAATTAT--CTTTTTTT---TTGTTGTTAA [ 320]
6771788a-69b9-4fd8-b4e0-073881fe  ATTTG-AATA---AT-TAAAGAG--AT--AATTTT-AGA----AAAATATAAATTAT--CTTTCTTTTGTTGTTAA-T- [ 320]
a6666a5a-6885-427a-9881-430dad33  ATTTG-AATA---AT-TAAAGAG--AT--AATTTTTAGA----AAAATATAAATTA--------TTTTTTTTGTTGTTAA [ 320]
5be1c0db-7e16-4441-a2ab-f8107a5d  ATTTG-AATA---AT-TAAAGAG--AT--AATTTT-AGA----AAAATATAAATTAT--CTTTTTTGTT-GTT-----AA [ 320]
99a0bd03-c448-4605-9def-185e9f2d  ATTTG-AATA---AT-TAAAGAG--AT--AATTTTTAGA----AAATACTAAATTAT--CTTTTTTT---TTGTTGTTAA [ 320]
c37ace00-c1f8-46e8-bcdc-c3c33bcb  ATTTG-AATA---AT-TAAAGAG--AT--AA-TTTTAGA----AAAATATAAATTAT--CTTTT------TTGTTGTTAA [ 320]
cdc9d647-e2ac-442b-8435-6dd7b82f  ATTTG-AATA---AT-TAAAGAG--AT--AATTTTTAGA----AAAATATAATTTAT--CTTTTTTT---TTGTTGTTAA [ 320]
bc10b467-59cf-4610-94fe-267549f8  ATTTG-AATA---AT-TAAAGAG--AT--AATTTTTACA----AAATATTAAATTAT--CTTTTTTTG-TTGTTAA-T- [ 320]
c8c3f0cb-c211-4877-8f07-1765b113  ATTTG-AATA---AT-TAAAGAG--AT--AATTTTTAGA----AAAATATAAATTAT--CTTTTTTTAT-CTTTGTT-AA [ 320]
720a0247-e02e-400b-901f-673d7527  ATTTG-AATA---AT-TAAAGAG--AT--AATTTTTAGA----AAAATATAAATTAT--CTTTTTTTG-TTGTTAA-T- [ 320]
f3f8defb-2c03-44af-abc8-d94623f7  ATTTG-AATA---AT-TAAAGAG--AT--AATTTTTCAA----AAATATGA-----------TTCTTTTTTTGTTGTTAA [ 320]
fc2a9b4c-2c5f-43ed-8ea7-f2af0743  ATTTG-AATA---AT-TAAAGAG--AT--AATTTTTAGA----AAA-TATAAATTATC-TT---TTTTTTTTGTTGTTAA [ 320]
477953c0-cb9c-400c-bf05-dfadb08e  ATTTG-AATA---AT-TAAAGAG--AT--AATTTTTAGA----AAAATATAAATTATC-TT----TTTTTTTGTTGTTAA [ 320]
28c5afa3-4ebf-4ffa-a71d-2171913e  ATTTG-AATA---AT-TAAAGAG--AT--AATTTTTAGA----AAAATATAAATTAT-------CTTTTTTTGTTGTTAA [ 320]
52a4f968-3394-459b-9ee3-10eab4a7  ATTTG-AATA---AT-TAAAGAG--AT--AATTTTTAGA----AAAATATAAATTAT--------CTTTTTTGTTGTTAA [ 320]
d36d5471-42a7-440b-bd5e-48d77156  ATTTG-AATA---AT-TAAAGAG--AT--AATTTTTAGA----AAAATATAAATTATA-CTTTTTTTTTTTGTTGTTAA [ 320]
f7886442-23fe-473a-9ead-a2b8a2a7  ATTTG-AATA---AT-TAAAGAG--AT--AA-TTTTAGA----AAAATATAAATTAT--C----TTTTTTTTGTTGTTAA [ 320]
f4770134-2a4b-449d-a4f6-c99f3383  ATTTG-AATAATTAT-TAAAGAG--AT--AATTTTTAGA----AAAATATAAATTAT--CTTTTTTTTTTTGTTGTTAA [ 320]
4dd53951-3b2d-42af-82ef-399bb712  ATTTG-AATA---AT-TAAAGAG--AT--AAATTTTAGA----AAAATATAAATTAT--CT---TTTTTTTTGTTGTTAA [ 320]
81ea3bac-8208-46c7-9546-bbd55b7c  ATTTG-AATA---AT-TAAAGAG--AT--AATTTTTAGA----AAAATATAAATTAT--CT---TTTTTTTTGTTGTTAA [ 320]
1e66a4e2-42fd-4fa9-b001-9bb7f263  ATTTG-AATA---AT-TAAAGAG--AT--AATTTTTAGA----AAAATAAATTATCT--TT---TTTT---TGTTGTTAA [ 320]
b75fad9a-877e-4f52-968b-9fcbb7a4  ATTTG-AATA---AT-TAAAGAG--AT--AATTTTTAGA----AAAATATAAATTAT--CT---TTTTTTTTGTTGTTAA [ 320]
f512b649-4f14-4eb6-b5a2-b0543441  ATTTG-AATA---AT-TAAAGAG--AT--AATTTTTAG--------------AAAAAT--ATAATTATTTTTGTTGTTAA [ 320]
5021e737-de02-42f2-920d-18868672  ATTTG-AATA---AT-TAAAGAG--AT--AATTTTTAGA----AAAATATAAATTAT--CTTTTTTTTGTTGTTAATTTT [ 320]
a46adbf5-3ba1-48b8-a762-2403bda7  ATTTG-AATA---AT-TAAAGAG--AT--AATTTTTAGA----AAAATATAAATTAT--CTTTTTTTTTGTTGTTAATTT [ 320]
088a7e31-4c6b-4ff7-a19b-ff14531f  ATTTG-AATA---AT-TAAAGAG--AT--AATTTTTAGA----AATATAAATTATCT--TTTTTT-----TTGTTGTTAA [ 320]
e82639d1-c451-4263-90a8-2b1ce6a1  ATTTG-AATA---AT-TAAAGAG--AT--AATTTTTAGA----AAAATATAAATTAT--CTTTTTTTTTTTGTTGTTAA [ 320]
3cca4221-4eee-44ed-9fb4-ebbac961  ATTTG-AATA---AT-TAAAGAG--AT--AATTTTTAGA----AAAATATAAATTAT--C----TTTTTTTTTTTGTTGT [ 320]
d7a80796-8741-4878-b6ca-23aa66f7  ATTTG-AATA---AT-TAAAGAG--AT--AATTTTTAGA----AAAATATAAATTAT--C----TTTTTTTTTTTGTTGT [ 320]
421e3eda-83a0-4ebd-972c-f24667ee  ATTTG-AATA---AT-TAAAGAG--AT--AATTTTTAGA----AAAATATAAATTAT--C----TTTTTTTTGTTGTTAA [ 320]
e6a39029-4e34-4457-b2a9-0cf6f207  ATTTG-AATA---AT-TAAAGAG--AT--AATTTTTAGA----AAAATATAAATTAT--C----TTTTTTTTGTTGTTAA [ 320]
a1507163-34c9-4b24-b21e-3b5dfc54  ATTTG-AATA---AT-TAAAGAG--AT--AATTTTTAGA----AAAATATAAATTAT---CTTTTTTTT---GTTGTTAA [ 320]
51b01061-aaf5-4585-ba72-1a83671b  ATTTG-AATA---AT-TAAAGAGTCAT--AATTTTTAGA----AAAATATAAATTATC--TTTTTTTTG---TTGTTTAA [ 320]
85703d1f-2c69-4541-ab66-acb7b838  ATTTG-AATA---AT-TAAAGAG--AT--AATTTTTAGA----AAAATATAAATTAT---CTTTTTTTT---GTTGTTAA [ 320]
ed39aa3b-d227-4de1-92bd-5ace4420  ATTTG-AATA---AT-TAAAGAG--AT--AATTTTTAGA----AAAATATAAATTATC-TT----TTTG---TTGTTAA- [ 320]
40e2af31-875d-446e-90a0-6e16a943  ATTTG-AATA---AT-TAAAGAG--AT--AATTTTTAGA----AAAATATCATTCTCT-GT----TTTT---GTTGTTAA [ 320]
873c76b4-8a29-4846-98df-cb5abcc9  ATTTG-AATA---AT-TAAAGAG--AT--AATTTTTAGA----AAAA--TATAAATTA-TCTTTTTTTT--TGTTGTTAA [ 320]
59ef08a5-aa1d-4d15-ba6f-1a925971  ATTTG-AATA---AT-TAAAGAG--AT--AATTTTTAGA----AAAA--TATAAATTA-TCTTTTTTTT---GTTGTTAA [ 320]
e1cae4d8-4f39-4401-8d82-5c5edaf3  ATTTG-AATA---AT-TAAAGAG--AT--AATTTT--TAG---AAAAATATAAATTAT----CTTTTTT---GTTGTTAA [ 320]
5f47af75-253d-48b0-8b3e-ae59a63d  ATCTG-AATA---AT-TAAAGAG--AT--AATTTTTAGA----AAAATATAGAAATTA-TC---TTTTT---GTTGTTAA [ 320]
44f1962a-fce6-436a-9ea0-ef4a969d  ATTTG-AATA---AT-TAAAGAG--AT--AATTTTTAGA----AAAATATAAATTATC-TTTTTTTTTT---GTTGTTAA [ 320]
4544809e-f4f9-4ff0-8142-5a2a2bc9  ATTTG-AATA---AT-TAAAGAG--AT--AATTTTTAGA----AAAATCTAAACTATC-TTTTTTTTTT---GTTGTTAA [ 320]
9faa5557-cdc1-466c-815f-fb18a15f  ATTTG-AATA---AT-TAAAGAG--AT--AATTTTTAGA----AAAATATAAATTATCTTTTTTTTTTG-----TTGTTAA [ 320]
ef1a361c-a80e-43f6-8e1f-aeafa681  ATTTG-AATA---AT-TAAAGAG--AT--AATTTTTAGA----AAAATATAAATTAT--CTTTTTTTTG-TT---GTTAA [ 320]
```

```
Cp_PARENT                               TTTTTAA---TAATATTTACACAA--A----AATATAAATAAAT-ATATAAATACTTG---AAAACT------------T [ 400]
Ct_PARENT                               TTTTTAA---TAATATTTAAACAA--A----AATATAAATAAAT-ATATAAATACTTC---AAAAC-------------- [ 400]
bdf46f5b-2833-4b26-892f-323dd4c3        TTTTTAA---TAATATTTACACAA-A-----AATATAAATAAAT-ATATAAATAC-TCAAA-----AC--------TTTT [ 400]
c40b234c-b999-47c3-93ca-016af61e        TTTTTAA---TAATATTTACACAA-A-----AATATAAATAAAT-ATATAAATACTTGAAA-----AC--------TTTT [ 400]
0d8edcbc-b350-4d00-afbc-fd78a1f6        TTTTTAA---TAATATTTACACAA-A-----AATATAAATAAAT-ATATAAATACTTGAAA-----AC--------TTTT [ 400]
9cb07723-65ed-4c96-ac19-4234ea98        TTTTTAA---TAATATTTACACAA-A-----AATATAAATAAAT-ATATAAATACTTGAAA-----AC--------TTTT [ 400]
ed18c856-4d5b-4622-9fe8-9b5740af        TTTTTAA---TAATATTTACACAA-A-----AATATAAATAAAT-ATATAAATACTTGAAA-----AC--------TTTT [ 400]
c344612d-f873-4743-818c-65979638        TTTTTAA---TAATATTTACACAA-A-----AATATAAATAAAT-ATATAAATACTTGAAA-----AC---------TTT [ 400]
baaaf62e-d0ee-4c49-9d83-0313d3df        TTTTTAA---TAATATTTGCACAA-AG----AATATAAATAAAT-ATATAAATACTTGAAA-----AC---------TTT [ 400]
6771788a-69b9-4fd8-b4e0-073881fe        -TTTTAA---TAATATTTACACAA-A-----AATATAAATAAAT-ATATAAATACTTGAAA-----AC--------TTTT [ 400]
a6666a5a-6885-427a-9881-430dad33        TTTTTAA---TAATATTTACACAA-A-----AATATAAATAAAT-ATATAAATACTTGAAA-----A-----------CT [ 400]
5be1c0db-7e16-4441-a2ab-f8107a5d        TTTTTAA---TAATATTTACACAA-A-----AATATAAATAAAT-ATATAAATACTTGAAA-----AC--------TTTT [ 400]
99a0bd03-c448-4605-9def-185e9f2d        TTTTTAA---TAATATTTACACAA-A-----AATATAAATAAAT-ATATAAATACTTGAAA-----ACT--------TTT [ 400]
c37ace00-c1f8-46e8-bcdc-c3c33bcb        TTTTTAA---TAATATTTGCACAA-A-----AATATAAATAAAT-ATATAAATATCT-AAA-----ACT--------TTT [ 400]
cdc9d647-e2ac-442b-8435-6dd7b82f        TTTTTAA---TAATATTTACACAA-A-----AATATAAATAAAT-ATATAAATACTT-GAA-----AAC--------TTT [ 400]
bc10b467-59cf-4610-94fe-267549f8        -TTTTAA---TAATATTTACACAA-A-----AATATAAATAAAT-ATATAAATACTT---------GAAAACTTTTTT- [ 400]
c8c3f0cb-c211-4877-8f07-1765b113        TTTTTAA---TAATATTTACACAA-A-----AATATAAATAAAT-ATATAAA--TAC---------T-TGAAAACTTTT- [ 400]
720a0247-e02e-400b-901f-673d7527        -TTTTAA---TAATATTTACACAA-A-----AATATAAATAAAT-ATA-TAA--ATT---------TCAAAATCTTTTT- [ 400]
f3f8defb-2c03-44af-abc8-d94623f7        TTTTTAA---TAATATTTACACAA--A----AATATAAATAAAT-ATATAAATACTTGAA------AAC-------TTTT [ 400]
fc2a9b4c-2c5f-43ed-8ea7-f2af0743        TTTTTAA---TAATATTTACACAA--A----AATATAAATAAAT-ATATAAATACTTGAAA---------------ACTT [ 400]
477953c0-cb9c-400c-bf05-dfadb08e        TTTTTAA---TAATATTTACACAA--A----AATATAAATAAAT-ATATAAATACTTGAAA----AACT-------TTTT [ 400]
28c5afa3-4ebf-4ffa-a71d-2171913e        TTTTTAA---TAATATTTACAAAA--A------TATAAATAAAT-ATATAAATACTTGAAA-----AC----------TT [ 400]
52a4f968-3394-459b-9ee3-10eab4a7        TTTTTAA---TAATATTTACACAA--A----AATATAAATAAAT-ATATAAATAC-TTGAA-----AAC-------TTTT [ 400]
d36d5471-42a7-440b-bd5e-48d77156        TTTTTAA---TAATATTTACACAA--A----AATATAAATAAAT-ATATAAA---ATACAA-----AAC-------TTTT [ 400]
f7886442-23fe-473a-9ead-a2b8a2a7        TTTTTAA---TAATATTTACACAA--A----AATATAAATAAAT-ATATAAATACTTGAAA-----AC---------TTT [ 400]
f4770134-2a4b-449d-a4f6-c99f3383        TTTTTAA---TAATATTTACACAA--A----AATATAAATAAAT-ATATAAATAC-TTGAA-----AAC-------TTTT [ 400]
4dd53951-3b2d-42af-82ef-399bb712        TTTTTAA---TAATATTTACACAA--A----AATATAAATAAAT-ATATAAATACTTGAAA-----AC---------TTT [ 400]
81ea3bac-8208-46c7-9546-bbd55b7c        TTTTTAA---TAATATTTACACAA--A----AATATAAATAAAT-ATATAAATACTTGAAA-----ACT-------TTTT [ 400]
1e66a4e2-42fd-4fa9-b001-9bb7f263        TTTTTAA---TAATCCTTGCACAA--A----AATATAAATAAAT-ATATAAATACTTGAAA-----ACT-------TTTT [ 400]
b75fad9a-877e-4f52-968b-9fcbb7a4        TTTTTAA---TAATATTTACACAA--AGAAAAATATAAATAAAT-ATATAAATACTTGAAA-----ACT----------T [ 400]
f512b649-4f14-4eb6-b5a2-b0543441        TTTTTAA---TAATATTTACACAA--A----AATATAAATAAAT-ATATAAATACTTGAAA-----ACT-------TTTT [ 400]
5021e737-de02-42f2-920d-18868672        T----AA---TAATATTTACACAA--A----AATATAAATAAAT-ATATAAATAC--TTGA-----AAA-------CTTT [ 400]
a46adbf5-3ba1-48b8-a762-2403bda7        TTAATAA------TATTTACACAA--A----AATATAAATAAAT-ATATAAATAC-TTGAA-----AAC-------TTTT [ 400]
088a7e31-4c6b-4ff7-a19b-ff14531f        TTTTTAA------TAATATTTACA--C----AAAGTAAATAAAT-ATATAAATAC-TTGAA-----AAC-------TTTT [ 400]
e82639d1-c451-4263-90a8-2b1ce6a1        TTTTTAATAAT---ATTTACACAA--A----AATATAAATAAAT-ATATAAATAC-TTGAA-----AAC-------TTTT [ 400]
3cca4221-4eee-44ed-9fb4-ebbac961        TAATTTTTAATAATATTTACACAA--A----AATATAAATAAAT-ATATAAATAC-TTGAA-----AA--------TTTT [ 400]
d7a80796-8741-4878-b6ca-23aa66f7        TAATTTTTAATAATATTT-ACAGA--A----AATATAAATAAAT-ATATAAATAC-TTGAA-----AAC----------T [ 400]
421e3eda-83a0-4ebd-972c-f24667ee        TTTTTCA---T--GAGTA-CACAA--A----AATATAAATAAAT-ATATAAATAC-TTGAA-----AAC-------TTTT [ 400]
e6a39029-4e34-4457-b2a9-0cf6f207        TTTTTAATAAT--ATTTA-CACAA--A----AATATAAATAAAT-ATATAAATAC-TTGAA-----AAC-------TTTT [ 400]
a1507163-34c9-4b24-b21e-3b5dfc54        TTTTTAA---TAATATTTACACAA--A----AATATAAATAAAT-ATATAAATACTTG-AA----AAC------------ [ 400]
51b01061-aaf5-4585-ba72-1a83671b        TTTTTAA---TAATATTTACAC-A--A----AATATAAATAAAT-ATATAAATACTTG-AA----AAC---------TTT [ 400]
85703d1f-2c69-4541-ab66-acb7b838        TTTTTAA---TAATATTTACAC-A--A----AATATAAATAAAT-ATATAAATAC-TG-AA----AAC---------TTT [ 400]
ed39aa3b-d227-4de1-92bd-5ace4420        -TTTTAA---TAATATTTACACAA--A----AATATAAATAAAT-ATATAAATACTTG-AA----AGC--------TTTT [ 400]
40e2af31-875d-446e-90a0-6e16a943        TTTTTAA---TAATATTTACACAA--A----AATATAAATAAAT-ATATAAATACTTG-AA----AAC--------TTTT [ 400]
873c76b4-8a29-4846-98df-cb5abcc9        TTTTTAA---TAATATTTACACAA--A----AATATAAATAAAT-ATATAAATACTTCAAA-----AC--------TTTT [ 400]
```

```
59ef08a5-aa1d-4d15-ba6f-1a925971  TTTTTAA---TAATATTTACACAA--A----AATATAAATAAAT-ATATAAATACTTGGAA----AAC--------TTTT [ 400]
e1cae4d8-4f39-4401-8d82-5c5edaf3  TTTTTAA---TAATATTTACACAA--A----AATATAAATAAAT-ATATAAATACTTG-AA----AAC--------TTTT [ 400]
5f47af75-253d-48b0-8b3e-ae59a63d  TTTTTAA---TCAT------ACAA--A----AATATAAATAAAT-ATAAAT-----CT-TA----AAA--------ATTT [ 400]
44f1962a-fce6-436a-9ea0-ef4a969d  TTTTTAA---TAATATTTACACAA--A----AATATAAATAAAT-ATATAAATACTTG-AA----AAC--------TTTT [ 400]
4544809e-f4f9-4ff0-8142-5a2a2bc9  TTTTTAA---TAATATTTACACAA--A----AT-ATAAATAAAT-ATATAA-----AT-CT----AAA--------ACTT [ 400]
9faa5557-cdc1-466c-815f-fb18a15f  TTTTTAA---TAATATTTACACAA--A----AATATAAATAAAT-ATATAAAATATGA---AAACTT------------- [ 400]
ef1a361c-a80e-43f6-8e1f-aeafa681  TTTTTAA---TAATATTTACACAAAA-----ATGTAAAATAAAT-ATATAAATACTTGAAA-----AC--------TTTT [ 400]

Cp_PARENT                         TTTTTTTATTATTCTAGTTGTATTCCTG-TATA-AC--TTCAAT-GGAAGA-TCATCCAAGCGAGGCT----AGTCT--- [ 480]
Ct_PARENT                         TTTTTTTAAATATTCTAGTTGTATTCCTG-TATA-AC--TTCAAT-GGAAGA-TCATCCAAGCGAGGCT----AGTCT--- [ 480]
bdf46f5b-2833-4b26-892f-323dd4c3  TTTTT--ATTATTCATATTGTATAG--------A-C--TTCAAT-GGAAGATAGT-GCAAGCGCG-GCTAGTC--TT--- [ 480]
c40b234c-b999-47c3-93ca-016af61e  TTTTT--ATTATTCTAGTTGTATTCCTG-TATAA-C--TTCAAT-GGAAAATCAT-CCAAGCGAG-GCTAGTC--TT--- [ 480]
0d8edcbc-b350-4d00-afbc-fd78a1f6  TTTTT--ATTATTCTAGTTGTATTCCTG-TATAA-C--TTAAAT-GGAAGATGAC-AGACAGACT-ATCTTTA------- [ 480]
9cb07723-65ed-4c96-ac19-4234ea98  TTTTT--ATTATTCTAGTTGTATTCCTG-TATAA-C--TTCAAT-GGAAGATCAT-CCAAGCGAG-GCTAGTC--TT--- [ 480]
ed18c856-4d5b-4622-9fe8-9b5740af  TTTTT--ATTATTCTAGTATCTGTCCT--TATAA-C--TTCAAT-GGAAGATCATCCCAAGCGAG-GCTAGTC--TT--- [ 480]
c344612d-f873-4743-818c-65979638  TTTTT--ATTATTCTAGTTGTATTCCTG-TATAA-C--TTCAAT-GGAAGATCAT-CCAAGCGAG-GCTAGTC--TT--- [ 480]
baaaf62e-d0ee-4c49-9d83-0313d3df  TTTTT--ATTATTCTAGTTGTATTCCTG-TATAA-C--TTCAAT-GGAAGATCAT-CCAAGCGAG-GCTAGTC--TT--- [ 480]
6771788a-69b9-4fd8-b4e0-073881fe  TTTTT--ATTATTCTAGTTGTA-TCCTGTATAAG-T--TTCAAT-GGAAGATCAT-CCAAGCGAG-GCTAGTC--TT--- [ 480]
a6666a5a-6885-427a-9881-430dad33  TTTTT--TTTATTATTCGTGTATTCCTG-TATAA-C--TCAAAT-GGAAGATCAT-CCAAGCGAG-GCTAGTC--TT--- [ 480]
5be1c0db-7e16-4441-a2ab-f8107a5d  TTTTT--ATTATTCTAGTTGTATTCCTATTATAA-C--TTCAAT-GGAAGATCAT-CCAAGCGAG-GCTAGTC--TT--- [ 480]
99a0bd03-c448-4605-9def-185e9f2d  TTTTT--ATTATTCTAGTTGTATTCCTG-TATAA-C--TTCAAT-GGAAGATCAT-CCAAGCGAGGCTAGT-C--TT--- [ 480]
c37ace00-c1f8-46e8-bcdc-c3c33bcb  TTTTT--ATTATTCTAGTTGTATTCCTG-TATAA-C--TTCAAT-GGAAGATCAT-CCAAGCGAG-GCTAGTC--TT--- [ 480]
cdc9d647-e2ac-442b-8435-6dd7b82f  TTTTT--ATTATTCTAGTTGTATTCCTG-TATAA-C--TTCAAT-GGAAGATCAT-CCAAGCGAG-GCTAGTC--TT--- [ 480]
bc10b467-59cf-4610-94fe-267549f8  ---TT--ATTATTCTAGTTGTATTCCTG-TATAA-C--TTCAAT-GGAAGATCAT-CCAAGCGAG-GCTAGTC--TT--- [ 480]
c8c3f0cb-c211-4877-8f07-1765b113  ---TT--TTTATTCTAGTTGTATTCCTG-TATAA-C--TTCAAT-GGAAGATCAT-CCAAGCAAG-GCTAGTC--TT--- [ 480]
720a0247-e02e-400b-901f-673d7527  ---TT--ATTATTCTAGTTGTATTCCTG-TATAA-C--TTCAAT-GGAAGATCAT-CCAAGCGAG-GCTAGTC--TT--- [ 480]
f3f8defb-2c03-44af-abc8-d94623f7  TTTTT--ATTATTCTAGTTGTATTCCTG-TATAA-C--TTCAAT-GGAAGATCAT-CCAAGCGAGGCT-AGTC--TT--- [ 480]
fc2a9b4c-2c5f-43ed-8ea7-f2af0743  TTTTT--ATTATTCTAGTTGTATTCCTG-TATAA-C--TTCAAT-GGAAGATCAT-CCAAGCAAGGCT-AGTC--TT--- [ 480]
477953c0-cb9c-400c-bf05-dfadb08e  TTTTT--ATTATTCTAGTTGTATTCCTG-TATAA-C--TTCAAT-GGAAGATAAT-TA-AGCGAGGCT-AGTC--TT--- [ 480]
28c5afa3-4ebf-4ffa-a71d-2171913e  TTTTT--ACTATTCTAGTTGTATTCCTG-TATAA-C--TTCAAT-GGAAGATCAT-CCAAGCGAGGCT-AGTC--T---- [ 480]
52a4f968-3394-459b-9ee3-10eab4a7  TTTTT--ATTATTCTAGTTGTATTCCTG-TATAA-C--TTCAAT-GGAAGATCAT-CCAAGCGAGGCT-AGTC--TT--- [ 480]
d36d5471-42a7-440b-bd5e-48d77156  TTTTT--ATTATTCTAGTTGTATTCCTG-TATAA-C--TTCAAT-GGAAGATCAT-CCAAGCGAGGCT-AGTC--TT--- [ 480]
f7886442-23fe-473a-9ead-a2b8a2a7  TTTTT--TATCTGTGAGTTGTATTCCTG-TATAA-C--TTCAAT-GGAAGATCAT-CCAAGCAAGGCTAAGTC--TT--- [ 480]
f4770134-2a4b-449d-a4f6-c99f3383  TTTTT--ATTATTCTAGTTGTATTCCTG-TATAA-C--TTCAAT-AGAA-ATCAT-CCAAGCGAGGCT-AGTC--TT--- [ 480]
4dd53951-3b2d-42af-82ef-399bb712  TTTTT--ATTATTCTAGTTGTATTCCTG-TATAA-C--TTCAAT-GGAAGATAGT-CC-AGCGAGGCT-AGTC--TT--- [ 480]
81ea3bac-8208-46c7-9546-bbd55b7c  TTCT------CTTCTAGTTGTATTCCTG-TATAA-C--TTCAAT-GGAAGATCAT-CCAAGCGAGGCT-AGTC--TT--- [ 480]
1e66a4e2-42fd-4fa9-b001-9bb7f263  TTTTT--ATTATTCTAGTTGTATTCCTG-TATAA-C--TTCAAT-GGAAGATCAT-CCAAGCGAGGCT-AGTC--TT--- [ 480]
b75fad9a-877e-4f52-968b-9fcbb7a4  TTTTT--TTTATTCTAGTTGTATTCCTG-TATAA-C--TTCAAT-GGAAGATCAT-CCAAGCGAGGCT-AGTC--TT--- [ 480]
f512b649-4f14-4eb6-b5a2-b0543441  TTTTT--ATTATTCTAGTTGTATTCCTG-TATAA-C--TTCAAT-GGAAGATCAT-CCAAGCGAGGCT-AGTC--TT--- [ 480]
5021e737-de02-42f2-920d-18868672  TTTTT--ATTATTATCATTGTATTCCTG-TATAA-C--TTCAAT-GGAAGATCAT-CCAAGCGAGGCT-AAGT--CT--- [ 480]
a46adbf5-3ba1-48b8-a762-2403bda7  TTTTT--ATTATTCTAGTTGTATTCCTG-TATAA-C--TTCAAT-GGAAGATCAT-CCAAGCAGAGCT-AGTC--TT--- [ 480]
088a7e31-4c6b-4ff7-a19b-ff14531f  TTTTT--ATTATTCTAGTTGTATTCCTG-TATAA-C--TTCAAT-GGAAGATCAT-CCAAGCGAGGCT-AGTC--TT--- [ 480]
e82639d1-c451-4263-90a8-2b1ce6a1  TTTTT--ATTATTCTAGTTGTATTCCTG-TATAA-C--TTCAAT-GGAAGATCAT-CCAAGCGAGGCT-AGTC--TT--- [ 480]
3cca4221-4eee-44ed-9fb4-ebbac961  CTTTT--ATTATTCTAGTTGTATTCCTG-TATAA-C--TTCAAT-GGAAGATCAT-CCAAGCGAGGCT-AGTC--TT--- [ 480]
d7a80796-8741-4878-b6ca-23aa66f7  TTTTT--ATTATTCTAGTTGTATTCCTG-TATAA-C--TTCAAT-GGAAGATCAT-CCAAGCGAGGCT-AGTC--TT--- [ 480]
```

```
421e3eda-83a0-4ebd-972c-f24667ee  TTTTT--ATTATTCTAGTTGTATTCCTG-TATAA-C--TTCAAT-GGAAGATCAT-CCAAGCGAGGCT-AGTC--TT--- [ 480]
e6a39029-4e34-4457-b2a9-0cf6f207  TTTTT--ATTATTCTAGTTGTATTCCTG-TATAA-C--TTCAAT-GGAAGATCAT-CCAAGCGAGGCT-AGTC--TT--- [ 480]
a1507163-34c9-4b24-b21e-3b5dfc54  TTTTTTTTTTATTACTCATCTATTCCTG-TATA-AC--TTCAAT-GGAAGATCAT-CCAAGCAAGGCT----AG-TC-T- [ 480]
51b01061-aaf5-4585-ba72-1a83671b  TTTTTATTATTC--TCGTTGTATTCC---TATA-AC--TTCAAT-GGAAGATCAT-CCAAGCGAGGCT----AGTCT-T- [ 480]
85703d1f-2c69-4541-ab66-acb7b838  TTTTTATTATTC--TAGTTGTATTCCTG-TATA-AC--TTCAAT-GGAAGATCAT-CCAAGCGAGGCT----AGTCT-T- [ 480]
ed39aa3b-d227-4de1-92bd-5ace4420  TTATTATT-----CTAGTTGTATTCCTG-TATA-AC--TTCAATGGAAGATAATT-AAGACGAGGCTA----GT-CT-T- [ 480]
40e2af31-875d-446e-90a0-6e16a943  TTTTTATTATTCTCTCATTGTATTCCTG-TATA-AC--TTCAATGGGAAGATCAT-CCCAGCA-AACG----TC-CT-C- [ 480]
873c76b4-8a29-4846-98df-cb5abcc9  TTTTT--ATTATTCTAGTTGTATTCCTG-TATA-AC--TTCAAT-GGAAGATCAT-CCAAGCGAGGCT----AGTCT-T- [ 480]
59ef08a5-aa1d-4d15-ba6f-1a925971  TTTTT--ATTATTCTAGTTGTATTCCTG-TATA-ACCTTTCAAT-GGAAGATCAT-CCAAGCGAGGCT----AGTCT-T- [ 480]
e1cae4d8-4f39-4401-8d82-5c5edaf3  TTTTTTATTCTTACTAGTTGTATTCCTG-TATA-AC--TTGGAT-GGAAGATCAT-CCAAGCGAGGCT----AGTCT-T- [ 480]
5f47af75-253d-48b0-8b3e-ae59a63d  TTTTT--TATTCTCTCATTGC-ATGCTG-TATA-GC--TCAATG-GGAGGATCAT-CCAAGCAAGGCT----AGTCT-T- [ 480]
44f1962a-fce6-436a-9ea0-ef4a969d  TTTTT--CTT-CTCTAGTTGTATTCCTG-TATA-AC--TTAAAT-GGAAT-TCAT-CCAAGCGAGGCT----AGTCTCT- [ 480]
4544809e-f4f9-4ff0-8142-5a2a2bc9  TTTTT--CTCTCTCTAGTTGTATTCCTG-TATA-AC--TTCAAT-GGAAGATCAT-CCAAGCGAGGCT----AGTCT-T- [ 480]
9faa5557-cdc1-466c-815f-fb18a15f  --TTTTTTATTCTCTAGTTGTATCCTTG-TATA-AC--TTCAAT-GGAAGA-TCATCCAAGCGAGGCT----AGTCT--- [ 480]
ef1a361c-a80e-43f6-8e1f-aeafa681  TTTTT--ATTATTCTAGTTGTATTCCTG-TATAA-C--TTCAAT-GGAAGATCAT-CCAAGCGAG-GCTAGTC--TT--- [ 480]

Cp_PARENT                         TAAA--TCTGCAA-AAGATTTTATTAGAAATATTC------------C-AGATC-AATGTAAA---TTTACTCAAATAGA [ 560]
Ct_PARENT                         TAAA--TCTGCAA-AAGATTTTATTAGAAATATTC------------C-AGATC-AATGTAAA---TTTACTCAAATAGA [ 560]
bdf46f5b-2833-4b26-892f-323dd4c3  AAAT--CT-GCAA-AAAG--GTTTCTTA--------AAATGTC----T-AATAC-AATGTAAA---TTTACTCA------ [ 560]
c40b234c-b999-47c3-93ca-016af61e  AATC--TT-GCAAA-AGA--TTT-------TATTCAAAATGTC----C-AGATG-AATGTAAA---TTTACTCAAATAG- [ 560]
0d8edcbc-b350-4d00-afbc-fd78a1f6  -ATA-----CTCC-AAAG--AGTTTATTC------AAATATTG----C-AGATC-AATGTAAA---TTTACCTCAATAG- [ 560]
9cb07723-65ed-4c96-ac19-4234ea98  AAAT--CT-GCAA-AAGA--TTTTATTAG------AAATATTC----C-AGATC-AATGTAAA---TTTACTCAAATAG- [ 560]
ed18c856-4d5b-4622-9fe8-9b5740af  AAAT--CT-GCAA-AAGA--TTTTATTCG-------AAATATC----A-GAATA-AATATAAA---TTTACTCAAATAG- [ 560]
c344612d-f873-4743-818c-65979638  AAAT--CT-GCAA-AAGA--TTTTATTGA-------AA-----TCTGC-AGATC-AATGTAAA---TTTATCAAAATAG- [ 560]
baaaf62e-d0ee-4c49-9d83-0313d3df  AAAT--CT-GCAA-AAGA--TTTTATTAG-------AAATATTC---C-AGATC-AATGTAAA---TTTACTGGATGGA- [ 560]
6771788a-69b9-4fd8-b4e0-073881fe  AAAT--CT-GCAA-AAGA--TTTTATTAG-------AAATATTC---C-AGATC-AATGTAAA---TTGCCCCTCAATAG [ 560]
a6666a5a-6885-427a-9881-430dad33  AAAT--CT-GCAA-AAGA--TTTTATTAG-------AAATATTC---C-AGATC-AATGTAAA---TTTACTCAAATAG- [ 560]
5be1c0db-7e16-4441-a2ab-f8107a5d  AAAT--CT-GCAA-AAGA--TTTTATTAG-------AAATATTC---C-AGATC-AATGTAAA---TTTACTCAAATAG- [ 560]
99a0bd03-c448-4605-9def-185e9f2d  AAAT--ATCGCAA-AAGA--TTTTATTAG-------AAATATTC---C-AGATC-AATGTAAA---TTTACTCAAATAGA [ 560]
c37ace00-c1f8-46e8-bcdc-c3c33bcb  AAAT--CT-GCAA-AAGA--TTTTATTAG-------AA------TGTT-AGATC-AATGTAAA---TTTACTCAAATAG- [ 560]
cdc9d647-e2ac-442b-8435-6dd7b82f  AAAT--CT-GCAA-AAGA--TTTTATTCG-------AAATATT---GC-GGATC-AATGTAAA---TTTACTCAAATAG- [ 560]
bc10b467-59cf-4610-94fe-267549f8  AAAT--CT-GCAA-AAGA--TTTTATTAG-------AAATATTC---C-AGAAC-AATGTAAATTTACTCAATGA-AGA- [ 560]
c8c3f0cb-c211-4877-8f07-1765b113  AAAT--CT-GCAA-AAGA--TTTTATTCA-------AATA-TTC---C-AGATC-AATGTAAATTTACTCAAATAGAAG- [ 560]
720a0247-e02e-400b-901f-673d7527  AAAT--CT-GCAA-AAGA--TTTTATTAG-------AAATATTC---C-AGATC-AATGTCAAA--TTTACTCAAATAG- [ 560]
f3f8defb-2c03-44af-abc8-d94623f7  AAAT--CT-GCAA-AAGATTTTATTAGA--------AATA----TTCC-AGATC-AATGTAAA---TTTACTCAAA-TAG [ 560]
fc2a9b4c-2c5f-43ed-8ea7-f2af0743  AAAT--CT-GCAA-AAGA--TTTTATTAG------AAATA----TTCC-AGATC-AATGTAAA---TTTACTCAAA-TAG [ 560]
477953c0-cb9c-400c-bf05-dfadb08e  AAAT--CT-GCAA-AAGA--TTTTGTAGA------ATGTT-------T-AGATC-AATGTAAA---TTTACTCAAA-TAG [ 560]
28c5afa3-4ebf-4ffa-a71d-2171913e  AAAT--CT-GCAA-AAGA--TTTTATTAG------AAATA----TTCC-AATAA-AATCTAAA---TTTACTCAAA-TCA [ 560]
52a4f968-3394-459b-9ee3-10eab4a7  AAAT--CT-GCAA-AAGA--TTTTATTAG------AAATA----TTCC-AGATC-AATGTAAA---TTTACCTCAAATAG [ 560]
d36d5471-42a7-440b-bd5e-48d77156  AAAT--CT-GCAA-AAGATTTTATTAGAG------AAATA----TTGC-AGATC-AATGTAAA---TTTACTCAAA-TAG [ 560]
f7886442-23fe-473a-9ead-a2b8a2a7  AAAT--CT-GCAA-AAGA--TTTTATTAG------AAATA-TTC---C-AGATC-AATGTAAA---TTTACTCAAA-TAG [ 560]
f4770134-2a4b-449d-a4f6-c99f3383  AAAT--CT-GCAA-AAGA--TTT------------ATAGAAATATTCC-AGATC-AATGTAAA---TTTACTCAAA-TAG [ 560]
4dd53951-3b2d-42af-82ef-399bb712  AAAT--CT-CCAA-AGAA--TTTTCTCAA------AATGT-------A-CAATC-AATGTAAT---TTACCTCAAA-TAG [ 560]
81ea3bac-8208-46c7-9546-bbd55b7c  AAAT--AT-GCAA-AAGA--TTTTATTAG------AAATA-TTC---C-AGATC-AATGTAAA---TTTACTCAAA-TAG [ 560]
1e66a4e2-42fd-4fa9-b001-9bb7f263  AAAT--CT-GCAA-AAGA--TTTTATTAG------AAATATTCC---C-AGATC-AATGTAAA---TTTACTCAA--TAG [ 560]
```

```
b75fad9a-877e-4f52-968b-9fcbb7a4  AAAT--CT-GCAA-AAGA--TTTTATTAG------AAATATTCC---A-GATCA-ATGTAAAT---TTACTCAAA--TAG [ 560]
f512b649-4f14-4eb6-b5a2-b0543441  AAAT--CT-GCAA-AAGA--TTTTATTCA------AATATTC-----C-AGATC-AATGTAAA---TTTACTCAAA-TAG [ 560]
5021e737-de02-42f2-920d-18868672  TAAA--AT-ATCC-AAAA--GATTTTCCT------CAATC------TT-AGATC-AATGTAAA---TTTACTCAAA-TAG [ 560]
a46adbf5-3ba1-48b8-a762-2403bda7  AAAT--CT-GCAA-AAGA--TTTTGTAGG------AAAAT-ATTG--C-AGATC-AATGTAAA---TTTACTCAAA-TGA [ 560]
088a7e31-4c6b-4ff7-a19b-ff14531f  AAAT--CT-GCAA-AAGA--TTTTATTTC------AGAAATATTC--C-AGATC-AATGTAAA---TTTACTCAAA-TAG [ 560]
e82639d1-c451-4263-90a8-2b1ce6a1  GAAT--CT-GCAA-AAGA--TTTTATTAG------AAATATTTG---C-AGATC-AATATTGA---T---CCTCAA-TAG [ 560]
3cca4221-4eee-44ed-9fb4-ebbac961  AAAT--CT-GCAA-AAGA--TTTTCTCAA------ATAT--TCC---G-AGATC-AATGTAAA---TTTACTCAAA-TCG [ 560]
d7a80796-8741-4878-b6ca-23aa66f7  AAAT--CT-GCAA-AAGA--TTTTATTAG------AAATA-TTC---C-AGATC-AATGTAAA---TTTACTCAAA-TAG [ 560]
421e3eda-83a0-4ebd-972c-f24667ee  AAAT--CT-GCAA-AAGA--TTTTATTAG------AAATA-TTC---C-AGATC-AATGTAAA---TTTACTCAA--TAG [ 560]
e6a39029-4e34-4457-b2a9-0cf6f207  AAAT--CT-GCAA-AAGA--TTTTCTAGA------AATAT-TCC---A-GATCA-ATGTAAAT---TTACTCAAA--TAG [ 560]
a1507163-34c9-4b24-b21e-3b5dfc54  AAAT--CT-CCAA-AAGATTTTACTAGAAAATGTC------------C-AGATC-AATGTAAA---TTTATCCAAATGGG [ 560]
51b01061-aaf5-4585-ba72-1a83671b  AAAT--CT-GCAA-AAGATTGTTG---AATATCTC------------C-AGATC-AATGTAAA---TTTACTCAAATAGA [ 560]
85703d1f-2c69-4541-ab66-acb7b838  AAAT--CT-GCAA-AAGATTTTATTAGAAATATTC------------C-AGATC-AATGTAAA---TTTACTCAAATAGA [ 560]
ed39aa3b-d227-4de1-92bd-5ace4420  AAAT--CT-GCAA-AAGATTTTATTAGAAATATTC------------C-AGATC-AATGTAAA---TTTACTCAAATCGA [ 560]
40e2af31-875d-446e-90a0-6e16a943  AAAT--CT-GCAA-AAGATTTTATTAGAAATATTC------------C-AGATC-AATGTAAA---TTTACTCAATCAAA [ 560]
873c76b4-8a29-4846-98df-cb5abcc9  AAAT--CT-GCAA-AAGATTTTATTAGAAATATTC------------C-AGATC-AATGTAAA---TTTACTCAAATAGA [ 560]
59ef08a5-aa1d-4d15-ba6f-1a925971  AAAT--CT-GCAA-AAGATTTTATTAG-AATGTTC------------C-AGATC-AATGTAAA---TTTACTCAAATAGA [ 560]
e1cae4d8-4f39-4401-8d82-5c5edaf3  AAAT--CT-GCAA-AAGATTTTATTAGAAATATTC------------C-ATCAA-T--GTAAA--------TTTACTCAA [ 560]
5f47af75-253d-48b0-8b3e-ae59a63d  AA-T--CT-GCAA-AAGATTTTATT-GAGAATGTT------------T-AGATA-AATGTAAA---TTTACTCAAATAGA [ 560]
44f1962a-fce6-436a-9ea0-ef4a969d  AAAT--CT-GCAA-AAGATTTTATT-CAAATATTC------------C-AGATC-AATGTAAA---TTTACTCAAATAGA [ 560]
4544809e-f4f9-4ff0-8142-5a2a2bc9  AATC--TT-GCAA-AAGAT--TTTGTCAAATATTC------------C-AGATC-AATGTAAA---TTTACT-AAATAGA [ 560]
9faa5557-cdc1-466c-815f-fb18a15f  AAAT--CTGCAAA-AGA-T---TTTATTAGAATGT------------T-AGATC-AATGTAAA---TTTTATTCAAATAA [ 560]
ef1a361c-a80e-43f6-8e1f-aeafa681  AAAT--CT-GCAA-AAGA--TTTTATTAG------AAATATTC----C-AGATC-AATGTAAA---TTTACTCAA-TAG- [ 560]

Cp_PARENT                         AG-ATACATTT-TC----AGAAGGAATAAAAGA-TTTT--GATAACG-CTA-TTTCCA-CT-GCAA---A--C-TCA-AA [ 640]
Ct_PARENT                         AG-ATACATTT-TC----AGAAGGAATAAAAGA-TTTT--GATAACG-CTA-TTTCCA-CT-GCAA---A--C-TCA-AA [ 640]
bdf46f5b-2833-4b26-892f-323dd4c3  ATCAAAGATAGTTTAGAAGGAAT----AAAAGA-TTTT--GATAACG-CTA-TTTCCA-CT-GCAA---A--C-TCA-AA [ 640]
c40b234c-b999-47c3-93ca-016af61e  AAGATACAT--TTTCAAAGGAAT----AAAAGA-TTTT--GATAACG-CTA-TTTCCA-CT-GCAA---A--C-TCA-AA [ 640]
0d8edcbc-b350-4d00-afbc-fd78a1f6  AAGATACATTTTCAG-AAGGAAT----AAAAGA-TTTT--GATAACG-CTA-TTTCCA-CT-GCAA---A--C-TCA-AA [ 640]
9cb07723-65ed-4c96-ac19-4234ea98  AAGATACATTTTCAG-AAGGAAT----AAAAGA-TTTT--GATAACG-CTA-TTTCCA-CT-GCAA---A--C-TCA-AA [ 640]
ed18c856-4d5b-4622-9fe8-9b5740af  AAGATACATTTTCCAGAAGGGAT----GGAAAA-GAT----ATAACG-CTA-TTTCCA-CT-GCAA---A--C-TCAGAA [ 640]
c344612d-f873-4743-818c-65979638  AAGATACATTT-TCAGAAGGAAT----AAAAGA-TTTT--GATAACG-CTA-TTTCCA-CT-GCAA---A--C-TCA-AA [ 640]
baaaf62e-d0ee-4c49-9d83-0313d3df  GATTTTCAGTTTTCAGAAGGAAT----AAAAGA-TTTT--GATAACA-CTA-TTC-CA-CT-GCAA---A--C-TCA-AA [ 640]
6771788a-69b9-4fd8-b4e0-073881fe  AAAATACATTTTTCCAAAGAAAT----AAAAGA-TTTT--GATAACG-CTA-TTTCCA-CT-GCAA---A--C-TC---A [ 640]
a6666a5a-6885-427a-9881-430dad33  AAGATACATTTTCAGAAGGAATA----AAAGAT-TTT---GATAACG-CTA-TTTCCA-CT-GCAA---A--C-TCA-AA [ 640]
5be1c0db-7e16-4441-a2ab-f8107a5d  AAGATACATTT-TCAGAAGGAAT----AAAAGA-TTTT--GATAACC-CTA-TTT-CA-CT-GCAA---A--C-TC---G [ 640]
99a0bd03-c448-4605-9def-185e9f2d  AAGATACATTTTCAGAAGGAATA----AAAGA--TTTT--GATAACG-CTA-TTTCCA-CT-GCAA---A--C-TCA-AA [ 640]
c37ace00-c1f8-46e8-bcdc-c3c33bcb  AAG-TACATTTTCAGAAGGAATA----AAAGA--TTTT--GATAACG-CTA-TTTCCA-CT-GCAA---A--C-TCA-AA [ 640]
cdc9d647-e2ac-442b-8435-6dd7b82f  AAGATACATTTTCAGAAGGAATA----AAAGAT-TTTT--GATAACG-CTA-TTTCCA-CT-GCAA---A--C-TCG-AA [ 640]
bc10b467-59cf-4610-94fe-267549f8  ATACATTTTCA--GAAGGAATAA------AAGA-TTTT--GATAACG-CTA-TTTCCA-CT-GCAA---A--C-TCA-AA [ 640]
c8c3f0cb-c211-4877-8f07-1765b113  ATACATTTTCA--GAAGGAATAA------AAGA-TTTT--GATAACG-CTA-TTTCCA-CT-GCAA---A--C-TCA-AA [ 640]
720a0247-e02e-400b-901f-673d7527  AAGATACATTT--CAGAAGAATA------AAGA-TTTT--GATAACG-CTA-TTTCCA-CT-GCAA---A--C-TCA-AA [ 640]
f3f8defb-2c03-44af-abc8-d94623f7  AAGATACATTTTC------AGAAGGAATAAAAGATTTT--GATAACG-CTA-TTTCCA-CT-GCAA---A--C-TCA-AA [ 640]
fc2a9b4c-2c5f-43ed-8ea7-f2af0743  AAGAATCATTTC-------AGAAGGAATAAAAGATTTT--GATAACG-CTA-TTTCCA-CT-GCAA---A--C-TCA-AA [ 640]
477953c0-cb9c-400c-bf05-dfadb08e  AAGATACATTTTC------AGAAGGAATAAAAGATTTT--GATAACG-CTA-TTTCCA-CT-GCAA---A--C-TCG-AA [ 640]
```

```
28c5afa3-4ebf-4ffa-a71d-2171913e  AGAATACATTTC-------AGAAGGAATAAAAGATTTT--GATAACG-CTA-TTTCCA-CT-GCAA---A--C-TAG-AA [ 640]
52a4f968-3394-459b-9ee3-10eab4a7  AAGATACATTTTC------AGAAGGAATAAAAGATTTT--GATAACA-TTA-TTGCC--CT-GCAA---A--C-TCA-AA [ 640]
d36d5471-42a7-440b-bd5e-48d77156  AAGATACATTTC-------AGAAGGAATAAAAGATTTT--GATAACG-CTA-TTTCCA-CT-GCAA---A--C-TCA-AA [ 640]
f7886442-23fe-473a-9ead-a2b8a2a7  AAGATACATTTT--AGAAGGAAT----AAAAGA-TTTT--GATAACG-CTA-TTTCCA-CT-GCAA---A--C-TCA-AA [ 640]
f4770134-2a4b-449d-a4f6-c99f3383  AAGATACATTTT-CCAAAGAAATCAGAAAAAGA-TTTT--GATAACG-CTA-TTTCCA-CT-GCAA---A--C-TCA-AA [ 640]
4dd53951-3b2d-42af-82ef-399bb712  AAGATACATTTT-CAGAAGGAAT----AAAAGA-TTTT--GATAACG-CTA-TTTCCA-TC-CAAA---------CT-CA [ 640]
81ea3bac-8208-46c7-9546-bbd55b7c  AAGATACATTTT-CAGAAGGAAT----AAAAGA-TTTT--GATAACG-CTA-TTTCCA-CT-GCAA---A--C-TCA-AA [ 640]
1e66a4e2-42fd-4fa9-b001-9bb7f263  AAGATACATTTT-CAGAAGGAAT----AAAAGA-TTTT--GATAACG-CTA-TTTCCA-CT-GCAA---A--C-TCA-AA [ 640]
b75fad9a-877e-4f52-968b-9fcbb7a4  AAGATACATTTT-CAGAAGGGAT----AAAAGA-TTTT--GATAACG-CTA-TTTCCA-CT-GCAA---A--C-TCA-AA [ 640]
f512b649-4f14-4eb6-b5a2-b0543441  AAGATACATTTT-CAGAAGGAAT----AAAAGA-TTTT--GATAACG-CTA-TTTCCA-CT-GCAA---A--C-TCA-AA [ 640]
5021e737-de02-42f2-920d-18868672  AAGATACATTTTA----GAAGGA--ATAAAAGA-TTTT--GATAACG-CTA-TTTCCA-CT-GCAA---A--C-TCA-AA [ 640]
a46adbf5-3ba1-48b8-a762-2403bda7  AGAATACAT-TTTCAAAAGAGAT--GAAAAAGA-TTTT--GATAACG-CTA-TTTCCA-CT-GCAA---A--C-TCA-AA [ 640]
088a7e31-4c6b-4ff7-a19b-ff14531f  AAGATACAT-TTTCAGAAGGAAT----AAAAGA-TTTT--GATAACG-CTA-TTTCCA-CT-GCAA---ACTG-GAA-GA [ 640]
e82639d1-c451-4263-90a8-2b1ce6a1  AAGATACATTTTCCAGAAGGAAT----AAAAGA-TTTT--GATAACA-CTA-ATTTCC-AT-AC-C---AAAC-TCA-AA [ 640]
3cca4221-4eee-44ed-9fb4-ebbac961  AAGATACATTTT-CAGAAGGAAT----AAAAGA-TTTT--GATAACG-CTA-TTTCCA-CT-GCAA---A--C-TCA-AA [ 640]
d7a80796-8741-4878-b6ca-23aa66f7  AAGATACATTTT-CAGAAGGAAT----AAAAGA-TTTT--GATAACG-CTA-TTTCCA-CT-GCAA---A--C-TCA-AA [ 640]
421e3eda-83a0-4ebd-972c-f24667ee  AAGATACATTTT-CAGAAGGAAT----AAAAGA-TTTT--GATAACG-CTA-TTTCCA-CT-GCAG---AAAC-TCA-AA [ 640]
e6a39029-4e34-4457-b2a9-0cf6f207  AAGATACATTTT-CAGAAGGAAT----AAAAGA-TTTT--GATAACG-CTA-TTTCCA-CT-GCAA-----AC-TCA-AA [ 640]
a1507163-34c9-4b24-b21e-3b5dfc54  AA---------CACTCC-AGAAGGAATAAAAGA-TTTT--GATAACG-CTA-TTTCCA-CT-GCAA---A--A-CCA-AA [ 640]
51b01061-aaf5-4585-ba72-1a83671b  AGAATACATTAGGAAGA-AAAATTGAAAAAAGA-TTTT--GATAACA-CTA-TTTCCA-CT-GCAA---A--C-TCA-AA [ 640]
85703d1f-2c69-4541-ab66-acb7b838  AG-ATACATTTTCAGAA-GGA---ATAAAAGAT-TTTT--GATAACG-CTA-TTTCCA-CT-GCAA---A--C-TCA-AA [ 640]
ed39aa3b-d227-4de1-92bd-5ace4420  AG-ATACATTTTCAGAA-GGA---ATAAAAGA--TTTT--GATAACG-CTA-TTTCCA-CT-GCAA---A--C-TCA-AA [ 640]
40e2af31-875d-446e-90a0-6e16a943  G----ATAATTTCAGAA-GGA---ATAAAAGA--TTTT--GATAACG-CTA-TTTCCA-CT-GCAA---A--C-TCA--A [ 640]
873c76b4-8a29-4846-98df-cb5abcc9  AG-ATACATTTTAGAAG-GAAT--AAAAGA----TTTT--GATAACG-CTA-TTTCCA-CT-GCAA---A--CTAGA-AA [ 640]
59ef08a5-aa1d-4d15-ba6f-1a925971  AG-ATACATTTTCAGAA-GGAAT-GAAGGA-----TTT--GATAACG-CTA-TTTCCA-CT-GCAA---A--C-TCA-A- [ 640]
e1cae4d8-4f39-4401-8d82-5c5edaf3  AT-GAAGATTTTCAGAA-GGAAT-AAAAGA----TTTT--GATAACG-CTA-TTTCCA-CT-GCAA---A--C-TCA--- [ 640]
5f47af75-253d-48b0-8b3e-ae59a63d  AG-ATACATTTTCCAAA-GGAAT-AAAAAGATT-TTTT--GATAAGGTCTA-TTTCCA-CT-GCAA---A--C-TCA-AA [ 640]
44f1962a-fce6-436a-9ea0-ef4a969d  AG-ATACATTTTCAGAA-GGAAT-CA-AAA----TTTT--GATAACGCCTA-TTTCCA-CT-GCAA---A--C-TCA-AA [ 640]
4544809e-f4f9-4ff0-8142-5a2a2bc9  AG-ATACGT-TTCAGAA-GGAAT-AAAAGA----TTTT--GATAACG-CTA-TTTCCA-CT-GCAA---A--C-TCA-AA [ 640]
9faa5557-cdc1-466c-815f-fb18a15f  GA-AGTACATTTTC----AGAAGGAATAAAAGA-TTTT--GATAACG-CTA-TTTCCA-CT-GCAA---A--C-TCA-AA [ 640]
ef1a361c-a80e-43f6-8e1f-aeafa681  AAGATACATTTTCAG-AAGGAAT----AAAAGA-TTTT--GATAACG-CTA-TTTCCA-TG-CAAA---C--T-CAA-AG [ 640]

Cp_PARENT                         G--AGAGA-ACTTCT-TCT--TTTGTTATG-TACAATTGAAAACA-AA--TGTAGGCAT--A-AG-TTAGGAAAG--CCA [ 720]
Ct_PARENT                         G--AGAGA-ACTTCT-TCT--TTTGTTATG-TACAATTGAAAACA-AA--TGTAGGCAT--A-AG-TTAGGAAAG--CCA [ 720]
bdf46f5b-2833-4b26-892f-323dd4c3  G--A---A-AACTCT-TCT--TTTGTTATG-TACAATTGAAAACA-AA--TGTAGGCAT----AG-TTCGGAAAG--CCA [ 720]
c40b234c-b999-47c3-93ca-016af61e  G--AGAGA-ACTTCT-TCT--TTTGT--TA-TGTAATTGAAAACA-AA--TGTAGGCAT--A-AG-TTAGGAAAG--CCA [ 720]
0d8edcbc-b350-4d00-afbc-fd78a1f6  G--AGAGA-ACTTCT-TCT--TTTGTTATG-TACAATTGAAAACA-AA--TGTAGGCAT--A-AG-TTAGGAAAG--CCA [ 720]
9cb07723-65ed-4c96-ac19-4234ea98  G--AGAGA-ACTTCT-TCT--TTTGTTATG-TACAATTGAAAACA-AA--TGTAGGCAT--A-AG-TTAGGAAAG--CCA [ 720]
ed18c856-4d5b-4622-9fe8-9b5740af  G--AGAGA-ACTTCT-TCT--TTTGTTATG-TA-AATTGAAAACA-AA--TGTAGGCAT----AG-TTAGGAAAG--CCA [ 720]
c344612d-f873-4743-818c-65979638  A--AGAGA-ACTTCT-TCT--TTTGTTATG-TA-AATTGAGAACA-AA--TG-TAGCAT--A-AG-TTAGGAAAG--CCA [ 720]
baaaf62e-d0ee-4c49-9d83-0313d3df  G--AGAGA-ACTTCT-TCT--TTTGTTATG-TACAATTGAAAACA-AA--TGTAGGCAT--A--G-TTAGGAAAG--CCA [ 720]
6771788a-69b9-4fd8-b4e0-073881fe  A--AGAGA-ACTTCT-TCT--TTTGTTATG-TACAATTGAAAACA-AA--TGTAGGCAT--A-AG-TTAGGAAAG--CCA [ 720]
a6666a5a-6885-427a-9881-430dad33  G--AGAGA-ACTTCT-TCT--TTTGTTATG-TACAATTGAAAACA-AA--TGTAGGCAT--A-AG-TTAGGAAAG--CCA [ 720]
5be1c0db-7e16-4441-a2ab-f8107a5d  A--AGAGA-ACTTCT-TCT--TTTGTTATG-TACAATTGAAAACA-AA--TGTAGGCAT--A-AG-TTAGGAAAG--CCA [ 720]
99a0bd03-c448-4605-9def-185e9f2d  G--AGAGA-ACTTCT-TCT--TTTGTTATG-TACAATTGAAAACA-AA--TGTAGGCAT--A-AG-TTAGGAAAG--CCA [ 720]
```

```
c37ace00-c1f8-46e8-bcdc-c3c33bcb  A--GAG---ACTTCT-TCT--TTTGTTATG-TACAATTGAAAACA-AA--TGTAGGCAT----AG-TTAGGAAAG--CCA [ 720]
cdc9d647-e2ac-442b-8435-6dd7b82f  G-----GA-ACTTCT-TCT--TTTGTTATG-TACAATTGAAAACA-AA--TGTAGGCAT--A-AG-TTAGGAAAG--CCA [ 720]
bc10b467-59cf-4610-94fe-267549f8  G--AGAGA-ACTTCT-TCT--TTTGTTATG-TACAATTGAAAACA--AATGTCAGGCAT--A-AG-TTAGGAAAG--CCA [ 720]
c8c3f0cb-c211-4877-8f07-1765b113  G--AGAGA-ACTTCT-TCT--TTTGTTATG-TAAA-TTGAAAACA--AATGT-AGGCAT--A-AG-TTAGGAAAG--CCA [ 720]
720a0247-e02e-400b-901f-673d7527  G--AGAGA-ACTTCT-TCT--TTTGTTATG-TACAATTGAAAACA--AATGT-AGGCAT--A-AG-TTAGGAAAG--CCA [ 720]
f3f8defb-2c03-44af-abc8-d94623f7  G--AGAGA-ACTTC---TTCTTTTGTTATG-TACAATTGAAAACA-AA--TGTAGGCAT--A-AG-TTAGGAAAG--CCA [ 720]
fc2a9b4c-2c5f-43ed-8ea7-f2af0743  G--AGAGA-ACTTCT-TCT--TTTGTTATG-TACAATTGAAAACA-AA--TGTAGGCAT--A-AG-TTAGGAAAG--CCA [ 720]
477953c0-cb9c-400c-bf05-dfadb08e  G--AGAGA-ACTTCT-TCT--TTTGTTATG-TACAATTGAAAACA-AA--TGTAGGCAT--A-AG-TTAGGAAAG--CCA [ 720]
28c5afa3-4ebf-4ffa-a71d-2171913e  G--AAAAC-TCTT-T-TTTCTTTTGTTATG-TACAATTAAAACAA-AT--GTGTCGCAT--A--G-CTAGGAAAG--CCA [ 720]
52a4f968-3394-459b-9ee3-10eab4a7  G--AGAGA-ACTTCT-TCT--TTTGTTATG-TACAATTGAAAACA-AA--TGTAGGCAT--A-AG-TTAGGAAAG--CCA [ 720]
d36d5471-42a7-440b-bd5e-48d77156  G--AGAGA--CTTCT-TCT--TTTGTTATG-TACAATTGAAAACA-AA--TGT-GGCGT--A-AG-TTAGGAAAG--CCA [ 720]
f7886442-23fe-473a-9ead-a2b8a2a7  G--AGAGA-ACTTCT-TCT--TTTGTTATG-TACAATTGAAAACA-AA--TGTAGGCAT--A-AG-TTAGGAAAG--CCA [ 720]
f4770134-2a4b-449d-a4f6-c99f3383  G--AGAGA-ACTTCT-TCT--TTTGTTATG-TACAATTGAAAACA-AA--TGTAGGCAT--A-AG-TTAGGAAAG--CCA [ 720]
4dd53951-3b2d-42af-82ef-399bb712  A--AGAGA-ACTTCT-TCT--TTTGTTATG-TACAATTGAAAACA-AA--TGTAGGCAT--A-AG-TTAGGAAAG--CCA [ 720]
81ea3bac-8208-46c7-9546-bbd55b7c  G--AGAGA-ACTTCT-TCT--TTTGTTATG-TACAATTGAAAACA-AA--TGTAGGCAT--A-AG-TTAGGAAAG--CCA [ 720]
1e66a4e2-42fd-4fa9-b001-9bb7f263  G--AGAGA-ACTTCT-TCT--TTTGTTATG-TACAATTGAAAACA-AA--TGTAGGCAT--A-AG-TTAGGAAAG--CCA [ 720]
b75fad9a-877e-4f52-968b-9fcbb7a4  G----AGA-ACTTCT-TCT--TTTGTTATG-TACAATTGAAAACA-AA--TGTAGGCAT--A-AG-TTAGGAAAG--CCA [ 720]
f512b649-4f14-4eb6-b5a2-b0543441  G--AGAGA-ACTTCT-TCT--TTTGTTATG-TACAATTGAAAACA-AA--TGTAGGCAT--A-AG-TTAGGAAAG--CCA [ 720]
5021e737-de02-42f2-920d-18868672  G--AGAGA-ACTTCT-T--CTTTTGTTATG-TACAATTGAAAACA-AA--TGTAGGCAT--A-AG-TTAGGAAAG--CCA [ 720]
a46adbf5-3ba1-48b8-a762-2403bda7  G--AGAGA-ACTTCT-TCT--TTTGTTATG-TACAATTGAAAACA-AA--TGTAGGCAT--A-AG-TTAGGAAAG--CCA [ 720]
088a7e31-4c6b-4ff7-a19b-ff14531f  A--AGAGA-ACTTCT-TCT--TTTGTTATG-TACAATTGAAAACA-AA--TGTAGGCAT--A-AG-TTAGGAAAG--CCA [ 720]
e82639d1-c451-4263-90a8-2b1ce6a1  G--AGAGA-ACTTCT-TCT--TTTGTTATG-TACAATTGAAAACA-AA--TGTAGGCAT--A-AG-TTAGGAAAG--CCA [ 720]
3cca4221-4eee-44ed-9fb4-ebbac961  G--AGAGA-ACTTCT-TCT--TTTGTTATG-TACAATTGAAAACA-AA--TGTAGGCAT--A-AC-TCA--AAGC--CAT [ 720]
d7a80796-8741-4878-b6ca-23aa66f7  G--AGAGA-ACTTCT-TCT--TTTGTTATG-TAAATTGAAAACAA-AT--GTAGGCATT--A-AG-TTAGGAAGA--CCA [ 720]
421e3eda-83a0-4ebd-972c-f24667ee  G--AGAGA-ACTTCT-TCT--TTTGTTATG-TACAATTGAAAACA-AA--TGTAGGCAT--A-AG-TTAGGAAAG--CCA [ 720]
e6a39029-4e34-4457-b2a9-0cf6f207  G--AGAGA-ACTTCT-TCT--TTTGTTATG-TACAATTGAAAACA-AA--TGTAGGCAT--A-AG-TTAGGAAAG--CCA [ 720]
a1507163-34c9-4b24-b21e-3b5dfc54  G--A-AGA-ACTTCT-TCT--TTTGTTATG-T--AATTAGAAACA-AA--TGTA-GCAT--A-AG-TTAGGAAAG--CCA [ 720]
51b01061-aaf5-4585-ba72-1a83671b  G--AGAGA-ACTTCT-TCT--TTTGTTATG-TACAATTGAAAACA-AA--TGTAGGCAT--A-GT-TACAGAAAA--CCA [ 720]
85703d1f-2c69-4541-ab66-acb7b838  G--AGAAG-C---CT-TTC--TTTGTTATG-TACAATTGAAAACA-AA--TGTAGGCAT--A-AG-TTAGGAAAG--CCA [ 720]
ed39aa3b-d227-4de1-92bd-5ace4420  G--AGAGA-ACTTCT-TCT--TTTGTTATG-TACAATTGAAAACA-AA--TGTAGGCAT--A-AG-TTAGGAAAG--CCA [ 720]
40e2af31-875d-446e-90a0-6e16a943  ---AGAGA-ACTTCT-TCT--TTTGTTATG-TACAATTGAAAACA-AA--TGTAGGCAT--A------ACCAAAG--CCA [ 720]
873c76b4-8a29-4846-98df-cb5abcc9  G--AGAGA-ATTTCT-TCT--TTTGTTATG-TACAATTGAAAACA-AA--TGTAGGCAT--A-AG-TTCAGAAAG--TCA [ 720]
59ef08a5-aa1d-4d15-ba6f-1a925971  ---AGAGA-ACTTCT-TCT--TTTGTTATG-TACAATTGAAAACA-AA--TGTAGGCAT--A-AG-TTAGGAAAG--CCA [ 720]
e1cae4d8-4f39-4401-8d82-5c5edaf3  --AAGAGA-ATTTCT-TCT--TTTGTTATG-TACAATTGAAAACA-AA--TGTAGGCAT--A-AG-TTAGGAAAG--CCA [ 720]
5f47af75-253d-48b0-8b3e-ae59a63d  G--AGAGA-ACTTCT-TCT--TTTGTTATG-TACAATTGAAAACA-AA--TGTAGGCAT--A-AC-TTAGGAAAG--CCA [ 720]
44f1962a-fce6-436a-9ea0-ef4a969d  G--AGAGA-ACTTCT-TCT--TTTGTTATG-TACAATTGAAAACA-AA--TGTAGGCAT-AC-AT-TTAGGAAAG--CCA [ 720]
4544809e-f4f9-4ff0-8142-5a2a2bc9  G--AGAGA-ACTTCT-TCT--TTTGTTATG-TTC-ATTGAAAACA-AA--TGTGCGCAT--A-AG-TTAGGAAAG--CCA [ 720]
9faa5557-cdc1-466c-815f-fb18a15f  G--AG-AG-ACTTCT-TCT--TTTGTTATG-TACAATTCAAAACA-AA--TGTAGGCAT--A-AG-TTAGGAAAG--CCA [ 720]
ef1a361c-a80e-43f6-8e1f-aeafa681  A----AGA-ATTTCT-TCT--TTTGTTATG-TACAATTAAA-ACA-AA--TGTAGGCAT--A-AG--TTCAAAAC--TCA [ 720]

Cp_PARENT                         TT-AGT-CTCTTGTAA-TTA--TTCT-GAG---ATAATGAAAAGATG-G---------TAT-AAAAAGGATTAATGAAA-  [ 800]
Ct_PARENT                         TT-AGT-CTCTTGTAA-TTA--TTCT-GAG---ATAATGAAAAGATG-G---------TAT-GAAAAGGATTAATGAAA-  [ 800]
bdf46f5b-2833-4b26-892f-323dd4c3  TT-AGTTCTCTTGTAA-TTA--TTCT-GAG---ATAATGAAAAGAT--------GG--TAT-GAAAAGGATTAATGAAAT  [ 800]
c40b234c-b999-47c3-93ca-016af61e  TT-AGT-CTCTTGTAA-TTA--TTCT-GAG---ATAATGAAAAGAT--------GG--TAT-GAAAAGGATTAATGAAAT  [ 800]
0d8edcbc-b350-4d00-afbc-fd78a1f6  TT-AGT--TCTCTTGA-TCT--TTCT-GAG---ATAATGAAAAGAT--------GG--TTG-TAAAAGGATTAATGAAAT  [ 800]
```

```
9cb07723-65ed-4c96-ac19-4234ea98  TT-AGT-CTCTTGTAA-TTA--TTCT-GAG---ATAATGAAAAGAT--------GG--TAT-GAAAAGGATTAATGAAAT [ 800]
ed18c856-4d5b-4622-9fe8-9b5740af  TT-AGT-CTCTTGTGA-TTC--CTTC-TCA---ATAATGAAAAGAT--------GG--TAT-GAAAAGGATTAATGAGAT [ 800]
c344612d-f873-4743-818c-65979638  TT-AGT-CTCTTGTAA-TTA--TTCT-GAG---ATAATGAAAAGAT--------GG--TAT-GAAAAGGATTAATAAAAT [ 800]
baaaf62e-d0ee-4c49-9d83-0313d3df  TT-AGT-CTCTTGTAA-TTA--TTCT-GAG---ATAATGAAAAGAT--------GG--TAT-GAAAAGGATTAATGAAAT [ 800]
6771788a-69b9-4fd8-b4e0-073881fe  TT-AGT-CTATTGTAA-TTA--TTCT-AG----ATAATGAAAAGAT--------GG--TAT-GAAAAGGATTAATGAAAT [ 800]
a6666a5a-6885-427a-9881-430dad33  TT-AGT-CTCTTGTAA-TTA--TTCT-GAG---ATAATGAAAAGAT--------GG--TAT-GAAAAGGATTAATGAAAT [ 800]
5be1c0db-7e16-4441-a2ab-f8107a5d  TT-AGT-CTCTTGTAA-TTA--TTCT-GAG---ATAATGAAAAGAT--------GG--TAT-GAAAAGGATTAATGAAAT [ 800]
99a0bd03-c448-4605-9def-185e9f2d  TT-AGT-CTCTTGTAA-TTA--TTCT-GAG---ATAATGAAAAGAT--------GG--TAT-GAAAAGGATTAATGAAAT [ 800]
c37ace00-c1f8-46e8-bcdc-c3c33bcb  TT-AGT-CTCTTGTAA-TTA--TTCT-GAG---ATAATGAAAAGAT--------GG--TAT-GAAAAGGATTAATGAAAT [ 800]
cdc9d647-e2ac-442b-8435-6dd7b82f  TT-AGT-CTCTTGTAA-TTA--TTCT-GAG---ATAATGAAAAGAT--------GG--TAT-GAAAAGGATTAATGAAAT [ 800]
bc10b467-59cf-4610-94fe-267549f8  TT-AGT-CTCTTGTAA-TTA--TTCT-GAG---ATAATGAAAAGAT--------GG--TAT-GAAAAGGATTAATGAAAT [ 800]
c8c3f0cb-c211-4877-8f07-1765b113  TT-AGT-CTATTGTAA-TTA--TTCT-GAG---ATAATGAAAAGAT--------GG--TAT-GAAAAGGATTAATGAAAT [ 800]
720a0247-e02e-400b-901f-673d7527  TT-AGT-CTCTTGTAA-TTA--TTCT-GAG---ATAATGAAAAGAT--------GG--TAT-GAAAAGGATTAATGAAAT [ 800]
f3f8defb-2c03-44af-abc8-d94623f7  TT-AGT-CTCTTGTAA-TTA--TTCT-GAG---ATAATGAAAAGATG-GTATGA----------AAAGGATTAATGAAAT [ 800]
fc2a9b4c-2c5f-43ed-8ea7-f2af0743  TT-AGT-CTCTTGTA----A--TTCT-GAG---ATAATGAAAAGATG-GTATGA----------AAAGGATTAATGAAAT [ 800]
477953c0-cb9c-400c-bf05-dfadb08e  TT-AGT-CTCTTGTAA-TTA--TTCT-GAG---ATAATGAAAAGATG-GTATGA----------AAAGGATTAATGAAAT [ 800]
28c5afa3-4ebf-4ffa-a71d-2171913e  TT-AGT-CTCTTGTAA-TTA--TTCT-GAG---ATAATGAAAAGATG-GTATGA----------AAAGGATTAATGAAAT [ 800]
52a4f968-3394-459b-9ee3-10eab4a7  TT-AGT-CTCTTGTAA-TTA--TTCT-GAG---ATAATGAAAAGATG-GTATGA----------AAAGGATTAATGAAAT [ 800]
d36d5471-42a7-440b-bd5e-48d77156  TT-AGT-CTCTTGTAA-TTA--TTCT-GAG---ATAATGAAAAGATG-GTATGA----------AAAGGATTAATGAAAT [ 800]
f7886442-23fe-473a-9ead-a2b8a2a7  TT-AGT-CTCTTGTAA-TTA--TTCT-GAG---ATAATGAAAAGATG-GT---------AT-GAAAAGGATTAATGAAAT [ 800]
f4770134-2a4b-449d-a4f6-c99f3383  TT-AGT-CTCTTGTAA-TTA--TTCT-GAG---ATAATGAAAAGATG-GT---------AT-GAAAAGGATTAATGAAAT [ 800]
4dd53951-3b2d-42af-82ef-399bb712  TT-AGT-CTATTGTAA-TTA--TTCT-GAG---ATAATGAAAAGATG-GT---------AT-GAAAAGGATTAATGAAAT [ 800]
81ea3bac-8208-46c7-9546-bbd55b7c  TT-AGT-CTCTTGTAA-TTA--TTCT-GAG---ATAATGAAAAGATG-GT---------AT-GAAAAGGATTAATGAAAT [ 800]
1e66a4e2-42fd-4fa9-b001-9bb7f263  TT-AGT-CTCTTGTAA-TTA--TTCT-GAG---ATAATGGAAAGAAT-------GG--TAT-GAAAAGGATTAATGAAAT [ 800]
b75fad9a-877e-4f52-968b-9fcbb7a4  TT-AGT-CTCTTGTAA-TTA--TTCT-GAG---ATAATGAAAAGATG-------GT--A-T-GAAAAGGATTAATGAAAT [ 800]
f512b649-4f14-4eb6-b5a2-b0543441  TT-AGT-CTCTTGTAA-TTA--TTCT-GAG---ATAATGAAAAGATG-GT---------AT-GAAAAGGATTAATGAAAT [ 800]
5021e737-de02-42f2-920d-18868672  TT-AGT-CTCTTGTAA-TTA--TTCT-GAG---ATAATGAAAAGATG-GT---------AT-GAAAAGGA-TTAATGAAA [ 800]
a46adbf5-3ba1-48b8-a762-2403bda7  TT-AGT-CTCTTGTAA-TTA--TTCT-GAG---ATAATG-AAAAGAT-------GG--TAT-GAAAAGGATTAATCAAAT [ 800]
088a7e31-4c6b-4ff7-a19b-ff14531f  TT-AGT-CTCTTGTAA-TTA--TTCT-GAG---ATAATG-AAAAGAT-------GG--TAT-GAAAAGGATTAATGAAAT [ 800]
e82639d1-c451-4263-90a8-2b1ce6a1  TT-AGT-CTCTTGTAA-TTA--TTCT-GAG---ATAATG-AAAAGAT-------GG--TAT-GAAAAGGATTAATGAAAT [ 800]
3cca4221-4eee-44ed-9fb4-ebbac961  TT-AGT-CTCTTGTAA-TTA--TTCT-GAG---ATAATG-AAAAGAT-------GG--TAT-GAAAAGGATTAATGAAAT [ 800]
d7a80796-8741-4878-b6ca-23aa66f7  TT-AGT-CTCTTGTAA-TTA--TTCT-GAG---ATAATGA-AAAGAT-------GG--TAT-GAAAAGGATTAATGAAAT [ 800]
421e3eda-83a0-4ebd-972c-f24667ee  TT-AGT-CTCTTGTAA-TTA--TTCT-GAG---ATAATG-AAAAGAT-------GG--TAT-GAAAAGGATTAATGAAAT [ 800]
e6a39029-4e34-4457-b2a9-0cf6f207  TT-AGT-CTCTTGTAA-TTA--TTCT-GAG---ATAATTGAAAAGAT-------GG--TAT-GAAAAGGATTAATGAAAT [ 800]
a1507163-34c9-4b24-b21e-3b5dfc54  TC-ATT-TTGTAATTA-TTG----------TAGATAATGAAAAGATG-G---------TAT-GAAAAGGATTAATGAAAT [ 800]
51b01061-aaf5-4585-ba72-1a83671b  TT-AGT-CTCTTGTAA-TTA--TTCT-GAG---ATAATGAAAAGATG-G---------TAT-GAAAAGGATTAATGAAAT [ 800]
85703d1f-2c69-4541-ab66-acb7b838  TT-AGT-CTCTTGTAA-TTA--TTCT-GAG---ATAATGAAAAGATG-G---------TAT-GAAAAGGATTAATGAAAT [ 800]
ed39aa3b-d227-4de1-92bd-5ace4420  TT-AGT-CTCTTGTAA-TTA--TTGT-CAG---ATAATGAAAAGATG-G---------TAT-GAAAAGGATTAATGAAAT [ 800]
40e2af31-875d-446e-90a0-6e16a943  TT-AGT-CTCTTGTAA-TTA--TTCT-GAG---ATAATGAAAAGATG-G---------TAT-GAAAAGGATTAATGAAAT [ 800]
873c76b4-8a29-4846-98df-cb5abcc9  TT-AGT-CTCTTGTAA-TTA--TTCT-GAG---ATAATGAAGGATTG-G---------TAT-GAAAAGGATTAATGAAAT [ 800]
59ef08a5-aa1d-4d15-ba6f-1a925971  TT-AGT-CTCTTGTAA-TTA--TTCT-GAG---ATAATGAAAAGATG-G---------TAT-GAAAAGGATTAATGAAAT [ 800]
e1cae4d8-4f39-4401-8d82-5c5edaf3  TT-AGT-CTCTTGTAA-TTA--TTCT-GAG---ATAATGAAAAGATG-G---------TAT-GAAAAGGATTAATGAAAT [ 800]
5f47af75-253d-48b0-8b3e-ae59a63d  TT-AGT-CTCTTGTAA-TTA--TTCT-GAG---ATAATGAAAAGATG-G---------TAT-GAAAAGGATTAATGAAAT [ 800]
44f1962a-fce6-436a-9ea0-ef4a969d  TT-AGT-CTCTTGTAA-TTA--TTCT-GAG---ATAATGAAAAGATG-G---------TAT-GAAAAGGATTAATGAAAT [ 800]
4544809e-f4f9-4ff0-8142-5a2a2bc9  TT-AGT-CTCTTGTAA-TTA--TTAT-GAG---ATAATGAAAAGATG-G---------TAT-GAAAAGGATTAATGAAAT [ 800]
```

```
9faa5557-cdc1-466c-815f-fb18a15f  TT-AGT-CTCTTGTAA-TTA--TTCT-GAG---ATAATGAAAAGATG-G---------TAT-GAAAAGGATTAATGAAA- [ 800]
ef1a361c-a80e-43f6-8e1f-aeafa681  TT-AGT-CTCTTGTAA-TTA--TTCT-GAG---ATAATGAAAAGAT--------GG--TAT-GAAAAGGATTAATCAAAT [ 800]

Cp_PARENT                         --TAA-AATTCTTTG--AA--TTTAGA-TT---TACGATATGA--TAAGAA-TTTAAAAA-----AAGAAGACGA----- [ 880]
Ct_PARENT                         --TAAAAATTCTTTG--AA--TTTAGA-TT---TACGATATGA--TAAGAA-TTTAAAAA-----AAGAAGACGA----- [ 880]
bdf46f5b-2833-4b26-892f-323dd4c3  --AA-AAATTCTTTG--AA--TTTAG-ATT---TACGATATGA--TAAGAATTT-AAAAAAAGAA---G----AC-GA-- [ 880]
c40b234c-b999-47c3-93ca-016af61e  --AA-AAATTCTTTG--AA--TTTCA-AGAATGTACGATATGA--TAAGAATTT-AAAAAAAGAA---G----AC-GA-- [ 880]
0d8edcbc-b350-4d00-afbc-fd78a1f6  --AA-AAATTCTTTG--AA--TTTAG-ATT---TACGATATGA--TAAGAATTT-AAAAAAAGAA---G----AC-GA-- [ 880]
9cb07723-65ed-4c96-ac19-4234ea98  --AA-AAATTCTTTG--AA--TTTAG-ATT---TACGATATGA--TAAGAATTT-AAAAAAAGAA---G----AC-GA-- [ 880]
ed18c856-4d5b-4622-9fe8-9b5740af  AGAA-AAATTCTTTG--AA--TTTAG-ATT---TACGATATGA--TAAGAATTT-AAAAAAAGAA---G----AC-GA-- [ 880]
c344612d-f873-4743-818c-65979638  --AA-AAATTATTTG--AA--TTTAG-ATT---TACGATATGA--TAAGAATTT-TAAAAAAGAA---G----AC-GA-- [ 880]
baaaf62e-d0ee-4c49-9d83-0313d3df  --AA-AAATTCTTTG--AA--TTTAG-ATT---TACGATATGA--TAAGAATTT-AAAAAAAGAA---G----AC-GA-- [ 880]
6771788a-69b9-4fd8-b4e0-073881fe  --AA-AAATTCTTTG--AA--TTTAG-ATT---TACGATATGA--TCTAAGAAT-TTAAAAAGAA---G----AC-GA-- [ 880]
a6666a5a-6885-427a-9881-430dad33  --AA-AAATTCTTTG--AA--TTTAG-ATT--TA-CGATATGA--TAGAA--TTTAAAAAAAGAA---G----AC-GA-- [ 880]
5be1c0db-7e16-4441-a2ab-f8107a5d  --AA-AAATTCTTTG--AA--TTTAG-ATT--TT--GATATGA--TAAGAATTTAAAAAAAGAAA---G----AC-GA-- [ 880]
99a0bd03-c448-4605-9def-185e9f2d  --AA-AAATTCTTTG--AA--TTTAG-ATT---TACGATATGA--TAAGAATTT-AAAAAAAGAA---G----AC-GA-- [ 880]
c37ace00-c1f8-46e8-bcdc-c3c33bcb  --AA-AAATTCTTTG--AA--TTTAG-ATT---TACGATATGA--TAAGAATTT-AAAAAAAGAA---G----AC-GA-- [ 880]
cdc9d647-e2ac-442b-8435-6dd7b82f  --AA-AAATTCTTTG--AA--TTTAG-ATT--T--AGATATGA--TAAGAATTT-AAAAAAAGAA---G----AC-GA-- [ 880]
bc10b467-59cf-4610-94fe-267549f8  --AA-AAATTCTTTG--AA--TTTAG-ATT---TACGATATGA--TAAGAATTT-AAAAAAAGAA---G----AC-GA-- [ 880]
c8c3f0cb-c211-4877-8f07-1765b113  --AA-AAATTCTTTG--AA--TTTAG-ATT---TACGATATGA--TAAGAATTT-AAAAAAAGAA---G----AC-GA-- [ 880]
720a0247-e02e-400b-901f-673d7527  --AA-AAATTCTTTG--AA--TTTAG-ATT---TACGATATGA--TAAGAATTT-AAAAAAAGAA---G----AC-GA-- [ 880]
f3f8defb-2c03-44af-abc8-d94623f7  --AA-AAATTCTTTG--AA--TTTAG-ATT---TACGATATGA--TAAGAATTTAAAAAA-AGAA---G----AC-GA-- [ 880]
fc2a9b4c-2c5f-43ed-8ea7-f2af0743  --AA-AAATTCTTTG--AA--TTT---AGA---TTTGATATGA--TAAGAATTTAAAAAA-AGAA---G----AC-GA-- [ 880]
477953c0-cb9c-400c-bf05-dfadb08e  --AA-AAATTCTTTG--AA--TTTAG-ATT---TACGATATGA--TAAGAATTTAAAAAA-AGAA---G----AC-GA-- [ 880]
28c5afa3-4ebf-4ffa-a71d-2171913e  --AA-AAATTCTTTG--AA--TTTAG-ATT---TACGATATGA--TAAGAATTTAAAAAA-AGAA---G----AC-GA-- [ 880]
52a4f968-3394-459b-9ee3-10eab4a7  --AA-AAATTCTTTG--AA--TTTAG---------TCATATGA--TAAGAATTTAAAAAA-AGAA---G----AC-GA-- [ 880]
d36d5471-42a7-440b-bd5e-48d77156  --AA-AAATTCTTTG--AA--TTTAG-ATT---TACAATATGA--TAAGAATTTAAAAAA-AGAA---G----AC-GA-- [ 880]
f7886442-23fe-473a-9ead-a2b8a2a7  --AA-AAATTCTTTG--AA--TTTAG-ATT---TACGATATGA--TAAGAATTTAAAAAAGAGGA---G----AC-GA-- [ 880]
f4770134-2a4b-449d-a4f6-c99f3383  --AA-AAATTCTTTG--AA--TTTCCAATT---TTAGATATGA--TAAGAATTTAAAAAA-AGAA---G----AC-GA-- [ 880]
4dd53951-3b2d-42af-82ef-399bb712  --AA-AAATTCTTTG--AA--TTTAG-ATT---TACGATATGA--TAAGAATTTAAAAAA-AGAA---G----AC-GA-- [ 880]
81ea3bac-8208-46c7-9546-bbd55b7c  --AA-AAATTCTTTG--AA--TTTAG-ATT---TACGATATGA--TAAGAATTTAAAAAA-AGAA---G----ACAGA-- [ 880]
1e66a4e2-42fd-4fa9-b001-9bb7f263  --AA-AAATTCTTTG--AA--TTTAG-ATT---TACGATATGA--TAAGAATTTAAAAAA-AGAA---G----AC-GA-- [ 880]
b75fad9a-877e-4f52-968b-9fcbb7a4  --GA-ATCCCTTTAC--AA--AATAG-AAT---TTTCATGTGGTAAAGAATTCAAAAAA-AAAGACG----AT-TA-- [ 880]
f512b649-4f14-4eb6-b5a2-b0543441  --AA-AAATTCTTTG--AA--TTTAG-ATT---TACGATATGA--TAAGAATTTAAAAAA-AGAA---G----AC-GA-- [ 880]
5021e737-de02-42f2-920d-18868672  --TA-AAAATTCTTT--AA--TTTCA-GTT---TACGATATGA--TAAGAATTTAAAAAA-AGAA---G----AC-GA-- [ 880]
a46adbf5-3ba1-48b8-a762-2403bda7  --AA-AAATTCTTTG--AA--TTTAG-ATT---TACGATATGA--TAAGAATTTA--------GA---G----AC-GA-- [ 880]
088a7e31-4c6b-4ff7-a19b-ff14531f  --AA-AAATTCTTTG--AA--TTTAG-ATT---TACGATATGA--TAAGAATTTAAAAAA-AGAA---G----AC-GA-- [ 880]
e82639d1-c451-4263-90a8-2b1ce6a1  --AA-AAATTCTTTG--AA--TTTAG-ATT---TACGATATGA--TAAGAATTTAAAAAA-AGAA---G----AC-GA-- [ 880]
3cca4221-4eee-44ed-9fb4-ebbac961  --AA-AAATTCTTTG--AA--TTTCA-GAT---TTAGATATGA--TAAGAATTTAAAAAA-AGAA---G----AC-GA-- [ 880]
d7a80796-8741-4878-b6ca-23aa66f7  --AA-AAATTCTTTG--AA--TTTAG-ATT---TACGATATGA--TAAGAATTTAAAAAA-AGAA---G----AC-GA-- [ 880]
421e3eda-83a0-4ebd-972c-f24667ee  --AA-AAATTCTTTG--AA--TTTAG-ATT---TACGATATGA--TAAGAATTTAAAAAA-AGAA---G----AC-GA-- [ 880]
e6a39029-4e34-4457-b2a9-0cf6f207  --AA-AAATTCTTTG--AA--TTTAG-ATT---TACGATATGA--TAAGAATTTAAAAAA-AGAA---G----AC-GA-- [ 880]
a1507163-34c9-4b24-b21e-3b5dfc54  --AA-AAATTCTTTT--AA--AAAATA-AT---TACAATGAT----AAGAATTTGAAAAAAAGAA---G----AC-GA-- [ 880]
51b01061-aaf5-4585-ba72-1a83671b  ---A-AAATTCTTTG--AA--TTTAGA-TT---TACGATATGA--TAAGA-ATTTAAAAAAAGAA---G----AC-AA-- [ 880]
85703d1f-2c69-4541-ab66-acb7b838  --AA-AAATTCTTTG--AA--TTTAGA-TT---TACGATATA----AATA-AGTTTAAAAAAGAA---G-----C-AA-- [ 880]
```

```
ed39aa3b-d227-4de1-92bd-5ace4420  --AA-AAATTCTTTG--AA--TTTCAGATT---TACGATATGA--TAAGA-ATTTAAAAAAAGAA---G----AC-GA-- [ 880]
40e2af31-875d-446e-90a0-6e16a943  --AA-AAATTCTTTG--AA--TTTCAG--------TCATATAA--TAAGA-ATTTAAAAAAAGAA---G----AC-GA-- [ 880]
873c76b4-8a29-4846-98df-cb5abcc9  --AA-AAATTCTTTG--AA--TTTAGA-TT---TACGATATGA--TAAGA-ATTTAAAAAAAGAA---G----AC-GA-- [ 880]
59ef08a5-aa1d-4d15-ba6f-1a925971  --AA-AAATTCTTTG--AA--TTTAAA-TT---TCAGATATGA--TAAGA-ATTTAAAAAAAGAA---G----AC-GA-- [ 880]
e1cae4d8-4f39-4401-8d82-5c5edaf3  --AA-AAATTCTTTG--AA--TTTAGA--T---TACGATATCG--ATAGA-ATTT-AAAAAAGAA---G----AC-GA-- [ 880]
5f47af75-253d-48b0-8b3e-ae59a63d  --AA-AAATTCTTTTTCCAAATTAGAA-TT---TACGATATGA--TAAGA-ATTTAAAAAAAGGG---T----AA-TA-- [ 880]
44f1962a-fce6-436a-9ea0-ef4a969d  --AA-AAATTCTTTG--AA--TTTAGA-TT---TACGATATGA--TAAGA-ATTTAAAAAAAGAA---G----AC-GA-- [ 880]
4544809e-f4f9-4ff0-8142-5a2a2bc9  --AA-AAATTCTTTG--AA--TTTAGA-TT---TACGATATGA--TAAGA-ATTTAAAAAAAGAA---G----AC-GA-- [ 880]
9faa5557-cdc1-466c-815f-fb18a15f  --TAAAAATTCTTTG--AT--TAGATA-CG---ATA----TGA--TAAGAATTTAAAAAA------AGAAGACGA----- [ 880]
ef1a361c-a80e-43f6-8e1f-aeafa681  --AA-AAATTCTTTG--AA--TTTAG-ATT---TACGATATGA--TAAGAATTT-AAAAAAGAA---G----AC-GA-- [ 880]

Cp_PARENT                         TTAATATAT-TGT-----TGATAATA-CTGAAA-ATATATTAATAATA-TAAATGGTAAT-TTAATAGCTT----TACTG [ 960]
Ct_PARENT                         TTAATATAT-AGT-----TGATAATA-CTGAAA-ATATATTAATAATA-TAGATGGTAAT-TTAATAGCTT----TACTG [ 960]
bdf46f5b-2833-4b26-892f-323dd4c3  TTAATATATTAT----T-AGTTGATA-ATGAAA-ATATATTAATAATAT-AGATGGTAAT-TTAATAGCTTGCT-T---G [ 960]
c40b234c-b999-47c3-93ca-016af61e  TTAATATAT-AG----T-TGATAATA-CTGAAA-ATATATTAATAATAT-AGATGGTAAT-TTAATAGCTTTAC-T---G [ 960]
0d8edcbc-b350-4d00-afbc-fd78a1f6  TTAATATATCAG----T-TGATAATA-CTGAAA-ATATATTAATAATAT-AGATGGTAAT-TTAATAGCTTTAC-T---G [ 960]
9cb07723-65ed-4c96-ac19-4234ea98  TTAATATAT-AG----T-TGATAATA-CTGAAA-ATATATTAATAATAT-AGATGGTAAT-TTAATAGCTTTAC-T---G [ 960]
ed18c856-4d5b-4622-9fe8-9b5740af  TTAATATAT-AG----T-TGATAATA-CTGAAA-ATATATTAATAATAT-AGATGGTAAT-TTAATAGCTTTAC-T---G [ 960]
c344612d-f873-4743-818c-65979638  TTAATATAT-AG----T-TGATAATA-CTGAAA-ATATATTAATAATAT-AGATGGTAAT-TTAATAGCTTTAC-T---G [ 960]
baaaf62e-d0ee-4c49-9d83-0313d3df  TTAATATAT-AG----T-TGATAATA-CTGAAA-ATATATTAATAATAT-AGATGGTAAT-TTAATAGCTTTAC-T---G [ 960]
6771788a-69b9-4fd8-b4e0-073881fe  TTAATATAT-AG----T-TGATAATA-CTGAAA-ATATATTAATAATAT-AGATGGTAAT-TTAATAGCTTTAT-T---G [ 960]
a6666a5a-6885-427a-9881-430dad33  TTAATATAT-AG----T-TGATAATA-CTGAAA-ATATATTAATAATAT-AGATGGTAAT-TTAATAGCTTTAC-T---G [ 960]
5be1c0db-7e16-4441-a2ab-f8107a5d  TTAATATAT-AG----T-TGATAATA-CTGAAA-ATATATTAATAATAT-AGATGGTAAT-TTAATAGCTTTAC-T---A [ 960]
99a0bd03-c448-4605-9def-185e9f2d  TTAATATAT-AG----T-TGATAATA-CTGAAA-ATATATTAATAATAT-AGATGGTAAT-TTAATAGCTTTAC-T---G [ 960]
c37ace00-c1f8-46e8-bcdc-c3c33bcb  TTAATATAT-AG----T-TGATAATA-CTGAAG-ATCTATTAATAATA----ATGGTAAT-TTAATAGCTTTAC-T---G [ 960]
cdc9d647-e2ac-442b-8435-6dd7b82f  TTAATATAT-AG----T-TGATAATA-CTGAAA-ATATATTAATAATAT-AGATGGTAAT-TTAATAGCTTTAC-T---G [ 960]
bc10b467-59cf-4610-94fe-267549f8  TTAATATAT-AG----T-CGTAAATA-CTGAAA-ATATATTAATAATAT-AGATGGTAAT-TTAATAGCTTTAC-T---G [ 960]
c8c3f0cb-c211-4877-8f07-1765b113  TTAATATAT-AG----T-TGATAATA-CTGAAA-ATATATTAATAATAT-AGATGGTAAT-TTAATAGCTTTAC-T---G [ 960]
720a0247-e02e-400b-901f-673d7527  TTAATATAT-AG----T-TGATAATA-CTGAAA-ATATATTAATAATAT-AGATGGTAAT-TTAATAGCTTTAC-T---G [ 960]
f3f8defb-2c03-44af-abc8-d94623f7  TTAATATAT-AG----T-TGATAATA-CTGAAA-ATATATTAATAATAT-AGATGGTAAT-TAATAG------C-TACTG [ 960]
fc2a9b4c-2c5f-43ed-8ea7-f2af0743  TTAATATAT-AG----T-TGATAATG-CTGAAA-ATATATTAATAATAT-AGATGGTAAT-TTAATAGCTTTAT-T---G [ 960]
477953c0-cb9c-400c-bf05-dfadb08e  TTAATATAT-AG----T-TGATAATA-CTGAAA-ATATATTAATAATCT-AGATGGTAAT-TTAATAGCTTT-C-T---G [ 960]
28c5afa3-4ebf-4ffa-a71d-2171913e  TTAATATAT-AG----T-TGATAATA-CTGAAA-ATATATTAATAATAT-AGATGGTAAT-TTAATAGCTTTAC-T---G [ 960]
52a4f968-3394-459b-9ee3-10eab4a7  TTAATATAT-CC----TGTCATAATA-CTGAAA-ATATATTAATAATAT-AGATGGTAAT-TTAATAGCTTTAC-T---G [ 960]
d36d5471-42a7-440b-bd5e-48d77156  TTAATATAT-AG----T-TGATAATA-CTGAAA-ATATATTAATAATAT-AGATGGTAAT-TTAATAGCTTTAC-T---- [ 960]
f7886442-23fe-473a-9ead-a2b8a2a7  TTAATATAT-AG----T-TGATAATA-CTGAAA-ATATATTAATAATAT-AGATGGTAAT-TTAATAGCTTTAC-T---G [ 960]
f4770134-2a4b-449d-a4f6-c99f3383  TTAATATAT-AG----T-TGATAATA-CTCGAC---ATATTAATAATAT-AGATGGTAAT-TTAATAGCTTTGC-T--CG [ 960]
4dd53951-3b2d-42af-82ef-399bb712  TTAATATAT-AG----T-TGATAATA-CTGAAA-ATATATTAATAATAT-AGATGGTAAT-TTAATAGCTTTAC-T---G [ 960]
81ea3bac-8208-46c7-9546-bbd55b7c  TTAATATAT-TGATTTT-TGATAATA-CTGAAA-ATATATTTAATAATATAGATGGTAAT-TTAATAGCTTTAC-T---G [ 960]
1e66a4e2-42fd-4fa9-b001-9bb7f263  TTAATATAT-AG----T-TGATAATA-CTGAAA-ATATATTAATAATAT-AGATGGTAAT-TTAATAGCTTTAC-T---G [ 960]
b75fad9a-877e-4f52-968b-9fcbb7a4  ATATATAAG-TC----T-TGATAATA-CTGAAA-ATATATTAATAATAT-AGATGGTAAT-TTAATAGCTTTAC-T---G [ 960]
f512b649-4f14-4eb6-b5a2-b0543441  TTAATATAT-AG----T-TGATAATA-CTGAAA-ATATATTAATAATAT-AGATGGTAAT-TTAATAGCTTTAT-T---G [ 960]
5021e737-de02-42f2-920d-18868672  TTAATATAT-AG----T-TGATAATA-CTGAAA-ATATATTAATAATAT-AGATGGTAAT-TTAATAGCTTTAC-T---G [ 960]
a46adbf5-3ba1-48b8-a762-2403bda7  TTAATATAT-AG----T-TGATAATA-CTGAAA-ATATATTAATAATAT-AGATGGTAAT-TTAATAGCTTTAC-T---G [ 960]
088a7e31-4c6b-4ff7-a19b-ff14531f  TTAATATAT-AG----T-TGATAATA-CTGAAA-ATATATTAATAATAT-AGATGGTAAT-TTAATAGCTTTAC-T---G [ 960]
```

```
e82639d1-c451-4263-90a8-2b1ce6a1  TTAATATAT-AG----T-TGATAATA-CTGAAA-ATATATTAATAATAT-AGATGGTAAT-TTAATAGCTTTAC-T---G [ 960]
3cca4221-4eee-44ed-9fb4-ebbac961  TTAATATAT-AG----T-TGATAATA-CTTAAA-ATATATTAATAATAT-AGATGGTAAT-TTAATAGCTTTAC-T---G [ 960]
d7a80796-8741-4878-b6ca-23aa66f7  TTAATATAT-AG----T-TGATAATA-CTGAAA-ATATATTAATAATATTAGATGGTAAT-TTAATAGCTTTAT-T---G [ 960]
421e3eda-83a0-4ebd-972c-f24667ee  TTAATATAT-AG----T-TGATAATA-CTGAAA-ATATATTAATAATAT-AGATGGTAAT-TTAATAGCTTTAC-T---G [ 960]
e6a39029-4e34-4457-b2a9-0cf6f207  TTAATATAT-AG----T-TGATAATA-CTGAAA-ATATATTAATAATAT-AGATGGTAAT-TTAATAGCTTTCT-C---A [ 960]
a1507163-34c9-4b24-b21e-3b5dfc54  TTAATATAGTCAATA-------------TGAAA-ATATATTAGATAATATAGATGGTAATTAA--TAGCTTTAT-C---G [ 960]
51b01061-aaf5-4585-ba72-1a83671b  TTAAGAGAATATATAGT-TGATAATA-CTGAAA-ATATATTAATAATA--TAATGGTAATTTA-ATAGCTTTAT-T---G [ 960]
85703d1f-2c69-4541-ab66-acb7b838  TT-----AATGTATAGT-TGATAATA-CTGAAA-ATATATTAATAATA-TAGATGGTAATTTA-ATAGCTTTAC-T---G [ 960]
ed39aa3b-d227-4de1-92bd-5ace4420  TT-----AATATATAGT-TGATAATA-CTGAAA-ATATATTAATAATA-TAGATGGTAATTTA-ATAGCTTTAC-T---G [ 960]
40e2af31-875d-446e-90a0-6e16a943  TT-----AATATATAGT-TGATAATA-CTGAAA-ATATATTAATAATC-TAGATGGTAATTTA-ATAGCTTTAC-T---G [ 960]
873c76b4-8a29-4846-98df-cb5abcc9  TTAATATAT-----AGT-TGATAATA-CTGAAA-ATATATTAATAATA-TAGATGGTAAT-TTAATAGCTTTAC-T---G [ 960]
59ef08a5-aa1d-4d15-ba6f-1a925971  TTAATATAT-----AGT-TGATAATA-CTGAAA-ATATATTAATAATA-TAGATGGTAAT-TTAATAGCTTTAC-T---G [ 960]
e1cae4d8-4f39-4401-8d82-5c5edaf3  TTAATATAT-----AGT-TGATAATA-CTGAAA-ATATATTAATAATA-TAGATGGTAAT-TTAATAGCTTTACTT---G [ 960]
5f47af75-253d-48b0-8b3e-ae59a63d  TATAGTTG------------ATAATA-CTGAAA-ATATATTAATAATA-TAGATGGTGAT-TTAATAGCTTTAC-T---G [ 960]
44f1962a-fce6-436a-9ea0-ef4a969d  TTAATATA-----TAGT-TGATAATA-CTGAAA-ATATATTAATAATA-TAGATGGTAAT-TTAATAGCTTTAC-T---A [ 960]
4544809e-f4f9-4ff0-8142-5a2a2bc9  TTAATATATGAGCTTGA-TGATGCT---TGAAA-ATATATTAATAATA-TAGATGGTAAT-TTAATAGCTTTCT-G---A [ 960]
9faa5557-cdc1-466c-815f-fb18a15f  TTAATATATAGT------TGATAATA-CTGAAA-ATATATTAATAATA-TAGATGGTAAT-TTAATAGCTT----TGAAT [ 960]
ef1a361c-a80e-43f6-8e1f-aeafa681  TTAATATAT-AG----T-TGATAATA-CTGAAA-ATATATTAATAATAT-AGATGGTAAT-TTAATAGCTTTAC-T---G [ 960]

Cp_PARENT                         AATTTC-TCCATTA---GAT---TTAGAATAATT-TATTAG-ATTTTCATCCAATGATCCGA--TTTGAG-ATACCT-TT [1040]
Ct_PARENT                         AATTTC-TCCATTA---GAT---TTAGAATAATT-TATTAG-ATTTTCATCCAATGATCCGA--TTTGAG-ATACCT-TT [1040]
bdf46f5b-2833-4b26-892f-323dd4c3  AATTTCTTCCATTA---GATT-T--AGAATAATT-TATTAG-ATTTTCATCCAATGATCCAA--TTTGAG-ATACCT-TT [1040]
c40b234c-b999-47c3-93ca-016af61e  AATTTC-TCCATTA---GATT-T--AGAATAATT-TATTAG-ATTTTCATCCAATAATCCGA--TTTGAG-ATACCT-TT [1040]
0d8edcbc-b350-4d00-afbc-fd78a1f6  AATTTC-TCCATTA---GATT-T--AAAATAATT-TATTAG-ATTTTCATCCAATGATCCGA--TTTGAG-ATACCT-TC [1040]
9cb07723-65ed-4c96-ac19-4234ea98  AATTTC-TCCATTA---GATT-T--AGAATAATT-TATTAG-ATTTTCATCCAATGATCCGA--TTTGAG-ATACCT-TT [1040]
ed18c856-4d5b-4622-9fe8-9b5740af  AATTTC-TCCATTA---GATT-T--AGAATAATT-TATTAG-ATTTTCATCCAATGATCCGA--TTTGAG-ATACCT-TT [1040]
c344612d-f873-4743-818c-65979638  AATTTC-TCCATTA---GATT-T--AGAATAATT-TCTTAG-AT-TTCATCCAATGATCCGA--TTTGAG-ATACCT-TT [1040]
baaaf62e-d0ee-4c49-9d83-0313d3df  AATTTC-TCCATTA---GATT-T--AGAATAATT-TATTAG-ATTTTCATCCAATGATCCGA--TTTGAG-ATACCT-TT [1040]
6771788a-69b9-4fd8-b4e0-073881fe  AATTTC-TCCATTCAT------T--AGAATAATT-TATTAG-ATTTTCATCCAATGATCCGA--TTTGAG-ATACCT-TT [1040]
a6666a5a-6885-427a-9881-430dad33  AATTTC-TCCATTA---GATT-T--AGAATAATT-TATTAG-ATTTTCATCCAATGATCCGA--TTTGAG-ATA-CCTTT [1040]
5be1c0db-7e16-4441-a2ab-f8107a5d  AAATTC-TCCATTA---GATT-T--AGAATAATT-TATTAG-ATTTTCATCCAATAATGAGA--TTTGAG-ATACC--TT [1040]
99a0bd03-c448-4605-9def-185e9f2d  AATTTC-TC-ATTA---GATT-T--AGAATAATT-TATTAG-ATTTTCATCCAATGATCCGA--TTTGAG-ATACCT-TT [1040]
c37ace00-c1f8-46e8-bcdc-c3c33bcb  AATTTC-TCCATTA---GATT-T--AGAATAATT-TATTAG-ATTTTCATTGATAGATCAA----TTTGA-GATACCTTT [1040]
cdc9d647-e2ac-442b-8435-6dd7b82f  AATTTC-TCCATTA---GATT-T--AGAATAATT-TATTAG-ATTTTCATCCAATAATAAAC--AT------AATACCTT [1040]
bc10b467-59cf-4610-94fe-267549f8  AATTTC-TCCATTA---GATT-T--AGAATAATT-TATTAG-ATTTTAATAAA-TAATCCGA--TTTGAG-ATA-CCTTT [1040]
c8c3f0cb-c211-4877-8f07-1765b113  AATTTC-TCCATTA---GATT-T--AGAATAATT-TATTAG-ATTTTCATCCAATGATCCGA--TTTGAG-ATACCTTTT [1040]
720a0247-e02e-400b-901f-673d7527  AATTTC-TCCATTA---GATT-T--AGAATAATT-TATTAG-AT-TTCATCCAATGATCCGA--TTTGAG-ATACCT-TT [1040]
f3f8defb-2c03-44af-abc8-d94623f7  AATTTCTC----CATTAGATT-T--AGAATAATT-TATTAGA-TTTTCATCCAATGATCCGA--TTTGAG-ATACCT-TT [1040]
fc2a9b4c-2c5f-43ed-8ea7-f2af0743  AATTTCTC----CATTAGATT-T--AGAATAATT-TATTAGA-TTTTCATCCAATGATCCC-GATTTGAG-ATACCT-TT [1040]
477953c0-cb9c-400c-bf05-dfadb08e  AATTTCTC----CATTAGATT-T--AGAATAATT-TATTAGA-TTTTCATCCAATAATG-----ATTGAG-ATACCT-TT [1040]
28c5afa3-4ebf-4ffa-a71d-2171913e  AATTTCTC----CATTAGATT-T--AGAATAATT-TATTAGA-TTTTCATCCAATGATACAG--ATTGAG-ATACCT-TT [1040]
52a4f968-3394-459b-9ee3-10eab4a7  AATTTCTC----CATTAGATT-T--AGAATAATT-TATTAGA-TTTTCATCCAATAATT---AATTTGAG-ATACCT-TT [1040]
d36d5471-42a7-440b-bd5e-48d77156  AAATTCTC----CATTAGATT-T--AGAATAATT-TATTAA--TTTTAGTGCG-TGATTGGA--TTTGAG-ATACC--TT [1040]
f7886442-23fe-473a-9ead-a2b8a2a7  AATTTCTC----CATTAGATT-T--AGAATAATT-TATTAGA-TTTTCATCCAATGATCCCA-TTTTGAG-ATACCT-TT [1040]
f4770134-2a4b-449d-a4f6-c99f3383  AATTTCTC----CATTAGATT-T--AGAATAATT-TATTAGA-TTTTCATCCAATGATCCGA--TTTGAG-ATACCT-TT [1040]
```

```
4dd53951-3b2d-42af-82ef-399bb712  AATTTCTC----CATTAGATT-T--AGAATAATT-TATTAGA-TTTTCATCCAATGATCCGA--TTTGAG-ATACCT-TT [1040]
81ea3bac-8208-46c7-9546-bbd55b7c  AATTTCTC----CATTAGATT-T--AGAATAATT-TATTAGA-TTTTCATCCAATGATCCGA--TTTGAG-ATACCT-TT [1040]
1e66a4e2-42fd-4fa9-b001-9bb7f263  AATTTCTC----CATTAGATT-T--AGAATAATT-TATTAGA-TTTTCATCCAATGATCCAT----TGAG-ATACCT-TT [1040]
b75fad9a-877e-4f52-968b-9fcbb7a4  AATTTCTC----CATTAGATT-T--AGAATAATT-TATTAGA-TTTTCATCCAATGATCCGA--TTTGAG-ATACCT-TT [1040]
f512b649-4f14-4eb6-b5a2-b0543441  AATTTTT-----CATTAGATT-TTGGGAATAATT-TATTAGA-TTTTCATCCAATGATCCGA--TTTGAG-ATACCT-TT [1040]
5021e737-de02-42f2-920d-18868672  AATTTCTC----CATTAGATT-T--AGAATAATT-TATTAA--ATTTCATCCAATCATCCGA--TTTGAG-ATACCT-TT [1040]
a46adbf5-3ba1-48b8-a762-2403bda7  AATTTCTC----CATTAGATT-T--AGAATAATT-TATTAGA-TTTTCATCCGATGATCCGA--TTTGAG-ATACCT-TT [1040]
088a7e31-4c6b-4ff7-a19b-ff14531f  AATTTCTC----CATTAGATT-T--AGAATAATT-TATTAGA-TTTTCATCCAATGATCCGA--TTTGAG-ATACCT-TT [1040]
e82639d1-c451-4263-90a8-2b1ce6a1  AATTTCTC----CATTAGATT-T--AGAATAATT-TATTAGA-TTTTCATCCAATGATCCGA--TTTGAG-ATACCT-TT [1040]
3cca4221-4eee-44ed-9fb4-ebbac961  AATTTCTC----CATTAGATT-T--AGAATAATT-TATTAGATTTTTCATCCAATGATCCGA--TTTGAG-ATACCT-TT [1040]
d7a80796-8741-4878-b6ca-23aa66f7  AATTTCTC----CATTAGATT-T--AGAATAATT-TATTAGA-TTTTCATCCAATGATCCGA--TTTGAG-ATACCT-TT [1040]
421e3eda-83a0-4ebd-972c-f24667ee  ATTCTTCC----CATTAGATT-T--AGAATAATT-TATTAGA-TTTTCATCCAATGATCCGA--TTTGAG-ATACCT-TT [1040]
e6a39029-4e34-4457-b2a9-0cf6f207  ATTCTT------CATTAGATT-T--AGAATAATT-TATTAGA-TTTTCATCCAATGATCCGA--TTTGAG-ATACCT-TT [1040]
a1507163-34c9-4b24-b21e-3b5dfc54  A---TT-ACCATTA---GAT---TTA--ATAATT-TATTAGA-TTTATGCCGTGATTTGGAT--TTTGAG-ATACCT-TT [1040]
51b01061-aaf5-4585-ba72-1a83671b  AATTTC-TCCATTA---GAT---TTAGAATAATT-TATTCCA-ATTTTCATCCAATAAAATA--GATGAG-ATACCT-TT [1040]
85703d1f-2c69-4541-ab66-acb7b838  AATTTC-TCCATTA---GAT---TTAGAATAATT-TATTAGA-TTTTCATACATAAATTTGA--TTTGAG-ATACCT-TT [1040]
ed39aa3b-d227-4de1-92bd-5ace4420  AATTTC-TCCATTA---GAT---TTAGAATAATT-TATTAGA-TTTTCATCCAATAA---TC--CGGTAG--TACCT-TT [1040]
40e2af31-875d-446e-90a0-6e16a943  AATTTC-TCCATTA---GAT---TTAGAATAATT-TATTAGA-TTTTCATCCAATAA---TA--CGGTGG-ATCACC-TT [1040]
873c76b4-8a29-4846-98df-cb5abcc9  AATTTC-TCCATTA---GAT---TTAGAATAATT-TATTAGA-TTTTCATCCAATGATCCGA--TTTGAG-ATACCT-TT [1040]
59ef08a5-aa1d-4d15-ba6f-1a925971  AATTTC-TCATTAG---ATT---TCAGAATAACT-TACTAGA-TTTTCATCCAATGATCCGA--TTTGAG-ATACCT-TT [1040]
e1cae4d8-4f39-4401-8d82-5c5edaf3  GATTTT-TCCATTA---GAT---TTAGAATAATT-TATTAGA-TTTTCATCCAATGATCCGA--TTTGAG-ATACCT-TT [1040]
5f47af75-253d-48b0-8b3e-ae59a63d  AATTTC-TCCATCA---ATTT-ATTAAAATCATT-TATTAGA-TTTTCATACGAATAATC-C--GATTTG-ATACCT-TT [1040]
44f1962a-fce6-436a-9ea0-ef4a969d  AAG-TC-TCCATTA---GAT---TTAGAATAATT-TATTAGA-TTTTCATCCAATGATCCGA--TTTGAG-ATACCT-TT [1040]
4544809e-f4f9-4ff0-8142-5a2a2bc9  ACCTTC---CATTC---CAG---TTAGAATAATT-TATTAGA-TTTTCATCCAAATAATGCA--TAAGGA-ATACCT-TT [1040]
9faa5557-cdc1-466c-815f-fb18a15f  TCTGCA-TAG---------A---TTAGAATAATT-TATTAG-ATTTCA---TCCAATGATTA--GTTGAG-ATACCT-TT [1040]
ef1a361c-a80e-43f6-8e1f-aeafa681  AATTTC-TCCATTA---GATT-T--AGAATAATT-TATTAG-ATTTTCATCCAATGATCCGA--TTTGAG-ATACCT-TT [1040]

Cp_PARENT                         C-TGT-ATTTGCTATTC-TT--AATTAATTGATCATAAT-TGTTT-GAAGAG-TTG-AAATGCT-CATT---AT-CAT--  [1120]
Ct_PARENT                         C-TGT-ATTTGCTATTC-TT--AATTAATTGATCATAAT-TGTTT-GAAGAG-TTG-AAATGCT-CATCATTAT-CAT--  [1120]
bdf46f5b-2833-4b26-892f-323dd4c3  C-TGT-ATTTGCTATTC-TT--AATTAATTGATCATAAT-TGTTT-GAAGAG-TTG-AAATGCT-TAGTCATTA-TCA--  [1120]
c40b234c-b999-47c3-93ca-016af61e  C-TGT-ATTTGCTATTC-TT--AATTAATTGATCATAAT-TGTTG-AAAGA--GTG-AAATGCT-CATCATTAT-CAT--  [1120]
0d8edcbc-b350-4d00-afbc-fd78a1f6  T-G--TATTTGCTATTC-TT--AATTAATTGATCATAAT-TGTTT-GAAGAG-TTG-AAATGCT-CATCATTAT-CAT--  [1120]
9cb07723-65ed-4c96-ac19-4234ea98  C-TGT-ATTTGCTATTC-TT--AATTAATTGATCATAAT-TGTTT-GAAGAG-TTG-AAATGCT-CATCATTAT-CAT--  [1120]
ed18c856-4d5b-4622-9fe8-9b5740af  C-TGT-ATTCCTCTATG-TT--AATTAATTGATCATAAT-TGTTTCGAAGAG-TTG-AAATGCT-CATCATTAT-CAT--  [1120]
c344612d-f873-4743-818c-65979638  C-TGT-ATTTGCTATTC-TT--AATTAATTGATCATAAT-TGTTT-GAAGAG-TTG-AAATGCT-CATCATTAT-CAT--  [1120]
baaaf62e-d0ee-4c49-9d83-0313d3df  C-TGT-ATTTGCTATTC-TT--AATTAATTGATCATAAT-TGTTT-GAAGAG-TTG-AAATGCT-CATCATTAT-CAT--  [1120]
6771788a-69b9-4fd8-b4e0-073881fe  C-TGT-ATTTGCTATTA-C---AATTAATTGATCATAAT-TGTTT-GAAGAG-TTG-AAATGCT-CATCATTAT-CAT--  [1120]
a6666a5a-6885-427a-9881-430dad33  C-TGT-ATTTGCTATTC-TT--AATTAATTG---ATAAT-TGTTT-GAAGAG-TTG-AAATGCT-CATCATTAT-CAT--  [1120]
5be1c0db-7e16-4441-a2ab-f8107a5d  C-TGT-ATTTGCTATTC-TT--AATTAATTGATCATAAT-TGTTT-GAAGAG-TTG-AAATGCT-CATCATTAT------  [1120]
99a0bd03-c448-4605-9def-185e9f2d  C-TGT-ATTTGCTATTC-TT--AATTAATTGATCATAAT-TGTTT-GAAGAG-TTG-AAATGCT-CATCATTAT-CAT--  [1120]
c37ace00-c1f8-46e8-bcdc-c3c33bcb  C-TGT-ATTTGCTATTC-TT--AATTAATTGATCATAAT-TGTTT-GAAGAG-TTG-AAATGCT-CATCATTAT-CAT--  [1120]
cdc9d647-e2ac-442b-8435-6dd7b82f  C-TGT-ATTTGCTATTC-TT--AATTAATTGATCATAAT-TGTTT-GAAGAG-TTG-AAATGCT-CATCATTAT-CAT--  [1120]
bc10b467-59cf-4610-94fe-267549f8  C-TGT-ATTTGCTATTC-TT--AATTAATTGATCATAAT-TGTTT-GAAGAG-TTG-AAATGCT-CATCATTAT-CAT--  [1120]
c8c3f0cb-c211-4877-8f07-1765b113  C-TGT-ATTTGCTATTC-TT--AATTAATTGATCATAAT-TGTTT-GAAGAG-TTG-AAATGCT-CATCATTAT-CAT--  [1120]
720a0247-e02e-400b-901f-673d7527  C-TGT-ATTTGCTATTC-TT--AATTAATTGATCATAAT-TGTTT-GAAGAG-TTG-AAATGCT-CATCATTAT-CAT--  [1120]
```

```
f3f8defb-2c03-44af-abc8-d94623f7  C-TGT-ATTTGCTATTC-TT--AATTAATTGATCATAAT-TGTTT-GAAGAG-TTG-AAATGCT-CATCATTAT-CAT--  [1120]
fc2a9b4c-2c5f-43ed-8ea7-f2af0743  C-TGT-ATTTGCTATTC-TTAAAATTAATTGATCATAAT-TGTTT-GAAGAG-TTG-AAATGCT-CATCATTAT-CAT--  [1120]
477953c0-cb9c-400c-bf05-dfadb08e  C-TGT-ATTTGCTATTC-TT--AATTAATTGATCATAAT-TGTT--GAAGAG-TTG-AAATGCT-CATCATTAT-CAT--  [1120]
28c5afa3-4ebf-4ffa-a71d-2171913e  C-TGT-ATTTGCTATTC-TT--AATTAATTGATCATAAT-TGTTT-GAAGAG--TG-AAATGCT-CATCATTAT-CAT--  [1120]
52a4f968-3394-459b-9ee3-10eab4a7  C-TG-----TGCTATCT-TC--TAGTGATTGATCATAATCTTTTC-GAAGAG-TTG-AAATGCT-CATCATTAT-CAT--  [1120]
d36d5471-42a7-440b-bd5e-48d77156  C-TGT-ATTTGCTATTC-TT--AATTAATTGATCATAAT-TGTTT-GAAGAG-TTG-AAATGCT-CATCATTATGCAT--  [1120]
f7886442-23fe-473a-9ead-a2b8a2a7  C-TGT-ATTTGCTATTC-TT--AATTAATTGATCATAAT-TGTTT-GAAGAG-TTG-AAATGCT-CATCATTAT-CAT--  [1120]
f4770134-2a4b-449d-a4f6-c99f3383  C-TGT-ATTTGCTATTC-TT--AATTAATTGATCATAAT-TGTTT-GAAGAG-TTG-AAATGCT-CATCATTAT-CAT--  [1120]
4dd53951-3b2d-42af-82ef-399bb712  C-TGT-ATTTGCTATTC-TT--AATTAATTGATCATAAT-TGTTT-GAAGAG-TTG-AAATGCT-CATCATTAT-CAT--  [1120]
81ea3bac-8208-46c7-9546-bbd55b7c  C-TGT-ATTTGCTATTC-TT--AATTAATTGATCATAAT-TGTTT-GAAGAG-TTG-AAATGCT-CATCATTAT-CAT--  [1120]
1e66a4e2-42fd-4fa9-b001-9bb7f263  C-TGT-ATTTGCTATTC-TT--AATTAATTGATCATAAT-TGTTT-GAAGAG-TTG-AAATGCT-CATCATTAT-CAT--  [1120]
b75fad9a-877e-4f52-968b-9fcbb7a4  C-TGT-ATTTGCTATTC-TT--AATTAATTGATCATAAT-TGTTT-GAAGAG-TTG-AAATGCT-CATCATTAT-CAT--  [1120]
f512b649-4f14-4eb6-b5a2-b0543441  C-TGT-ATTTGCTATTC-TT--AATTAACTGATCATAAT-TGTTT-GAAGAG-TTG-AAATGCT-CATCATTAT-CAT--  [1120]
5021e737-de02-42f2-920d-18868672  C-TGT-ATTTGCTATTC-TT--AATTAATTGATCATAAT-TGTTT-GAAGAG-TTG-AAATGCT-CATCATTAT-CAT--  [1120]
a46adbf5-3ba1-48b8-a762-2403bda7  C-TGT-ATTTGCTATTC-TT--AATTAATTGATCATAAT-TGTTT-GAAGAG-TTG-AAATGCT-CATCATTAT-CAT--  [1120]
088a7e31-4c6b-4ff7-a19b-ff14531f  C-TGT-ATTTGCTATTC-TT--AATTAATTGATCATAAT-TGTTT-GAAGAG-TTG-AAATGCT-CATCATTAT-CAT--  [1120]
e82639d1-c451-4263-90a8-2b1ce6a1  C-TGT-ATTTGCTATTC-TT--AATTAATTGATCATAAT-TGTTT-GAAGAG-TTG-AAATGCT-CATCATTAT-CAT--  [1120]
3cca4221-4eee-44ed-9fb4-ebbac961  C-TGT-ATTTGCTATTC-TT--AATTAATTGATCATAAT-TGTTT-GAAGAG-TTG-AAATGCT-CATCATTAT-CAT--  [1120]
d7a80796-8741-4878-b6ca-23aa66f7  C-TGT-ATTTGCTATTC-TT--AATTAATTGATCATAAT-TGTTT-GAAGAG-TTG-AAATGCT-CATCATTAT-CAT--  [1120]
421e3eda-83a0-4ebd-972c-f24667ee  C-TGT-ATTTGCTATTC-TT--AATTAATTGATCATAAT-TGTTT-GAAGAG-TTG-AAATGCT-CATCATTAT-CAT--  [1120]
e6a39029-4e34-4457-b2a9-0cf6f207  C-TGT-ATTTGCTATTC-TT--AATTAATTGATCATAAT-TGTTT-GAAGAG-TTG-AAATGCT-CATCATTAT-CAT--  [1120]
a1507163-34c9-4b24-b21e-3b5dfc54  CTTGT-ATTTGCTATTC-TT--AATTAATTGATCATAATTGTTTC-GAAGAG-TTG-GAATCCT-CATGGTTTT CAT--  [1120]
51b01061-aaf5-4585-ba72-1a83671b  C-TGT-ATTTGCTATTC-TT--AATTAATTGATAATAA---TCTT-GAAGAG-TTG-AAATGCT-CATCATTAT-CAT--  [1120]
85703d1f-2c69-4541-ab66-acb7b838  C-TGT-ATTTGCTATTC-TT--AATTAATTGATCATAATTG-TTT-C----G-TTG-AAATGCT-CATCATTAT-CGT--  [1120]
ed39aa3b-d227-4de1-92bd-5ace4420  C-TGT-ATTTGCTATTC-TT--AATTAATTGATCATAATTG-TTT-GAAGAG-TTG-AAATGCT-CATCATTAT-CAT--  [1120]
40e2af31-875d-446e-90a0-6e16a943  C-TGT-ATTTGCTATTC-TT--AATTAATTGATCATAATTG--TT-GAAGAG-TTG-AAATGCT-AATCATTAT-CAT--  [1120]
873c76b4-8a29-4846-98df-cb5abcc9  C-TGT-ATTTGCTATTC-TT--AATTAATTGATCAT-AATTGTTT-GAAGAG-TTG-AAATGCT-CATCATTAT-CAT--  [1120]
59ef08a5-aa1d-4d15-ba6f-1a925971  C-TGT-ATTTGCTATTC-TT--AATTAATTGATCAT-AATTGTTT-GAAGAG-TTC-AAATGCT-CATCATTAT-CAT--  [1120]
e1cae4d8-4f39-4401-8d82-5c5edaf3  C-TGT-ATTTGCTATTC-TT--AATTAATTGATCAT-AATTGTTT-GAAGA-GTTG-AAATGCT-CATCATTAT-CAT--  [1120]
5f47af75-253d-48b0-8b3e-ae59a63d  C-TGT-ATTTGCTATTC-TT--AATTAATTGATCAT-AATTGTTT-GAAGAG-TTG-AAATGCT-CATCATTAT-AGT--  [1120]
44f1962a-fce6-436a-9ea0-ef4a969d  C-TGT-ATTTGCTATTC-TT--AATTAATTGATCAT-AATTGTTT-GAAGAG-TTG-AAATGCT-CATCATTAT-CAT--  [1120]
4544809e-f4f9-4ff0-8142-5a2a2bc9  C-TGTAATTTGCTATTC-TT--AATTAATTGATCAT-AATTGTTT-GAAGAGTTGA-AAATGCT-CATCATTAT-CAT--  [1120]
9faa5557-cdc1-466c-815f-fb18a15f  C-TGT-ATTTGCTATTC-TT--AATTAATTGATCATAAT-TGTTT-GAAGAG-TTG-AAATGCTTCATCATTAT-CAT--  [1120]
ef1a361c-a80e-43f6-8e1f-aeafa681  C-TGTTATTTGCTATTC-TT--AATTAATTGATCATAAT-TGTTT-GAAGAG-TTG-AAATGCT-CATCATTAT-CAT--  [1120]

Cp_PARENT                         AGAAATTATCATTAATTT-CAGGTTTTTTACCACTATCATACTCT-AAAGAAGATGAGCCTGATAT--TGT-AGAA-T--  [1200]
Ct_PARENT                         AGAAATGATCATTCATTT-CAGGTTTTTTACCACTATCATATTCT-AAAGAAGATGACCCTGATAT--TGT-AGAA-T--  [1200]
bdf46f5b-2833-4b26-892f-323dd4c3  TGAAATGGTGACCCATTTTCAGGTTT-TTACCACTATCATATTCTA-AAGAAGATGACCCTGATAT--TGT-AGA-AT--  [1200]
c40b234c-b999-47c3-93ca-016af61e  CAAAATGATCATTCATTTCAGGTTTTCAC-----T----ATCATCT-AAAGAAGATGACCTGATAT--TGT-AGA-AT--  [1200]
0d8edcbc-b350-4d00-afbc-fd78a1f6  AGAAATGATCATTCATTTC-AGGTTTTTTACCACTATCATATTCTA-AAGAAGATGACCCTGATAT--TGT-AGA-AT--  [1200]
9cb07723-65ed-4c96-ac19-4234ea98  AGAAATGATCATTCATTTTGAGGTTTTTTACCACTATCATATTCTA-AAGAAGATGACCCTGATAT--TGT-AGA-AT--  [1200]
ed18c856-4d5b-4622-9fe8-9b5740af  AGAAATGATCATTCATTTTCAGGTTTTTA-CCACTATCATATTCTAGAGGAACATGACCCTGATAT--TGT-AGA-AT--  [1200]
c344612d-f873-4743-818c-65979638  AGAAATGATCATTCATTTCAGGTTTTT--ACCACTATCATATTGAAGAAATAAT--------------TAT-GAT-ATAG  [1200]
baaaf62e-d0ee-4c49-9d83-0313d3df  AGAAATGATCATTCATTTCAGGTTTTT--ACCACTATCATATTCTA-AAGAAGATGACCCTGATATCTGAG-AAA-AT--  [1200]
6771788a-69b9-4fd8-b4e0-073881fe  AGAAATGATCATTCATTTCAGGTTTTTTCTCCACTATCATATTCTA-AAGAAGATGACCCTGATAT--TGT-AGA-AT--  [1200]
```

```
a6666a5a-6885-427a-9881-430dad33   AGAAATGATCATTAGTTTCAGG-TTTTTA-CCACTATCATATTCTA-AAGAAGATGACCCTGATAT--TGT-AGA-AT-- [1200]
5be1c0db-7e16-4441-a2ab-f8107a5d   AGAAATGATCATTCATTTTGG--TTTTTA-CCACTATCATATTCTA-AAGAAGATGACCCTGATAT--TGT-AGA-AT-- [1200]
99a0bd03-c448-4605-9def-185e9f2d   AGAAATGATCATTCATTTCAGGTTTTTTA-CCACTATCATATTCTA-AAGAAGATGACCCTGATAT--TGT-AGA-AT-- [1200]
c37ace00-c1f8-46e8-bcdc-c3c33bcb   AGAAATGATCATTCATTTCAGGTTTTTTA-CCACTATCA---TCTA-AAGAAGATGACCTAAATCTTCTGT-AGA-AT-- [1200]
cdc9d647-e2ac-442b-8435-6dd7b82f   AGAAATGATCATTCATTTCAGGTTTTTTA-CCACTATCATATTCTA-AAGAAGATGACCCTGATAT--TGT-AGA-AT-- [1200]
bc10b467-59cf-4610-94fe-267549f8   AGAAATGATCATTCATTTCAGGTTT-TTA-CCACTATCATATTCTA-AAGAAGATGACCCTGATAT--TGT-AGA-AT-- [1200]
c8c3f0cb-c211-4877-8f07-1765b113   AGAAATGATCATTCATTTCAGGTTTTTTA-CCACTATCATATTCTA-AAGAAGATGACCCTGATAT--TGT-AGA-AT-- [1200]
720a0247-e02e-400b-901f-673d7527   AGAAATGATCATTCATTTCAGGTTTTTTA-CCACTATCATATTCTA-AAGAAGATGACCCTGATAT--CTA-GAG-AATC [1200]
f3f8defb-2c03-44af-abc8-d94623f7   AGAAATGATCATTCATTTCAG--GTTTTTACCACTATCATATTCT-AAAGAAGATGACCCTGATAT--TGT-AGA-AT-- [1200]
fc2a9b4c-2c5f-43ed-8ea7-f2af0743   AGAAATGATCATTAGTTTTCAGGTTT----CCACTATCATATTCT-AAAGAAGATGACCCTGATAT--TGT-AGA-AT-- [1200]
477953c0-cb9c-400c-bf05-dfadb08e   AGAAATGATCATTCATTT-CAGGTTTTTTACCACTATCATATTCT-AAAGAAGATGACCCTGATAT--TGT-AGA-AT-- [1200]
28c5afa3-4ebf-4ffa-a71d-2171913e   AGAAATGATCATTCATTT-CAGGT-TTTTACCACTATCATATTCT-AAAGAAGATGACCCTAATAT--GTT-AGA-AT-- [1200]
52a4f968-3394-459b-9ee3-10eab4a7   AGAAATGATCATTCATTT-CAGGTTTTTTACCACTATCATATTCT-AAAGAAGATGACCCTGATAT--TGT-AGA-AT-- [1200]
d36d5471-42a7-440b-bd5e-48d77156   AGAAATGATCATTCATTT-CAGGTTTTTTACCACTATCATG-TCT-AAAGAAGATGACCCTGATAT--TGT-AGA-AT-- [1200]
f7886442-23fe-473a-9ead-a2b8a2a7   AGAAATGATCATTAGTGA-----------CTTACTATCATATTCTAAG---AAATCATTCTGATAT--TGT-AGA-AT-- [1200]
f4770134-2a4b-449d-a4f6-c99f3383   AGAAATGATCATTCATTT-CAGGTTTT-TACCACTATCATATTCT-AAAGAAGATGACCCTGATAT--TGT-AGA-AT-- [1200]
4dd53951-3b2d-42af-82ef-399bb712   AGAAATGATCATTCATTT-CAGGTTTTTTACCACTATCATATTAT----CAAGATGACCCTGATAT--TGT-AGA-AT-- [1200]
81ea3bac-8208-46c7-9546-bbd55b7c   AGAAATGATCATTCATTT-CAGGTTTTTTACCACTATCATATTCT-AAAGAAGATGACCCTGATAT--TGT-AGA-AT-- [1200]
1e66a4e2-42fd-4fa9-b001-9bb7f263   AGAAATGATCATTCATTT-CAGGTTTTTTACCACTATCATATTCT-AAAGAAGATGACCCTGATAT--TGT-AGA-AT-- [1200]
b75fad9a-877e-4f52-968b-9fcbb7a4   AGAAATGATCATTCATTT-CAGGTT-TTTACCACTATCATATTCT-AAAGAAGATGACCCTGATAT--TGT-AGA-AT-- [1200]
f512b649-4f14-4eb6-b5a2-b0543441   AGAAATGATCATTCATTT-CAGGTTT----CCACTATCATATTCT-AAAGAAGATGACCCTGATAT--TGT-AGA-AT-- [1200]
5021e737-de02-42f2-920d-18868672   AGAAATGATCATTCATTTTCAGGTTTTTTACCACTATCATATTCT-AAAGAAGATGACCCTGATAT--TGT-AGA-AT-- [1200]
a46adbf5-3ba1-48b8-a762-2403bda7   AGAAATGATCATTCATTT-CAGGTTTTTTACCACTATCATATTCT-AAAGAAGATGACCCTGATAT--TGT-AGA-AT-- [1200]
088a7e31-4c6b-4ff7-a19b-ff14531f   AGAAATGATCATTCATTTCAG--GTTTTTACCACTATCATATTCT-AAAGAAGATGACCCTGATAT--TGT-ACA-TA-- [1200]
e82639d1-c451-4263-90a8-2b1ce6a1   AGAAATGATCATTCATTT-CAGGTTTTTTACCACTATCATATTCT-AAAGAAGATGACCCTGATAT--TGT-AGA-AT-- [1200]
3cca4221-4eee-44ed-9fb4-ebbac961   AGAAATGATCATTCATTT-CAGGTTTTTTACCACTATCATATTCT-AA--GAAATAATTATGATAT--TGT-AGA-AT-- [1200]
d7a80796-8741-4878-b6ca-23aa66f7   AGAAATGATCATTCATTTC-AGGTTTTTTACCACTATCATATTCT-AAAGAAGATGACCCTGATAT--TGT-AGA-AT-- [1200]
421e3eda-83a0-4ebd-972c-f24667ee   AGAAATGATCATTCATTT-CAGGTTTTTTACCACTATCATATTCT-AAAGAAGATGACCCTGATAT--TGT-AGA-AT-- [1200]
e6a39029-4e34-4457-b2a9-0cf6f207   AGAAATGATCATTCATTTTCAGGTTTTTTACCACTATCATATTCT-AAAGAAGATGACCCTGATAT--TGT-AGA-AT-- [1200]
a1507163-34c9-4b24-b21e-3b5dfc54   AGAAATGATCATTCATTTTAGGTTT-----CCACTATCATATTCTA-AAGAAGATGACCCTGATAT--TGT-AGA-AT-- [1200]
51b01061-aaf5-4585-ba72-1a83671b   AGAAATGATCATTCATTTCAGGTTTTTTA-CCACTATCATATTCTA-AAGAAGATGACCTAGA-------------AT-- [1200]
85703d1f-2c69-4541-ab66-acb7b838   --CAATGATCATTCATTTTAGGTTTTTTA-CCACTATCATATTCT--AAGAAGATGACATTGA-----TGT-AGA-AT-- [1200]
ed39aa3b-d227-4de1-92bd-5ace4420   AGAAATGATCATTCATTTTAGGTTTTTTA-CCACTATCATATTCTA-AAGAAGATGACCCTGATAT--TGT-AGA-AT-- [1200]
40e2af31-875d-446e-90a0-6e16a943   AGAAATGATCATTCATTTCAGG-TTTTTA-CCACTATCATATTCTA-AAGAAGATGACCCTGATAT--TGT-AGA-AT-- [1200]
873c76b4-8a29-4846-98df-cb5abcc9   AGAAATGATCATTCATTTCGGT--TTTTA-CCACTATCATATTCT-AAAGAAGATGACCCTGATAT--TGTTCAA-AT-- [1200]
59ef08a5-aa1d-4d15-ba6f-1a925971   AGAAATGATCATTCATTTCAGGTTTTTTA-CCACTATCATATTCT-AAAGAAGATGACCCTGATAT--TGTCAGA-AT-- [1200]
e1cae4d8-4f39-4401-8d82-5c5edaf3   AGAAATGATCATTCATTTCAGGTTTTTTA-CCACTATCATATTCT-AAAGAAGATGACCCTGATAT--TGT-AGA-AT-- [1200]
5f47af75-253d-48b0-8b3e-ae59a63d   GGATG-GATCATTCATTTTAGGATTTACC-ACTATAATATATTCT-AAAGAAGATAACTATGATAT--TGT-AGA-AT-- [1200]
44f1962a-fce6-436a-9ea0-ef4a969d   AGAAATGATCATTCATTT-CAGGTTTTTA-CCACTATCATATTCT-AAAGAAGATGACCCTGATAT--TGT-AGA-AT-- [1200]
4544809e-f4f9-4ff0-8142-5a2a2bc9   AGAAATGATCATTCATTTCAGGTTTTTTA-CCACTATCATATTCT-------AAGAAACATGATAT--TGT-AGA-AT-- [1200]
9faa5557-cdc1-466c-815f-fb18a15f   AGAAATGATCATTCATTT-CAGGTTTTTTACCACTATCATATTCT-AAAGAAGATGACCCTGATAT--TGT-AGAATC-- [1200]
ef1a361c-a80e-43f6-8e1f-aeafa681   AGAAATGATCATTCATTTT-AGGTTTTTTACCACTATCATATTCTA-AAGAAGATGACCCTGATAT--TGT-AGA-AT-- [1200]

Cp_PARENT                          -----CATGATCATAATAGTTTTCTTGTTG-AATAG-GTT-CTTGTTGTTGTAGCTGGTGTCC-AAAGGTTATATT-AAA [1280]
Ct_PARENT                          -----CATGATCATAATAGTTTTCTTGTTG-AATAG-GTT-CTTGTTGTTGTAGCTGGTGTCC-AAAGGTTATATT-AAA [1280]
```

```
bdf46f5b-2833-4b26-892f-323dd4c3  -----CATGATCATAATAGTTTTCTTGTTG-AATAG-GTTC-TTGTTGTTGTAGCTGGTGTCCA-AAGGTTATATT-AAA [1280]
c40b234c-b999-47c3-93ca-016af61e  -----CATGATCATAATAGTTTTCTTGTTG-AATAG-GTTC-TTGTTGTTGTAGCTGGTGTCCAAAGGTTATATTA-AAG [1280]
0d8edcbc-b350-4d00-afbc-fd78a1f6  -----CATGATCATAATAGTTTTCTTGTTG-AATAG-GTTC-TTGTTGTTGTAGCTGGTGTCCA-AAGGTTATATT-AAA [1280]
9cb07723-65ed-4c96-ac19-4234ea98  -----CATGATCATAATAGTTTTCTTGTTG-AATAG-GTTC-TTGTTGTTGTAGCTGGTGTCCA-AAGGTTATCTA-AA- [1280]
ed18c856-4d5b-4622-9fe8-9b5740af  -----CATGATCATAATCGTTTTCTTGTTG-AATAG-GTTC-TTGTTGTTGTAGCTGGTGTC-CAAAGGTTATATT-AAA [1280]
c344612d-f873-4743-818c-65979638  AAAATCATGATCATAATAGTTTTCTTGTTG-AATAG-GTTC-TTGTTGTTGTAG-CTAGTATCCAAAGGTTATATT-AAA [1280]
baaaf62e-d0ee-4c49-9d83-0313d3df  -----CATGATCATAATAGTTTTCTTGTTG-AATAG-GTTC-TTGTTGTTGTAG-CTGGTGTCCAAAGGTTATATT-AAA [1280]
6771788a-69b9-4fd8-b4e0-073881fe  -----CATGATCATAATAGTTTTCTTGTTG-AATAG-GTTC-TTGTTGTTGTAG-CTGGTGTCCAAAGGTTATATT-CGA [1280]
a6666a5a-6885-427a-9881-430dad33  -----CATGATCATAATAGTTTTCTTGTTG-AATAG-GTTC-TTGTTGTTGTAG-CTGGTGTCCAAAGGTTATATT-AAA [1280]
5be1c0db-7e16-4441-a2ab-f8107a5d  -----CATGATCATAATAGTTTTCTTGTTG-AATAG-GTTC-TTGTTGTTGTAG-CTGGTGTCCAAAGGTTATATT-AAA [1280]
99a0bd03-c448-4605-9def-185e9f2d  -----CATGATCATAATAGTTTTCTTGTTG-AATAG-GTTC-TTGTTGTTGTAG-CTGGTGTCCAAAGGTTATATT-AAA [1280]
c37ace00-c1f8-46e8-bcdc-c3c33bcb  -----CATGATCATAATAGTTTTCTTGTTG-AATAG-GTTC-TTGTTGTTGTAG-CTGGTGTCCAAAGGTTATATT-AAA [1280]
cdc9d647-e2ac-442b-8435-6dd7b82f  -----CATGATCATAATAGTTTTCTTGTTG-AATAG-GTTC-TTGTTGTTGTAG-CTGGTGTCCAAAGGTTATATT-AAA [1280]
bc10b467-59cf-4610-94fe-267549f8  -----CATGATCATAATAGTTTTCTTGTTG-AATAG-GTTC-TTGTTGTTGTAG-CTAGTATCCAAAGGTTATATT-AAA [1280]
c8c3f0cb-c211-4877-8f07-1765b113  -----CATGATCATAATAGTTTTCTTGTTG-AATAG-GTTC-TTGTTGTTGTAG-CTGGTGTCCAAAGGTTATATT-AAA [1280]
720a0247-e02e-400b-901f-673d7527  ----ATGATCATAATAGTTTTTTCTTGTTG-AATAG-GTTC-TTGTTGTTGTAG-CTGGTGTCCAAAGGTTATATT-AAA [1280]
f3f8defb-2c03-44af-abc8-d94623f7  -----CATGATCATAATAGTTTTCTTGTTG-AATAG-GTT-CTTGTTGTTGTAGCTGG-TGTCCAAAGGTTATATT-AAA [1280]
fc2a9b4c-2c5f-43ed-8ea7-f2af0743  -----CATGATCATGATCGTTTTCTTGTTG-AATAG-GTT-CTTGTTGTTGTAGCTAC-TATACAAAGGTTATATT-AAA [1280]
477953c0-cb9c-400c-bf05-dfadb08e  -----CATGATCATAATAGTTTTCTTGTTG-AATAG-GTT-CTTGTTGTTGTAGCTGG-TGTCCAAAGGTTATATT-AAA [1280]
28c5afa3-4ebf-4ffa-a71d-2171913e  -----CATGATCATAATAGTTTTCTTGTTG-AATAG-GTT-TCTGTTGTTGTAGCTGG-TGTCCAAAGGTTATATT-AAA [1280]
52a4f968-3394-459b-9ee3-10eab4a7  -----CATGATCATAATAGTTTTCTTGTTG-AATAG-GTT-CTTGTTGTTGTAGCTGG-TGTCCAAAGGTTATATT-AAA [1280]
d36d5471-42a7-440b-bd5e-48d77156  -----CATGATAATAATGTTGTTTGTTTGG-AATAG-GTT-CTTGTTGTTGTAGCTA---GTACAAAGGTTATATT-AAA [1280]
f7886442-23fe-473a-9ead-a2b8a2a7  -----CATAATCATAAT-GTTTTCTTGTTG-AATAG-GTT-CTTGTTGTTGTAGCTGG-TGTCCAAAGGTTATATT-AAA [1280]
f4770134-2a4b-449d-a4f6-c99f3383  -----CATGATCATAATAGTTTTCTTGTTG-AATAG-GTT-CTTGTTGTTGTAGCTGG-TGTCCAAAGGTTATATT-AAA [1280]
4dd53951-3b2d-42af-82ef-399bb712  -----CATGATCATAATAGTTTTCTTGTTG-AATAG-GTT-CTTGTTGTTGTAGCTGG-TGTCCAAAGGTTATATT-AAA [1280]
81ea3bac-8208-46c7-9546-bbd55b7c  -----CATGATCATAATAGTTTTCTTGTTG-AATAG-GTT-CTTGTTGTTGTAGCTGG-TGTCCAAAGGTTATATT-AAA [1280]
1e66a4e2-42fd-4fa9-b001-9bb7f263  -----CATGATCATAATAGTTTTCTTGTTG-AATAG-GTT-CTTGTTGTTGTAGCTGG-TGTCCAAAGGTTATATT-AAA [1280]
b75fad9a-877e-4f52-968b-9fcbb7a4  -----CATGATCATAATAGTTTTCTTGTTG-AATAG-GTT-CTTGTTGTTGTAGCTGG-TGTCCAAAGGTTATATT-AAA [1280]
f512b649-4f14-4eb6-b5a2-b0543441  -----CATGATCATAATAGTTTGCTTGTTG-AATAG-GTT-CTTGTTGTTGTAGCTGG-TGTCCAAAGGTTATATT-AAA [1280]
5021e737-de02-42f2-920d-18868672  -----CATGATCATAATAGTTTTCTTGCTG-AATAG-GTT-CTTGTTGTTGTCA-------CATAAAGGTTATATT-AAA [1280]
a46adbf5-3ba1-48b8-a762-2403bda7  -----CATGATCATAATAGTTTTCTTGTTG-AATAG-GTT-CTTGTTGTTGTAGCTGG-TGTCCAAAGGTTATATT-AAA [1280]
088a7e31-4c6b-4ff7-a19b-ff14531f  -----CGTGATCATAATAGTTTTCTTGTTG-AATAG-GTT-CTTGTTGTTGTAGCTGG-TGTCCAAAGGTTATATT-AAA [1280]
e82639d1-c451-4263-90a8-2b1ce6a1  -----CATCATCATAATAGTTTTCTTGTTG-AATAG-GTT-CTTGTTGTTGTAGCTGG-TGTCCAAAGGTTATATT-AAA [1280]
3cca4221-4eee-44ed-9fb4-ebbac961  -----CATGATCATAATAGTTTTCTTGTTG-AATAG-GTT-CTTGTTGTTGTAGCTGG-TGTCCAAAGGTTATATT-AAA [1280]
d7a80796-8741-4878-b6ca-23aa66f7  -----CATGATCATAATAGTTTTCTTGTTG-AATAG-GTT-CTTGTTGTTGTAGCTGG-TGTCCAAAGGTTATATT-AAA [1280]
421e3eda-83a0-4ebd-972c-f24667ee  -----CATGATCATAATAGTTTTCTTGTTG-AATAG-GTT-CTTGTTGTTGTAGCTGG-TGTCCAAAGGTTATATT-AAA [1280]
e6a39029-4e34-4457-b2a9-0cf6f207  -----CATGATCATAATAGTTTTCTTGTTG-AATAG-GTT-CTTGTTGTTGTAGCTGG-TGTCCAAAGGTTATATT-AAA [1280]
a1507163-34c9-4b24-b21e-3b5dfc54  -----CATGATCATAATAGTTTTCTTGTTA-AAT--GGTA-TTTGTTGTTGTAGC-TAGTATCCAAAGGTTATAT-TAAA [1280]
51b01061-aaf5-4585-ba72-1a83671b  -----CATGATCATAATAGTTTTCTTGTTG-AATAG-GTT-CTTGTTGTTCTAGC-TGGTGTCCAAAGGTTATAT-TAAA [1280]
85703d1f-2c69-4541-ab66-acb7b838  -----CATGATCATAATAGTTTTCTTGTTG-AATAG-GTT-CTTGTTGTTGTAGC-TGGTGTCCAAAGGTTATAT-TAAA [1280]
ed39aa3b-d227-4de1-92bd-5ace4420  -----CATGATCATAATAGTTTTCTTGTTG-AATAG-GTT-CTTGTTGTTGTAGC-TGGTGTCCAAAGGTTATAT-TAAA [1280]
40e2af31-875d-446e-90a0-6e16a943  -----CATGATCATAATAGTTTTCTTGTTG-AATAG-GTT-CTTGTTGTTGTGGC-TCATATCCAAAGGTTATAT-TAAA [1280]
873c76b4-8a29-4846-98df-cb5abcc9  -----CATGATCATAATAGTTTTCTTGTTG-AATAG-GTT-CTTGTTGTTGTAGC-TGGTGTCCAAAGGTTATAT-TAAA [1280]
59ef08a5-aa1d-4d15-ba6f-1a925971  -----CATGATCATAATAGTTTTCTTGTTG-AATAG-GTT-CTTGTTGTTGTAGC-TGGTGTCCAAAGGTTATAT-TAAA [1280]
e1cae4d8-4f39-4401-8d82-5c5edaf3  -----CATGATCATAATAGTTTTCTTGTTG-AATAG-GTT-CTTGTTGTTGTAGC-TGGTGTCCAAAGGTTATAT-TAAA [1280]
```

```
5f47af75-253d-48b0-8b3e-ae59a63d  -----CATGATCATAATAGTTTTATCTCTG-AATAG-GTT-CTTGTTGTTGTAGC-TGGTGTCCAAAGGTTATA--TCAA [1280]
44f1962a-fce6-436a-9ea0-ef4a969d  -----CATGATCATAATAGTTTTCTTGTTG-AATAG-GTT-CTTGTTGTTGTAGC-TGGTGTCCAAAGGTTATAT-TAAA [1280]
4544809e-f4f9-4ff0-8142-5a2a2bc9  -----CATCATCATAATAGTTTTCTTGTC--AATAG-GTT-CTTGTT---GTAGC-TGGTGTCCAAAGGTTATAT-TAAA [1280]
9faa5557-cdc1-466c-815f-fb18a15f  ------ATGATCATAATAGTTTTCTTG-TTGAATAG-GTT-CTTGTTGTTGTAGCTGGTGTCC-AAAGGTTATATT-AAA [1280]
ef1a361c-a80e-43f6-8e1f-aeafa681  -----CATGATCATAATAGTTTTCTTGTTG-AATAG-GTTC-TTGTTGTTGTAGCTGGTGTCCA-AAGGTTATATT-AAA [1280]

Cp_PARENT                         GC-TTTC-ATTCAAATCAATATTTTTAT-TTGGAATAAATTTAAAA-AGATT-TGA-AAAAG-AATAAATAGCAGATCCT [1360]
Ct_PARENT                         GC-TTTC-ATTCAAATCAATATTTTTAT-TTGGAATAAATTTAAAA-AGATT-TGA-AAAAG-AATAAATAGCAGATCCT [1360]
bdf46f5b-2833-4b26-892f-323dd4c3  GC-TTTC-ATTCAAATCAATATTTTTATTTG-GAATAAATT----AAAGAA-TTGA-AAAAG-AATAAATAGCAGATCCT [1360]
c40b234c-b999-47c3-93ca-016af61e  CT-TTTC-ATTCAAATCAATATTTTTATTTG-GAATAAATTTAAAA-AGAT-TTGA-AAAAGAATGAATAAGCAGATCCT [1360]
0d8edcbc-b350-4d00-afbc-fd78a1f6  GC-TTTC-ATTCAAATCAATATTTTTATTTG-GAATAAATTTAAAA-AGAT-TTGA-AAAAG-AATCAATCGCAGATCCT [1360]
9cb07723-65ed-4c96-ac19-4234ea98  GC-TTTC-ATTCAAATCAATATTTTTATTTG-GAATGGATTAAAAA--GAT-TTGA-AAAAGA-ATAAATAGCAGATCCT [1360]
ed18c856-4d5b-4622-9fe8-9b5740af  GC-TTTC-ATTCAAATCAATA-TTTTTATTTGGAATAAATTTAAAAAGA-T-TTGA-AAAAG-AATAAATAGCAG-TCCT [1360]
c344612d-f873-4743-818c-65979638  GC-TTTC-ATTCAAATCAATATTTTTATTTG-GAATAAATTTAAAAAGA-T-TTGA-AAAAG-AATAAATAGCAGATCCT [1360]
baaaf62e-d0ee-4c49-9d83-0313d3df  GC-TTTC-ATTCAAATCAATATTTTTATTTG-GAATAAATTTAAAAAGA-T-TTGA-AAAAG-AATAAATAGCAGATCCT [1360]
6771788a-69b9-4fd8-b4e0-073881fe  GC-TTCC-ATTCAAATCAATATTTTTATTTG-GAATAAATTTAAAAAGA-T-TTGA-AAAAG-AATAAATAGCAAGTCCT [1360]
a6666a5a-6885-427a-9881-430dad33  GC-TTTC-ATTCAAATCAATATTTTTATTTG-GAATAAATTTAAAAAGA-T-TTGA-AAAAG-AATAAATAGCAGATCCT [1360]
5be1c0db-7e16-4441-a2ab-f8107a5d  GC-TTTC-ATTCAAATCAATATTTTTATT---TCAAATAATTAAAAAGA-T-TTGA-AAAAG-AATAAATAGCAGTCCTT [1360]
99a0bd03-c448-4605-9def-185e9f2d  GC-TTTC-ATTCAAATCAATATTTTTATTTG-GAATAAATTTAAAAAGA-T-TTGA-AAAAG-AATAAATAGCAGATCCT [1360]
c37ace00-c1f8-46e8-bcdc-c3c33bcb  GC-TTTCGATTCAAATCAATATTTTTATTTG-GAATAAATTTAAAAAGA-T-TTGA-AAAAG-AATAAATAGCAGATCCT [1360]
cdc9d647-e2ac-442b-8435-6dd7b82f  GC-TTTC-ATTCAAATCAATATTTTTATTTG-GAATAAATTTAAAAAGA-T-TTGA-AAAAG-AATAAATAGCAGATCCT [1360]
bc10b467-59cf-4610-94fe-267549f8  GC-TTTC-ATTCAAATAGATATATTTGGAATAAA-------ATTAAAGA-G-TTGA-AAAAG-AATAAATAGCAGATCCT [1360]
c8c3f0cb-c211-4877-8f07-1765b113  GC-TTTC-ATTCAAATCAATTATTTTTATTTGGAATAAATTTAAAAAGA-T-TTGA-AAAAG-AATAAATAGCAAGTCCT [1360]
720a0247-e02e-400b-901f-673d7527  GC-TTTC-ATTCAAATCAATATTTTTTATTTGGA-ATAAATTTAAAAAA-T-TAAA-AA----AATAAATAGCAGATCCT [1360]
f3f8defb-2c03-44af-abc8-d94623f7  GC-TTTC-ATTCAAATCAATA-TTTTTATTTGGAATAAATTTAAAAAGAT--TTGA-AAAAG-AATAAATAGCAGATCCT [1360]
fc2a9b4c-2c5f-43ed-8ea7-f2af0743  GC-TTTC-ATTCAAATCAATA-TTTTTATTTGGAATGATTAAAGAATTGAA--------AAA-GATAAATAGCAGGATAG [1360]
477953c0-cb9c-400c-bf05-dfadb08e  GC-TTTC-ATTCAAATCAATA-TTTTTATTTTGAAATAATTTAAAAAGAT--TTGA-AAAAG-AATAAATAGCAGATCCT [1360]
28c5afa3-4ebf-4ffa-a71d-2171913e  GC-TTTC-ATTCAAATCAATA-TTTTTATTTGGAATAAATTTAAAAAGAT--TTGA-AAAAG-AATAAATAGCAGATCCT [1360]
52a4f968-3394-459b-9ee3-10eab4a7  GC-TTTC-ATTCAAATCAATA-TTTTTATTTGGAATAAATTTAAAAAGAT--TTGA-AAAAG-AATAAATAGCAGATCCT [1360]
d36d5471-42a7-440b-bd5e-48d77156  GC-TTTC-ATTCAAATCAATA-TTTT---ATCGAAATAATTAAAGAATTGA--------AAA-AAAATAAATCACAATCC [1360]
f7886442-23fe-473a-9ead-a2b8a2a7  GC-TTTC-ATTCAAATCAATA-TTTTTATTTGGAATAAATTTAAAAAGATT-TGA--AAAAG-AATAAATAGCAGATCCT [1360]
f4770134-2a4b-449d-a4f6-c99f3383  GC-TTTC-ATTCAAATCAATA-TTTTTATTT---AAATAATTAAAAAGATT-TGA--AAAAG-AATAAATAGCAGATCCT [1360]
4dd53951-3b2d-42af-82ef-399bb712  GC-TTTC-ATTCAAATCAATA-TTTTTATTTGGAATAAATTTAAAAAGATT-TGA--AAAAG-AATAAATAGCAGATCCT [1360]
81ea3bac-8208-46c7-9546-bbd55b7c  GC-TTTC-ATTCAAATCAATA-TTTTTATTTGGAATAAATTTAAAAAGATT-TGA--AAAAG-AATAAATAGCAGATCCT [1360]
1e66a4e2-42fd-4fa9-b001-9bb7f263  GC-TTTC-ATTCAAATCAATA-TTTTTATTTGGAATAAATTTAAAAAGATT-TGA--AAAAG-AATAAATAGCAGATCCT [1360]
b75fad9a-877e-4f52-968b-9fcbb7a4  GC-TTTC-ATTCAAATCAATA-TTTTTATTTGGAATAAATTTAAAAAGATT-TGA--AAAAG-AATAAATAGCAGATCCT [1360]
f512b649-4f14-4eb6-b5a2-b0543441  GCTTTTC-ATTCAAATCAATA-TTTTTATTTGGAATAAATTTAAAAAGATT-TGA--AAAAG-AATAAATAGCAGATCCT [1360]
5021e737-de02-42f2-920d-18868672  GC-TTTC-ATTCAAATCAATA-TTTTTATTTGGAATAAATTTAAAAAGATT-TGA--AAAAG-AATAAATAGCAGATCCT [1360]
a46adbf5-3ba1-48b8-a762-2403bda7  GC--TTC-ATTCAAATCAATA-TTTTTATTTGGAATAAATTTAAAAAGATT-TGAA-A-AAG-AATAAATAGCAGATCCT [1360]
088a7e31-4c6b-4ff7-a19b-ff14531f  GC-TTTC-ATTCAAATCAATA-TTTTTATTTGGAATAAATTTAAAAAGATT-TGAA-A-AAG-AATGAATCGCAGATCCT [1360]
e82639d1-c451-4263-90a8-2b1ce6a1  GC-TTTC-ATTCAAATCAATA-TTTTTATTTGGAATAAATTTAAAAAGATT-TGAA-AAAG--AATAAATAGCAGATCCT [1360]
3cca4221-4eee-44ed-9fb4-ebbac961  GC-TTTC-ATTCAAATCAATA-TTTTTATTTGGAATAAATTTAAAAAGAT--TTGA-AAAAG-AATAAATAGCAGATCCT [1360]
d7a80796-8741-4878-b6ca-23aa66f7  GC-TTTC-ATTCAAATCAATA-TTTTTATTTGGAATAAATTTAAAAAGAT--TTGA-AAAAG-AATAAATAGCAGATCCT [1360]
421e3eda-83a0-4ebd-972c-f24667ee  GC-TTTC-ATTCAAATCAATA-TTTTTATTTGGAATAAATTTAAAAAGATT-TTGA-AAAAG-AATAAATAGCAGATCCT [1360]
e6a39029-4e34-4457-b2a9-0cf6f207  GC-TTTC-ATTCAAATCAATA-TTTTTATTTGGAATAAATTTAAAAAGAT--TTGA-AAAAG-AATAAATAGCAGATCCT [1360]
```

```
a1507163-34c9-4b24-b21e-3b5dfc54  GC-TTTC-ATTCAAATCAATATTTT-TATTTGGAATGA--TTTAAAAGA-AT-TGAAAAAGG-ATCAAATAGCAAGTCCT [1360]
51b01061-aaf5-4585-ba72-1a83671b  GC-TTTC-ATTCAAATCAATATTTT-TATTTGGAATAAATTTAAAAAGA-TT-TGA-AAAAG-AATAAATAGCAGATCCT [1360]
85703d1f-2c69-4541-ab66-acb7b838  ACTTTTC-ATTCAAATCAATATTTT-TATTTGGAATAAATTTAAAAAGA-TT-TGA-AAAAG-AATAAATAGCAAATTAT [1360]
ed39aa3b-d227-4de1-92bd-5ace4420  GC-TTTC-ATTCAAATCAATATTTT-TATTTGGAATAAATTTAAAAAGA-TT-TTGAAAAAG-AATAAATAGCAGATCCT [1360]
40e2af31-875d-446e-90a0-6e16a943  GC-TTTC-ATTCAAATCAATATTTT-TATTTGGAATAAATTTAAAAAGA-TT-T--GAAAAA-AATAAATAGCAGATCCT [1360]
873c76b4-8a29-4846-98df-cb5abcc9  GC-TTTC-ATTCAAATCAATATTTTTAT-TTGGAATAAATTTAAAAAGA-TT-TGA-AAAAA-AATAAATGGCCGATCCT [1360]
59ef08a5-aa1d-4d15-ba6f-1a925971  GC-TTTC-ATTCAAATCAATATTTTTAT-TTGGAATAAATTTAAAAAGA--T-TGA-AAAAG-AATAAATAGCAGATCCT [1360]
e1cae4d8-4f39-4401-8d82-5c5edaf3  GC-TTTC-ATTCAAATCAATATTTTTAT-TTGGAATAAATTTAAAAAGA-TT-TGA-AAAAG-AATAAATAGCAGATCCT [1360]
5f47af75-253d-48b0-8b3e-ae59a63d  GC-T--C-ATTCAAATCAATATTTTTAT-TTGGAATAAATTTAAAAAGA-TT-TGA-AAAAG-AATAAATAGCAGATCCT [1360]
44f1962a-fce6-436a-9ea0-ef4a969d  GC-TTTC-ATTCAAATCAATATTTTTAT-TTGGAATTGATTTAAAAAGA-TT-TGA-AAAAG-AATAAATAGCAGATCCT [1360]
4544809e-f4f9-4ff0-8142-5a2a2bc9  GC-TTTC-ATTCAAATCAATATTTTTAT-TTGGAATAAATTTAAAAAGA-TT-TGA-AAAAG-AATAAATAGCAAGTGCT [1360]
9faa5557-cdc1-466c-815f-fb18a15f  GC-TTTC-ATTCAAATCAATATTTTTAT-TTGGAATGATTTA---AAAAGAT-TCA-AAAAG-AATAAATAGCAGATCCT [1360]
ef1a361c-a80e-43f6-8e1f-aeafa681  GC-TTTC-ATTCAAATCAATATTTTTATTTG-GAATAAATTTAAAA-AGAT-TTGA-AAAAGA-ATAAATAGCAGATCCT [1360]

Cp_PARENT                         CC-AATTGTC-ATAACCAAT----CCCAAA---ATAGTTG--AAAATTTAAGC-TCT-TCGTTATTTAATA---A-A-AC [1440]
Ct_PARENT                         CC-AATTGTC-ATAACCAAT----CCCAAA---ATAGTTG--AAAATTTAAGC-TCT-TCGTTATTTAATA---A-A-AC [1440]
bdf46f5b-2833-4b26-892f-323dd4c3  CC-AATTGTC-ATAACCAAT----CCCAAA---ATAGTTG--AAAATTTAAG-CTCT-TGGTTCTTTAATCAAACTT-CC [1440]
c40b234c-b999-47c3-93ca-016af61e  CC-AATTGTC-ATAACCAAT----CCCAAA---ATAGTTG--AAAATTTAAG-CTCT-TCGTTATTTAATAAAAC-T-CC [1440]
0d8edcbc-b350-4d00-afbc-fd78a1f6  CC-AATTGTC-ATAACCAAT----CCCAAA---ATCGTTG--AAAATTTAAG-CTCT-TCGTTATTTAATAAAAC-T-CC [1440]
9cb07723-65ed-4c96-ac19-4234ea98  CC-AATTGTC-ATAACCAAT----CCCAAA---ATAGTTG--AAAATTTAAG-CTCT-TCGTTATTTAATAAAAC-T-CC [1440]
ed18c856-4d5b-4622-9fe8-9b5740af  CC-AATTGTC-ATAACCAAT---CCCCAAA---ATAGTTG--AAAATTTAAG-CTCT-TCGTTATTTAATAAAAC-T-CC [1440]
c344612d-f873-4743-818c-65979638  CC-AATTGTC-ATAACCAAT----CCCAAA---ATAGTTG--AAAATTTAAG-CTCT-CAGTTATTTGATCAAACTT-CC [1440]
baaaf62e-d0ee-4c49-9d83-0313d3df  CC-AATTGTC-ATAACCAAT----CCCAAA---ATAGTTG--AAAATTTAAG-CTCT-TCGTTATTTAATAAAAC-T-CC [1440]
6771788a-69b9-4fd8-b4e0-073881fe  CC-AATTGTC-ATAACCAAT----CCCAAA---ATAGT--TGAAAATTTAAG-CTCT-TCGTTATTTAATAAAAC-T-CC [1440]
a6666a5a-6885-427a-9881-430dad33  CC-AATTGTC-ATAACCAAT----CCCAAA---ATAGT--TGAAAATTTAAG-CTCT-TCGTTATTTAATAAAAC-T-CC [1440]
5be1c0db-7e16-4441-a2ab-f8107a5d  CC-AATTGTC-ATAACCAAT---CCCCAAA---ATAAGCTTGAAAATTTAAG-CTCT-TCGTTATTTAATAAAAC-T-CC [1440]
99a0bd03-c448-4605-9def-185e9f2d  CC-AATTGTC-ATAACCAAT----CCCAAA---ATAGTTG--AAAATTTAAG-CTCT-TCGTTATTTAATAAAAC-T-CC [1440]
c37ace00-c1f8-46e8-bcdc-c3c33bcb  CC-AATTGTC-ATAACCAAT----CCCAAA---ATAGTT--GAAAATTTAAG-CTCT-TCGTTATTTAATAAAAC-T-CC [1440]
cdc9d647-e2ac-442b-8435-6dd7b82f  CC-AATTGTC-ATAACCAAT----CCCAAA---ATAGTTC---AAACTTAAG-CTCT-TCGTTATTTAATAAAAC-T-CC [1440]
bc10b467-59cf-4610-94fe-267549f8  CC-AATTGTC-ATAACCAAT----CCCAAA---ATAGTTG--AAAATTTAAG-CTCT-TCGTTATTTAATAAAAC-T-CC [1440]
c8c3f0cb-c211-4877-8f07-1765b113  CC-AATTGTC-ATAACCAAT----CCCAAA---ATAGTTG--AAAATTTAAG-CTCT-TCGTTATTTAATAAAAC-T-CC [1440]
720a0247-e02e-400b-901f-673d7527  CC-AATTGTC-ATAACCAAT----CCCAAA---ATAGTTTTGAAAATTTAAG-CTCT-TCGTTATTTAATAAAAC-T-CC [1440]
f3f8defb-2c03-44af-abc8-d94623f7  CC-AATTGTC-ATAACCAAT---CCCAAAA---TAG---TTGAAAATT-TAAGCTCT-TCGTTATTTAATAAAAC-T-CC [1440]
fc2a9b4c-2c5f-43ed-8ea7-f2af0743  CC-AATTGTC-ATAACCAAT---CCCAAAA---TCA---TTGAAAATTTAAG-CTCT-TCATTATAAAATAAACT-C-CA [1440]
477953c0-cb9c-400c-bf05-dfadb08e  CC-AATTGTC-ATAACCAAT---CCCAAAA---TAG---TTGAAAATTTAAG-CTCT-TCGTTATTTAATAAAAC-T-CC [1440]
28c5afa3-4ebf-4ffa-a71d-2171913e  CC-AATTGTC-ATAACCAAT---CCCAAAA---TAG---TTGAAAATTTAAG-CTCT-TCGTTATTTAATAAAAT-T-CC [1440]
52a4f968-3394-459b-9ee3-10eab4a7  CC-AATTGTC-ATAACCAAT---CCCAAAA---TAG---TTGAAAATTTAAG-CTCT-TCGTTATTTAATAAAAC-T-CC [1440]
d36d5471-42a7-440b-bd5e-48d77156  TC-AATTGTC-ATAACCAAT---CCCAAAA---TCA---TTGAAAATTTAAG-CTCT-TCGTTATTTAATAAAAC-T-GC [1440]
f7886442-23fe-473a-9ead-a2b8a2a7  CC-AATTGTC-ATAACCAAT---CCCAAAA---TAG----TTAAAATTTAAG-CTCT-TGGTTATTTAACAAAAC-T-CC [1440]
f4770134-2a4b-449d-a4f6-c99f3383  CC-AATTGTC-ATAACCAAT---CCCAAAA---TAG---TTGAAAATT--AAGCTCT-TCGTTATTTAATAAAAC-T-CC [1440]
4dd53951-3b2d-42af-82ef-399bb712  CC-AATTGTC-ATAACCAAT---CCCAAAA---TAG---TTGAAAATTTGAC-CTCT-TCGTTATTTAATAAAAC-T-CC [1440]
81ea3bac-8208-46c7-9546-bbd55b7c  CC-AATTGTC-ATAACCAAT---CCCAAAA---TAG---TTGAAAATTTAAG-CTCT-TCGTTATTTAATAAAAC-T-CC [1440]
1e66a4e2-42fd-4fa9-b001-9bb7f263  CC-AATTGTC-ATAACCAAT---CCCAAAA---TAG---TTGAAAATTTAAG-CTCT-TCGTTATTTAATAAAAC-T-CC [1440]
b75fad9a-877e-4f52-968b-9fcbb7a4  CC-AATTGTC-ATAACCAAT---CCCAAAA---TAG---TTGAAAATTTAAG-CTCT-TCGTTATTTAATAAACT-C-CA [1440]
f512b649-4f14-4eb6-b5a2-b0543441  CC-AATTGTC-ATAACCAAT---CCCAAAA---TAG---TTGAAAATTTAAG-CTCT-TCGTTATTTAATAAAAC-T-CC [1440]
```

```
5021e737-de02-42f2-920d-18868672  CC-AATTGTC-ATAACCAAT---CCCAAAA---TAG---TTGAAAATCTAAG-CTCT-TCGTTATTTAATAAAAC-T-C- [1440]
a46adbf5-3ba1-48b8-a762-2403bda7  CC-AATTGTC-ATAACCAAT---CCCAAAA---TAG---TTGAAAATTTTAAGCTCT-TCGTTATTTAATAAAAC-T-CC [1440]
088a7e31-4c6b-4ff7-a19b-ff14531f  CC-AATTGTC-ATAACCAAT---CCCAAAA---TAG---TTGAAAATTTAAG-CTCT-TCGTTATTTAATAAAAC-T-CC [1440]
e82639d1-c451-4263-90a8-2b1ce6a1  CC-AATTGTC-ATAACCAAT---CCCAAAA---TAG---TTGAAAATTTAAGC-----TCTTCCATTTAATAAAC-T-CC [1440]
3cca4221-4eee-44ed-9fb4-ebbac961  CC-AATTGTC-ATAACCAAT---CCCAAAA---TAG---TTGAAAATTTAAG-CTCT-TCGTTATTTAATAAAAC-T-CC [1440]
d7a80796-8741-4878-b6ca-23aa66f7  CC-AATTGTC-ATAACCAAT---CCCAAAA---TAG---TTGAAAATTTGAGCTTCT-TCGTTATTTAATAAAAC-T-CC [1440]
421e3eda-83a0-4ebd-972c-f24667ee  CC-AATTGTC-ATAACCAAT---CCCAAAA---TAG---TTGAAAATTTAAG-CTCT-TCGTTATTTAATAAAAC-T-CC [1440]
e6a39029-4e34-4457-b2a9-0cf6f207  CC-AATTGTC-ATAACCAAT---CCCAAAA---TAG---TTGAAAATTTAAG-CTCT-TCGTTATTTAATAAAAT-T-CC [1440]
a1507163-34c9-4b24-b21e-3b5dfc54  CCA-ATTGTC-ATAACC--AATCC--CAAA---ATAGTTG--AAAATTTAAGC-TCT-TCGTTATTTAATAAAAC-T-CC [1440]
51b01061-aaf5-4585-ba72-1a83671b  GCA-ATTGTC-ATAACC--AATCC--CAAA---ATCGTTG--AAAATTTA-GC-TTC-TTTTTGGTTATTAATCG-T-CC [1440]
85703d1f-2c69-4541-ab66-acb7b838  CCA-ATTGTC-ATAACC--AATCC--CAAA---ATAGTTG--AAAATTTAAGC-TCT-TCGTTATTTAATAAAAC-T-CC [1440]
ed39aa3b-d227-4de1-92bd-5ace4420  CCA-ATTGTC-ATAACC--AATCC--CAAA---ATAGTTG--AAAATTTAAGC-TCT-TCGTTATTTAATAAAAC-T-CC [1440]
40e2af31-875d-446e-90a0-6e16a943  CCA-ATTGTC-ATAACC--AATCC--CAAA---ATAGTTG--AAAATTTAAGC-TCT-TCGTTATTTAATAAAAC-T-CC [1440]
873c76b4-8a29-4846-98df-cb5abcc9  CC-AATTGTC-ATAACC--AATACCCCAAA---ATAGTTG--AAAATTTAAGC-TCT-TCGTTATTTAATAAAAC-T-CC [1440]
59ef08a5-aa1d-4d15-ba6f-1a925971  CC-AATTGTC-ATAACC--AATCC--CAAA---ATAGTTG--AAAATTTAAGC-TCT-TCGTTATTTAATAAAAC-T-AA [1440]
e1cae4d8-4f39-4401-8d82-5c5edaf3  CC-AATTGTC-ATAACC--AATCC--CAAA---ATAGTTG--AAAATTTAAGC-TCT-TCGTTATTTAATAAAAC-T-CC [1440]
5f47af75-253d-48b0-8b3e-ae59a63d  CC-AATTGTC-ATAACC--AATCC--CAAA---ATAGTTG--AAAATTTAAGC-TCT-TCGTTATTTAATAAAAC-T-CC [1440]
44f1962a-fce6-436a-9ea0-ef4a969d  CC-AATTGTC-ATAACC--AATCC--AAAA---ATAGTTG--AAAATTTAAGC-TCT-TCGTTATTTAATAAAAC-T-CC [1440]
4544809e-f4f9-4ff0-8142-5a2a2bc9  CCAAATTGTC-ATAACC--AATAC--CCAAAATAACATTG--AAATTTTGAGCTTCT-TGGTTCTTTAATAAAAC-T-CC [1440]
9faa5557-cdc1-466c-815f-fb18a15f  CC-AATT-GTCATAACC---AATCCCAAA----ATAGTTG--AAAATTTAAGC-TCTT-CGTTATTTAATAAAAC-T-CC [1440]
ef1a361c-a80e-43f6-8e1f-aeafa681  CC-AATTGTC-ATAACCAAT----CCCAAA---ATCG-TG--AAAATTTAAG-CTCT-TCGTTATTTAATAAAAC-T-CC [1440]

Cp_PARENT                         TCCA—ATTAAAGT---TTGA-GC-TCCAGAT-TTAACA-TAG-CCTGT-TAGATTATA----AGCTATA-GGAGATGTAG [1520]
Ct_PARENT                         TCCA—ATTAAAGT---TTGA-GC-TGCAGAT TTAACA-TAG-CCTGT-TAGATTATA----AGCTATA-GGAGATGTAG [1520]
bdf46f5b-2833-4b26-892f-323dd4c3  AATT--AAAGT-----TTGA-GC-TGCAGAT-TTAACA-TAG-CCTGTT--CAT--TA-T-AAGCTATA-GGAGATGTAG [1520]
c40b234c-b999-47c3-93ca-016af61e  AATT--AAAGT-----TTGA-GC-TGCAGAT-TACAAC-ATA-GCCTGTTAGAT--TA-T-AAGCTATA-GGAGATGTAG [1520]
0d8edcbc-b350-4d00-afbc-fd78a1f6  AATT--AAAGT-----TTGA-GC-TGCAGAT-TTAACA-TAG-CCTGT-TAGAT--TA-T-AAGCTATA-GGAGATGTAG [1520]
9cb07723-65ed-4c96-ac19-4234ea98  AATT--AAAGT-----TTGA-GC-TGCAGAT-TTAACA-TAG-CCTGT-TAGAT--TA-T-AAGCTATA-GGAGATGTAG [1520]
ed18c856-4d5b-4622-9fe8-9b5740af  AATT--CAAGA-----TTTA-AC-TGCCAAT-TTCACA-TTA-GCCTG-TAGAT--TA-T-AAGCTATA-GGAGATGTAG [1520]
c344612d-f873-4743-818c-65979638  AATT--AAAGT-----TTGA-GC-TGCAG---TTAACA-TAG-CCTGT-TAGAT--TA-T-AAGCTATA-GGAGATGTAG [1520]
baaaf62e-d0ee-4c49-9d83-0313d3df  AATT--AAAGT-----TTGA-GC-TGCAGAT-TTAACA-TAG-CCTGT-TAGAT--TA-T-AAGCTATA-GGAGATGTAG [1520]
6771788a-69b9-4fd8-b4e0-073881fe  AAAT--TAAAC-----TTTA-AGTTACAGTT-GGCATT-TAG-CCTCT-TAGAT--TA-T-AAGCTATAGGGAGATGTAG [1520]
a6666a5a-6885-427a-9881-430dad33  AATT--AAAGT-----TTGA-GC-TGCAG-T-TTAACA-TCA-CCTG--TAGAT--TA-T-AAGCTATA-GGAGATGTAG [1520]
5be1c0db-7e16-4441-a2ab-f8107a5d  AATT--AAAGT-----TTGA-GCTGCCAGAT-TTAACA-TAG-CCTGT-TAGAT--TA-T-AAGCTATA-GGAGATGTAG [1520]
99a0bd03-c448-4605-9def-185e9f2d  AATT--AAAGT-----TTGA-GC-TGCAGAT-TTAACA-TAG-CCTGT-TAGAT--TA-T-AAGCTATA-GGAGATGTAG [1520]
c37ace00-c1f8-46e8-bcdc-c3c33bcb  AATT--AAAGT-----TTGA-GCTTGCAGAT-TTAACA-TAG-CCTGT-TAGAT--TA-T-AAGCTATA-GGAGATGTAG [1520]
cdc9d647-e2ac-442b-8435-6dd7b82f  AATT--AAAGT-----TTGA-GC-TGCAGAT-TTAACA-TAG-CCTGT-TAGAT--TA-T-AAGCTATA-GGAGATGTAG [1520]
bc10b467-59cf-4610-94fe-267549f8  AATT--AAAGT-----TTGA-GC-TGCAGAT-TTAACA-TAG-CCTGT-TAGAT--TA-T-AAGCTATA-GGAGATGTAG [1520]
c8c3f0cb-c211-4877-8f07-1765b113  AATT--AAAGT-----TTGA-GC-TGCAGAT-TTAACA-TAG-CCTGT-TAGAT--TA-T-AAGCTATA-GGAGATGTAG [1520]
720a0247-e02e-400b-901f-673d7527  AATT--AAAGT-----TTGA-GC-TGCAGAT-TTAACA-TAG-CCTGT-TAGAT--TA-T-AAGCTATA-GGAGATGTAG [1520]
f3f8defb-2c03-44af-abc8-d94623f7  AATT--AAAGT-----TTGA-GC-TGCAGAT-TTAACA-TCG-CCTCT-TAGAT--TA-T-AAGCTATA-GGAGATGTAG [1520]
fc2a9b4c-2c5f-43ed-8ea7-f2af0743  ATTG--AAAGT-----TTGA-GC-TGCAGAG-TTGACATTAG-CCTGT-TAGAT--TA-T-AAGCTATA-GGAGATGTAG [1520]
477953c0-cb9c-400c-bf05-dfadb08e  AATT--AAAGT-----TTGA-GC-T--AGAT-TTAACA-TAG-CCTGT-TAGAT--TA-T-AAGCTATA-GGAGATGTAG [1520]
28c5afa3-4ebf-4ffa-a71d-2171913e  AATT--AAAGT-----TTGA-GC-TGCAGAT-TTAACA-TAG-CCTGT--AGAT--TA-T-AAGCTATA-GGAGATGTAG [1520]
52a4f968-3394-459b-9ee3-10eab4a7  AATT--AAAGT-----TTCA-GC-TGCAGAT-TTAACA-TAG-CCTGT-TAGAT--TA-T-AAGCTATG-GGGAATGTTC [1520]
```

```
d36d5471-42a7-440b-bd5e-48d77156  GATT--GAGAA-----TTTT-GA-GCTAAAT-TTGACA-TTC-ACTGC-TAGAT--TA-T-AAGCTATA-GGAGATGTAG [1520]
f7886442-23fe-473a-9ead-a2b8a2a7  AATT--AAAGT-----TTGA-GC-T--AGAT-TTAACA-TAG-CCTGT-TAGAT--TA-T-AAGCTATA-GGAGATGTAG [1520]
f4770134-2a4b-449d-a4f6-c99f3383  AATT--AAAGT-----TTGA-GC-TGCAGAT-TTAACA-TAG-CCTGT-TAGAT--TA-T-AAGCTATA-GGAGATGTAG [1520]
4dd53951-3b2d-42af-82ef-399bb712  AATT--AAAGT-----TTGA-GC-TGCAGAT-TTAACA-TAG-CCTGT-TAGAT--TA-T-AAGCTATA-GGAGATGTAG [1520]
81ea3bac-8208-46c7-9546-bbd55b7c  AATT--AAAGT-----TTGA-GC-TGCAGAT-TTAACA-TAG-CCTGT-TAGAT--TA-T-AAGCTATA-GGGAGTGTAG [1520]
1e66a4e2-42fd-4fa9-b001-9bb7f263  AATT--AAAGT-----TTGA-GC-TGCAGAT-TTAACA-TAG-CCTGT-TAGAT--TA-T-AAGCTATA-GGAGATGTAG [1520]
b75fad9a-877e-4f52-968b-9fcbb7a4  -ATT--AAAGT-----TTGA-GC-TAGA--T-TTAACA-TAG-CCTGT-TAGATTATA-A-GCTATATA-GGAGATGTAG [1520]
f512b649-4f14-4eb6-b5a2-b0543441  AATT--AAAGT-----TTGA--C-TGCAGAT-TTAACA-TAG-CCTGT-TAGAT--TA-T-AAGCTATA-GGAGATGTAG [1520]
5021e737-de02-42f2-920d-18868672  -CAT--AAATT-----TTGA-GC-TGCAGAT-TTAACA-TAG-CCTGT-TAGAT--TA-T-AAGCTATA-GGAGATGTAG [1520]
a46adbf5-3ba1-48b8-a762-2403bda7  AATT--AAAGT-----TTGA-GC-TGCAGAT-TTAACA-TAG-CCTGT-TAGAT--TA-T-AAGCTATA-GGAGATGTAG [1520]
088a7e31-4c6b-4ff7-a19b-ff14531f  AATT--AAAGT-----TTGA-GC-TGCAGAT-TTAACA-TAG-CCTGT-TCATT--TA-T-AAGCTATA-GGAGATGTAG [1520]
e82639d1-c451-4263-90a8-2b1ce6a1  AATT--AAAGT-----TTGA-GC-TGCAGAT-TTAACA-TAG-CCTGT-TAGAT--TA-T-CAGCTATA-GGAGATGTAG [1520]
3cca4221-4eee-44ed-9fb4-ebbac961  AATT--AAAGT-----TTGA-GC-TGCAGAT-TTAACA-TAG-CCTGT-TAGAT--TA-T-AAGCTATA-GGAGATGTAG [1520]
d7a80796-8741-4878-b6ca-23aa66f7  AATT--AAAGT-----TTGA-GC-TGCAGAT-TTAACA-TAT-CACCT-ATAAT--TA-T-GGC--TCT-GGAGATGTAG [1520]
421e3eda-83a0-4ebd-972c-f24667ee  AATT--AAAGT-----TTGA-GC-TGCAGAT-TTAACA-TAG-CCTGT-TAGAT--TA-T-AAGCTATA-GGAGATGTAG [1520]
e6a39029-4e34-4457-b2a9-0cf6f207  AATT--AAAGT-----TTGA-GC-TGCAGAT-TTAACA-TAG-CCTGT-TAGAT--TA-T-AAGCTATA-GGAGATGTAG [1520]
a1507163-34c9-4b24-b21e-3b5dfc54  AATT--AAAGT-----TTGA-GC-TGCAGAT-TTAACA-TAG-CCTGT-TAGAT--TA-T-A-GCTATA-GGAGATGTAG [1520]
51b01061-aaf5-4585-ba72-1a83671b  AATT--AAAGT-----TTGA-GC-TGCAGTTTA---GT-CAC-CTATT-TAGAT--TA-T-AAGCTATA-GGAGATGTAG [1520]
85703d1f-2c69-4541-ab66-acb7b838  AATT--AAAGT-----TTGA-GC-TGCAAGA-TTAACA-TAG-CCTGT-TAGAT--TA-T-AAGCTATA-GGAGATGTAG [1520]
ed39aa3b-d227-4de1-92bd-5ace4420  AATT--AAAGT-----TTGA-GC-TGCAGAT-TTAACA-TAG-CCTG--TAGAT--TA-T-AAGCTAT--AGGAGATGTA [1520]
40e2af31-875d-446e-90a0-6e16a943  GTGC--CAAGT-----TTCA-GC-TGCAGAT-TTGACA-TAG-CCTGT-TAGAT--TA-T-AAGCTAT--CCAGGAGATG [1520]
873c76b4-8a29-4846-98df-cb5abcc9  AATT--AAAGT-----TTGA-GC-TGCAGAT-TTAACA-TAG-CCTGT-TAGAT--TA-T-AAGCTATA-GGAGATGTAG [1520]
59ef08a5-aa1d-4d15-ba6f-1a925971  AAT----AAAT-----TTTA-AC-TGCAGAT-TTAACA-TAG-CCTGT-TAGAT--TA-T-AAGCTATA-GGAGATGTAG [1520]
e1cae4d8-4f39-4401-8d82-5c5edaf3  AATT--AAAGT-----TTGA-GC-TGCAGAT-TTAACA-TAG-CCTGT-TAGAT--TA-T-AAGCTATA-GGAGATGTAG [1520]
5f47af75-253d-48b0-8b3e-ae59a63d  AATT--AAAGT-----TTGA-GC-T--AGAT-TTAACA-TAG-CCTGT-TAGAT--TA-T-GGCTCTAA-GGAGATGTGG [1520]
44f1962a-fce6-436a-9ea0-ef4a969d  AATT--AAAGT-----TTGA-GC-TGCAGAT-TTAACA-TAG-CCTGT-TAGAT--TA-T-AAGCTATA-GGAGATGTAG [1520]
4544809e-f4f9-4ff0-8142-5a2a2bc9  AATT--AAAGT-----TTGA-GC-TGCAAGA-TTAACA-TAGCCCTGT-TAGAT--TA-T-AAGCTATA-GGAGATGTAG [1520]
9faa5557-cdc1-466c-815f-fb18a15f  AATT--AAGAAAAATTTG---CAGCTGCAGATTTAACA-TAG-CCTGT-TAGATTATA----AGCTATA-GGAGATGTAG [1520]
ef1a361c-a80e-43f6-8e1f-aeafa681  AATT--AAAGT-----TTGA-GC-TGCAAGT-TTGACA-TCA-CCTGTTTCAAT--TA-T-AAGCTATA-GGAGATGTAG [1520]

Cp_PARENT                         TA-TGAATACACCATATT-ATAA-TTATCCC-----AAGGAAGAA-ATTCAATATACCACTAATTAAAAT---A-G--GG [1600]
Ct_PARENT                         TA-TGAATACACCATATT-ATAA-TTATCCC-----GAGGAAGAA-ATTCAATATACCACTAATTAAAAT---A-G--GA [1600]
bdf46f5b-2833-4b26-892f-323dd4c3  TA-TGAATACACCATATT-ATAA-TTATCCC-----GAGGAAGAA-ATTCAATATACCACTAATTAAAAT---A-G--GA [1600]
c40b234c-b999-47c3-93ca-016af61e  TA-TGAATACACCATATT-ATCA-TTACTCCC----GAGGAAGAA-ATTCAATATACCACTAATTAAAAT---A-G--GA [1600]
0d8edcbc-b350-4d00-afbc-fd78a1f6  TA-TGAATACACCATATT-ATAA-TTATCCCC----GAGGAAGAA-ATTCAATATACCACTAATTAAAAT---A-G--GA [1600]
9cb07723-65ed-4c96-ac19-4234ea98  TA-TGAATACACCATATT-ATAA-TTATCCCGAG--GA---AGAG-ATTCAATATACCACTAATTAAAAT---A-G--GA [1600]
ed18c856-4d5b-4622-9fe8-9b5740af  TA-TGAATACACCATATT-ATAA-TTGTGC---A--G----AGAA-ATTCAATATACCACTAATTCGA-----T-G--GA [1600]
c344612d-f873-4743-818c-65979638  TA-TGAATACACCATATT-ATAA-TTATCC---C--GAGGAAGAA-ATTCAATATACCACTAATTAAAAT---T-G--GA [1600]
baaaf62e-d0ee-4c49-9d83-0313d3df  TA-TGAATACACCATATT-ATAA-TTCTGC-------AAGGAAAA-AATCAATATACCACTAATTAAAAT---A-G--GA [1600]
6771788a-69b9-4fd8-b4e0-073881fe  TA-TGAATACACCATATT-TTG--TAAATTATGC--AAGGAAGAA-ATTCAATATACCACTAATTAAAAT---A-G--GA [1600]
a6666a5a-6885-427a-9881-430dad33  TA-TGAATACACCATATT-ATAA-TTATCC---C--GAGGAAGAA-ATTCAATATACCACTAATTAAAAT---A-G--GA [1600]
5be1c0db-7e16-4441-a2ab-f8107a5d  TA-TCAATACACCATATT-CTAA-TTTATC--CC--GAGGAAGAA-ATTCAATATACCACTAATTAAAAT---A-G--GA [1600]
99a0bd03-c448-4605-9def-185e9f2d  TA-TGAATACACCATATT-ATAA-TTATCC---C--AAGGAAGAA-ATTCAATATACCACTAATTAAAAT---A-G--GA [1600]
c37ace00-c1f8-46e8-bcdc-c3c33bcb  TA-TGAATACACCATATT-ATAA-TTAT-C---C--GAGGAAGAA-ATTCAATATACCACTAATTAAAAT---A-G--GA [1600]
cdc9d647-e2ac-442b-8435-6dd7b82f  TA-TGAATACACCATATT-ATAA-TTATCC---C--GGGGAAGAA-ATTCAATATACCACTAATTAAAAT---A-G--GA [1600]
```

```
bc10b467-59cf-4610-94fe-267549f8  TA-TGAATACACCATATT-ATAA-TTATCC---C--G---AAGAA-ATTCAATATACCACTAATTAAAAT---A-G--GA [1600]
c8c3f0cb-c211-4877-8f07-1765b113  TA-TGAATACACCATATT-ATAA-TTATCC---C--GAGGAAGAA-ATTCAATATACCACTAATTAAAAT---A-G--GA [1600]
720a0247-e02e-400b-901f-673d7527  TA-TGAATACACCATATT-ATGA-TTATTG---C--GAGGAAGAA-ATTCAATATACCACTAATTAAAAT---A-G--GA [1600]
f3f8defb-2c03-44af-abc8-d94623f7  TA-TGAATACACCATATTCCATAATTATCC---C--GAGGAAGAA-ATTCAATATACCACTAATTAAAAT---A-G--GA [1600]
fc2a9b4c-2c5f-43ed-8ea7-f2af0743  TA-TGAATACACCATC-T-ATAA-TTATCC---C--GAGGAAGAA-ATTGAATATACCACTAATTAAAAT--------CA [1600]
477953c0-cb9c-400c-bf05-dfadb08e  TA-TGAATACACCATATT-ATAA------T---T--ATGCAAGAA-ATTCAATATACCACTAATTAAAAT---A-G--GA [1600]
28c5afa3-4ebf-4ffa-a71d-2171913e  TA-TGAATACACCATATT-ATAA-TTAT-C---C--GAGGAAGAA-ATTCAATATACCACTAATTAAAAT---A-G--GA [1600]
52a4f968-3394-459b-9ee3-10eab4a7  -A-TGAATACACCATATT-ATAA-TTATCC---C--GAGGAAGAA-ATTCAATATACCACTAATTAAAAT---A-G--GA [1600]
d36d5471-42a7-440b-bd5e-48d77156  TA-TGAATACACCATATT-ATAA-TTATAC------AAGGAAGAA-ATTCAATATACCACTAATTGAATA---T-G--GA [1600]
f7886442-23fe-473a-9ead-a2b8a2a7  TA-TGAATACACCATATT-ATAA-TTCTGC---G--GAGGAAGAA-ATTCAATATACCACTAATTAAAAT---A-G--GA [1600]
f4770134-2a4b-449d-a4f6-c99f3383  TA-TGAATACACCATATT-ATAATTATGCA---A--G------AA-ATTCAATATACCACTAATTAAAAT---A-G--GA [1600]
4dd53951-3b2d-42af-82ef-399bb712  TA-TGAATACACCATATT-ATAA-TTATCC---C--GAGGAAGAA-ATTCAATATACCACTAATTAAAAT---A-G--GA [1600]
81ea3bac-8208-46c7-9546-bbd55b7c  TA-TGAATACACCATATT-GTAGTTTCTCC---C--GAGGAAGAA-ATTCAATATACCACTAATTAAAAT---A-G--GA [1600]
1e66a4e2-42fd-4fa9-b001-9bb7f263  TA-TGAATACACCATATT-ATAA-TTATCC---C--GAGGAAGAA-ATTCAATATACCACTAATTAAAAT---A-G--GA [1600]
b75fad9a-877e-4f52-968b-9fcbb7a4  TA-TGAATACACCATATT-ATAA-TTATCC---C--AAGGAAGAA-ATTCAATATACCACTAATTAAAAT---A-G--GA [1600]
f512b649-4f14-4eb6-b5a2-b0543441  TA-TGAATACACCATATT-ATAA-TTATCC---C--GAGGAAGAA-ATTCAATATACCACTAATTAAAAT---A-G--GA [1600]
5021e737-de02-42f2-920d-18868672  TA-TGAATACACCATATT-ATAA-TTATCC---C--GAGGAAGAA-ATTCAATATACCACTAATTAAAAT---A-G--GA [1600]
a46adbf5-3ba1-48b8-a762-2403bda7  TA-TGAATACACCATATT-ATAA-TTATCC---C--GAGGAAGAA-ATTCAATATACCACTAATTAAAATCACG-G--AC [1600]
088a7e31-4c6b-4ff7-a19b-ff14531f  TA-TGAATACACCATATT-ATAA-TTATCC---C--GAGGAAGAA-ATTCAATATACCACTAATTAAAAT--CA-G--AC [1600]
e82639d1-c451-4263-90a8-2b1ce6a1  TA-TGAATACACCATATT-ATAA-TTATCC---C--GAGGAAGAA-ATTCAATATACCACTAATTAAAAT--AG-G--AC [1600]
3cca4221-4eee-44ed-9fb4-ebbac961  TA-TGAATACACCATATT-ATAA-TTATCC---C--GAGGAAGAA-ATTCAATATACCACTAATTAAAAT--AG-G--AC [1600]
d7a80796-8741-4878-b6ca-23aa66f7  TA-TGAATACACCATATT-ATAA-TTACGA---G-----GAAGAA-AT----TCAATATACCATTAAAAT--AG-G--AC [1600]
421e3eda-83a0-4ebd-972c-f24667ee  TA-TGAATACACCATATT-ATAA-TTATCC---C--AAG----AA-ATTCAATATACCACTAATTAAAAT--AG-G--AC [1600]
e6a39029-4e34-4457-b2a9-0cf6f207  TA-TGAATACACCATATT-ATAA-TTATCC---C--GAGGAAGAA-ATTCAATATACCACTAATTAAAAT--AG-G--AC [1600]
a1507163-34c9-4b24-b21e-3b5dfc54  TA-TGAATACACCA---T-ATAA-TTATCC---C--GAGGAAGAA-ATTCAATATACCACTAATTAAAAT---A-G--GA [1600]
51b01061-aaf5-4585-ba72-1a83671b  TA-TAAATACACCATATT-ATAA-TTATCC---C--GAGGAAGAA-ATTCAATTATCCACTAATTAAAAT---A-G--GA [1600]
85703d1f-2c69-4541-ab66-acb7b838  TA-TGAATACACCATATT-ATAA-TTATCC---C--GAGGAAGAA-ATTCAATATACCACTAATTAAAAT---A-A--GA [1600]
ed39aa3b-d227-4de1-92bd-5ace4420  GT-ATAATACACCATATT-ATAA-TTATCC---C--GAGGAAGAA-ATTCAATATACCACTAATTAAAAT---A-G--GA [1600]
40e2af31-875d-446e-90a0-6e16a943  TA-TGAATACACCATATT-ATAA-TTAT-C---C--GAGGAAGAA-ATTCAATATACCACTAATTAAAAT---A-G--GA [1600]
873c76b4-8a29-4846-98df-cb5abcc9  TA-TGAATACACCATATT-ATAA-TTATCC---C--GAGGAAGAA-ATTCAATATACCACTAATTAAAAT---A-G--GA [1600]
59ef08a5-aa1d-4d15-ba6f-1a925971  TA-TGAATACACCATATT-ATAA-TTATCC---C--GAGGAAGAA-ATTCAATATACCACTAATTAAAAT---A-G--GA [1600]
e1cae4d8-4f39-4401-8d82-5c5edaf3  TA-TGAATACACCATATT-ATAA-TTATCC------GAGGAAGAA-ATTCAATATACCACTAATTAAAAT---G-GG-GC [1600]
5f47af75-253d-48b0-8b3e-ae59a63d  TATTGAATACACCATATT-ATAA-TTATCC---C--GAGGAAGAA-ATTCAATATACCACTAATTAAAAT---A-G--GA [1600]
44f1962a-fce6-436a-9ea0-ef4a969d  TA-TGAATACACCATATT-ATGA-TTCTGC------GAGGAAGAA-ATTCAATATACCACTAATTAAAAT---A-G--GA [1600]
4544809e-f4f9-4ff0-8142-5a2a2bc9  TA-TGAATACACCATATT-ATAA-TTAT--------ACGCAAGAG-ATTCAATATACCACTAATTGAGAA---T-ATGGA [1600]
9faa5557-cdc1-466c-815f-fb18a15f  TA-TGAATACACCATATT-ATAA-----TTATCCCGAGG-AAGAA-ATTCAATATACCACTAATTAGAAT---AG---GA [1600]
ef1a361c-a80e-43f6-8e1f-aeafa681  TA-TGAATACACCATATT-ATAA-TTATCCCGAG--GA---AGAA-ATTCAATATACCACTAATTAAAAT---A-G--GA [1600]

Cp_PARENT                        C--T-
Ct_PARENT                        C--CA
bdf46f5b-2833-4b26-892f-323dd4c3  C--CA
c40b234c-b999-47c3-93ca-016af61e  C--CA
0d8edcbc-b350-4d00-afbc-fd78a1f6  C--CA
9cb07723-65ed-4c96-ac19-4234ea98  C--CA
ed18c856-4d5b-4622-9fe8-9b5740af  C--CA
```

```
c344612d-f873-4743-818c-65979638  C--CA
baaaf62e-d0ee-4c49-9d83-0313d3df  C--CA
6771788a-69b9-4fd8-b4e0-073881fe  C--CA
a6666a5a-6885-427a-9881-430dad33  C--CA
5be1c0db-7e16-4441-a2ab-f8107a5d  C--CA
99a0bd03-c448-4605-9def-185e9f2d  C--CA
c37ace00-c1f8-46e8-bcdc-c3c33bcb  C--CA
cdc9d647-e2ac-442b-8435-6dd7b82f  C--CA
bc10b467-59cf-4610-94fe-267549f8  C--CA
c8c3f0cb-c211-4877-8f07-1765b113  C--CA
720a0247-e02e-400b-901f-673d7527  C--CA
f3f8defb-2c03-44af-abc8-d94623f7  C--CA
fc2a9b4c-2c5f-43ed-8ea7-f2af0743  C--TA
477953c0-cb9c-400c-bf05-dfadb08e  C--CA
28c5afa3-4ebf-4ffa-a71d-2171913e  C--CA
52a4f968-3394-459b-9ee3-10eab4a7  C--CA
d36d5471-42a7-440b-bd5e-48d77156  C--CA
f7886442-23fe-473a-9ead-a2b8a2a7  C--CA
f4770134-2a4b-449d-a4f6-c99f3383  C--CA
4dd53951-3b2d-42af-82ef-399bb712  C--CA
81ea3bac-8208-46c7-9546-bbd55b7c  C--CA
1e66a4e2-42fd-4fa9-b001-9bb7f263  C--CA
b75fad9a-877e-4f52-968b-9fcbb7a4  C--CA
f512b649-4f14-4eb6-b5a2-b0543441  C--CA
5021e737-de02-42f2-920d-18868672  C--CA
a46adbf5-3ba1-48b8-a762-2403bda7  C--AA
088a7e31-4c6b-4ff7-a19b-ff14531f  C--AA
e82639d1-c451-4263-90a8-2b1ce6a1  C--AA
3cca4221-4eee-44ed-9fb4-ebbac961  C--AA
d7a80796-8741-4878-b6ca-23aa66f7  C--AA
421e3eda-83a0-4ebd-972c-f24667ee  C--AA
e6a39029-4e34-4457-b2a9-0cf6f207  C--AA
a1507163-34c9-4b24-b21e-3b5dfc54  C--CA
51b01061-aaf5-4585-ba72-1a83671b  C--CA
85703d1f-2c69-4541-ab66-acb7b838  C--AA
ed39aa3b-d227-4de1-92bd-5ace4420  C--CA
40e2af31-875d-446e-90a0-6e16a943  C--CA
873c76b4-8a29-4846-98df-cb5abcc9  C--CA
59ef08a5-aa1d-4d15-ba6f-1a925971  C--CA
e1cae4d8-4f39-4401-8d82-5c5edaf3  C--CA
5f47af75-253d-48b0-8b3e-ae59a63d  C--CA
44f1962a-fce6-436a-9ea0-ef4a969d  C--CA
4544809e-f4f9-4ff0-8142-5a2a2bc9  C--CA
9faa5557-cdc1-466c-815f-fb18a15f  C--CA
ef1a361c-a80e-43f6-8e1f-aeafa681  C--CA
```
